# Synergistic Cytotoxicity Between Cold Atmospheric Plasma and Pyrazolopyrimidinones Against Glioblastoma Cells

**DOI:** 10.1101/2021.02.04.429831

**Authors:** Zhonglei He, Clara Charleton, Robert Devine, Mark Kelada, John M. D. Walsh, Gillian E. Conway, Sebnem Gunes, Julie Rose Mae Mondala, Furong Tian, Brijesh Tiwari, Gemma K. Kinsella, Renee Malone, Denis O’Shea, Michael Devereux, Wenxin Wang, Patrick J Cullen, John C. Stephens, James F Curtin

**Author notes:** **Corresponding author** Correspondence to Prof James Curtin and Dr Zhonglei He.

## Abstract

Pyrazolopyrimidinone is a fused nitrogen-containing heterocyclic system, which acts as a core scaffold in many pharmaceutically relevant compounds. Pyrazolopyrimidinones have been demonstrated to be efficient in treating several diseases, including cystic fibrosis, obesity, viral infection and cancer. We have tested the synergistic anti-cancer effects of 15 pyrazolopyrimidinones, synthesised in a two-step process, combined with cold atmospheric plasma (CAP), a novel innovation generating reactive species with other unique chemical and physical effects. We identify two pyrazolopyrimidinones that act as prodrugs and display enhanced reactive-species dependent cytotoxicity when used in combination with cold atmospheric plasma. Synergistic activation was evident for both direct CAP treatment on prodrug loaded tumour cells and indirect CAP treatment of prodrug in media prior to adding to tumour cells. Our results demonstrate the potential of CAP combined with pyrazolopyrimidinones as a programmable cytotoxic therapy against cancer.

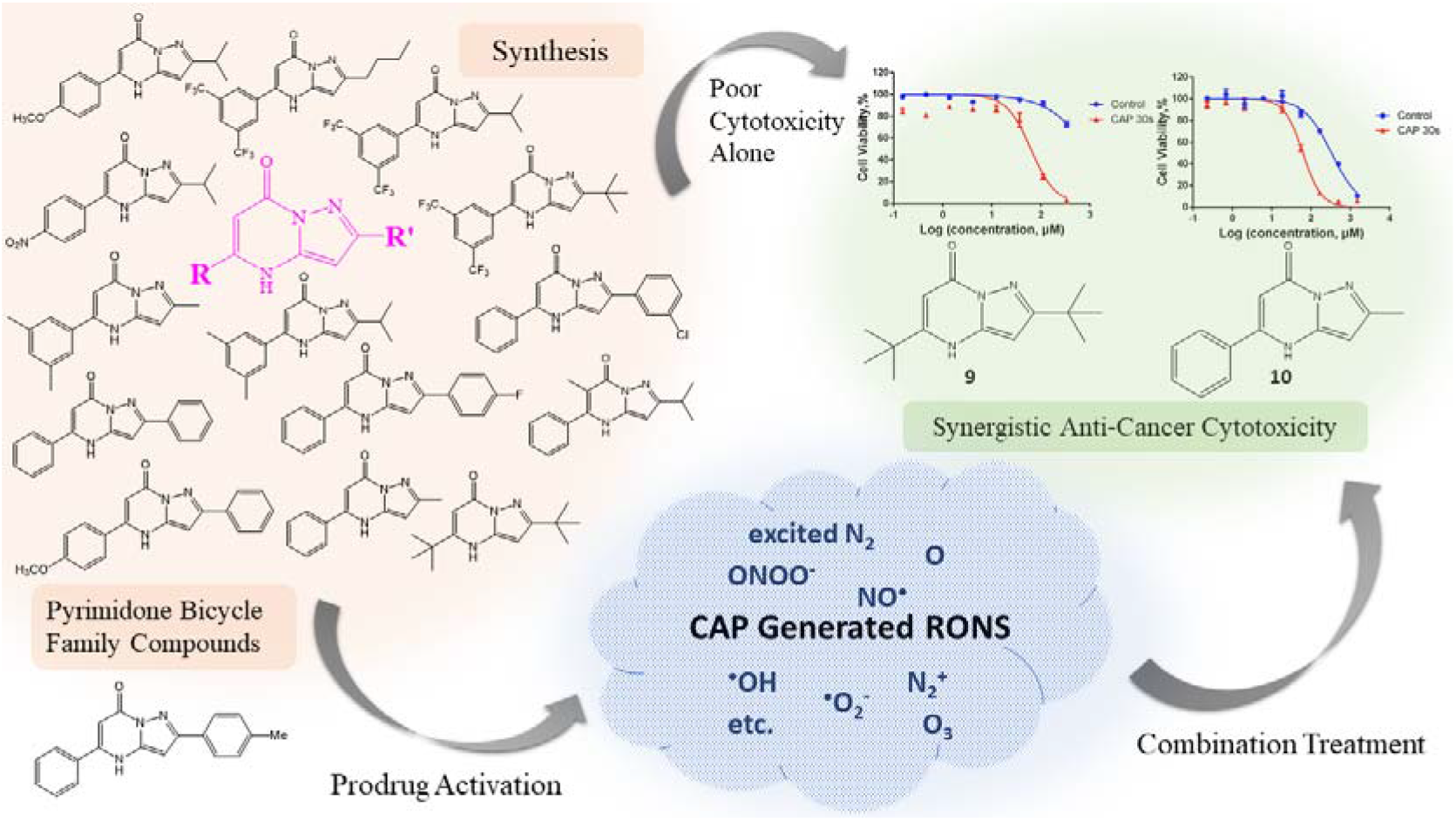

## Introduction

Despite extensive efforts made in the war against cancer over the past decades, many challenges remain due to tumour heterogeneity, varied patient characteristics, treatment efficacy, and toxicity considerations. A major limitation in treating advanced carcinoma is the delivery of sufficient concentrations of a chemotherapeutic agent to the tumour site without causing undue toxicity elsewhere. For this reason, prodrugs that can be locally activated at the site of the tumour with low toxicity elsewhere are of particular interest.

Reactive oxygen species (ROS), as a natural product generated during normal metabolism pathways, play critical roles in various cellular activities and are considered a carcinogenic factor as high levels of ROS are capable of inducing damage and mutation of intercellular DNA and therefore cause malignancy^1^. However, as understanding deepens, ROS was found to be a double-edged sword to cancer cells^2^. Evidence showed that higher ROS levels are generated in cancer cells compared to healthy cells, which is attributed to the higher metabolic activities and more rapid proliferation of transformed cells^2^. Hence, the cellular antioxidant system works under a larger load to protect tumour cells from oxidative stress, a common feature in many types of tumours that can be targeted to develop efficient therapies. Attempts have been made to exploit this difference with healthy tissues, and anti-cancer drugs have been developed that are conditionally activated by high levels of ROS^3^. Cold atmospheric plasma (CAP) can locally induce ROS generation in cells and tissues with a high degree of control of both the amount of ROS generated and location^4^ (Figure 1a,b). Meanwhile, prodrugs, which only have cytotoxicity after being activated by a high ROS level, have fewer side effects and are more specific to cancer cells^5^. Therefore, combining prodrugs with CAP to locally increase tumour ROS and activate cytotoxicity only in tumour tissue may provide a novel promising combination therapy against cancer.

**Figure 1.**
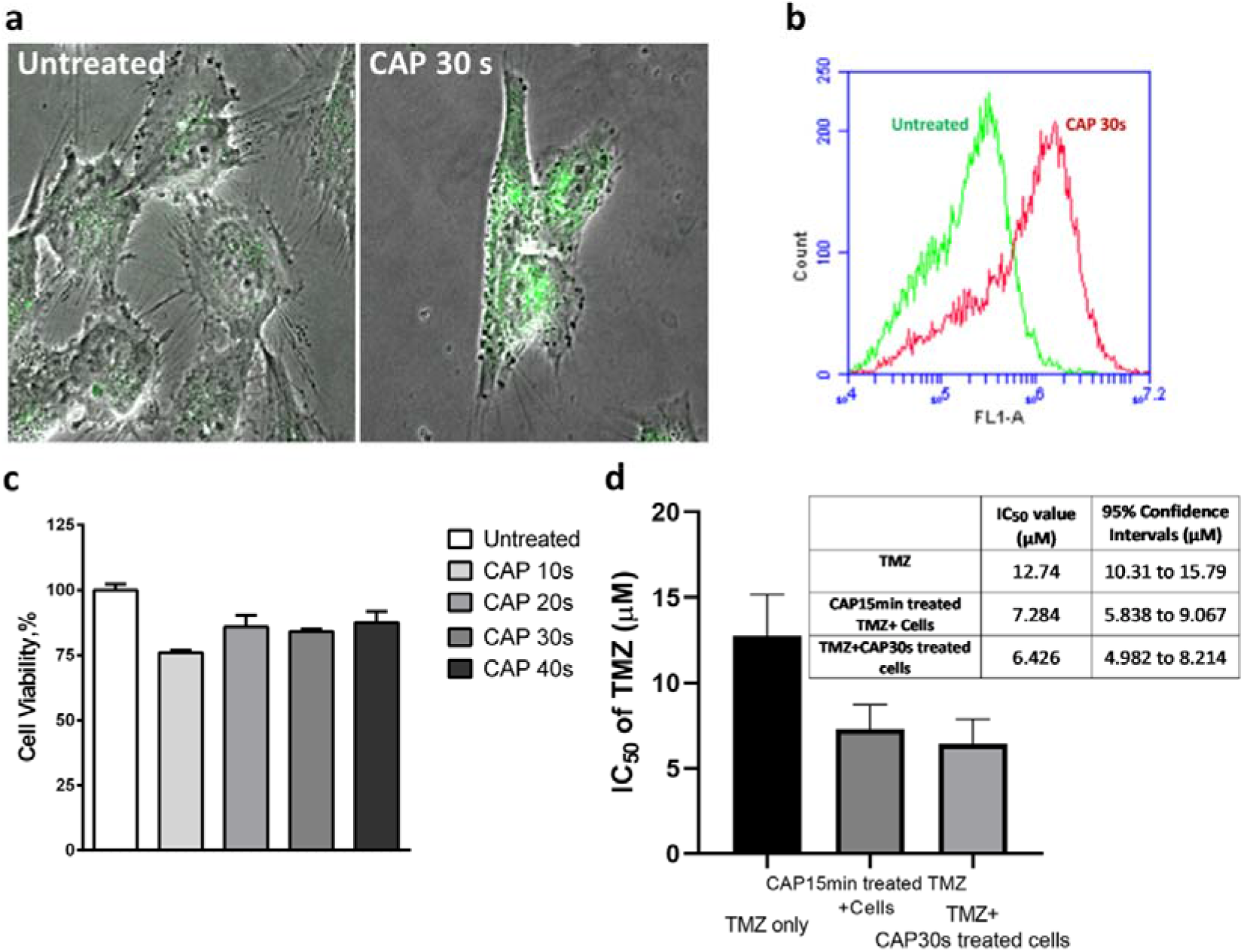
Treatment of U-251 MG cells with reactive species-generating CAP treatment alone or in combination with TMZ. (**a**) Fluorescence levels of oxidised H2DCFDA in untreated and 30s CAP treated cells were observed via confocal microscope. (**b**) Fluorescence level of intracellular oxidised H2DCFDA was measured in U-251 MG cells via Flow cytometry, left curve (green, untreated cells), right curve (red, 30 s CAP treated cells). (**c**) U-251 MG cells were treated with CAP for 10-40 s at 75 kV alone and then incubated for 48 hours before cell viability was assessed. (**d**) CAP untreated or CAP 30s treated U-251 MG cells were incubated with a serial dilution of TMZ only or 15min CAP treated TMZ stock solution for 6 days, Cell viability assays were then carried out, and IC50 values were calculated using GraphPad Prism.

CAP has previously been demonstrated to restore drug sensitivity in cisplatin-resistant ovarian cancer cells^6^, paclitaxel-resistant breast cancer cells^7^, tamoxifen-resistant breast cancer cells^8^, and temozolomide-resistant malignant glioma cells^9^. Therefore, CAP can modify cells and chemotherapeutic agents to enhance the selective cytotoxicity of some chemotherapeutic agents. Here, a systematic study of cytotoxicity was carried out using a panel of novel pyrazolopyrimidinones.

Pyrazolopyrimidinones have been identified with bioactivity for the treatment of cancer^10^, infections^11,12^, obesity^13,14^ and cystic fibrosis^15,16^. It also has been reported that pyrazolopyrimidinones present notable inhibiting effects to the activity or function of several kinases, including the PI3 kinase, glycogen synthase kinase −3 (GSK-3) amongst others. These kinases are involved and can play critical roles in a variety of cellular activities including cell differentiation, motility, cell growth, proliferation, survival and intracellular trafficking^17–20^. We have tested 15 pyrazolopyrimidinones (Figure 2), and screened those candidates for their potential synergistic anti-cancer effect combined with CAP treatment and identified two leading prodrug candidates **9** and **10**. In combination with low dose CAP treatment, which had little or no-toxicity to cancer cells, the leading prodrug candidates’ cytotoxicity was synergistically enhanced more than five times. The synergistic cytotoxicity between 10 and CAP treatment has undergone further investigation, where the ROS generated in culture medium by CAP treatment has been determined to play the primary role in the activation of prodrug **10**.

**Figure 2.**
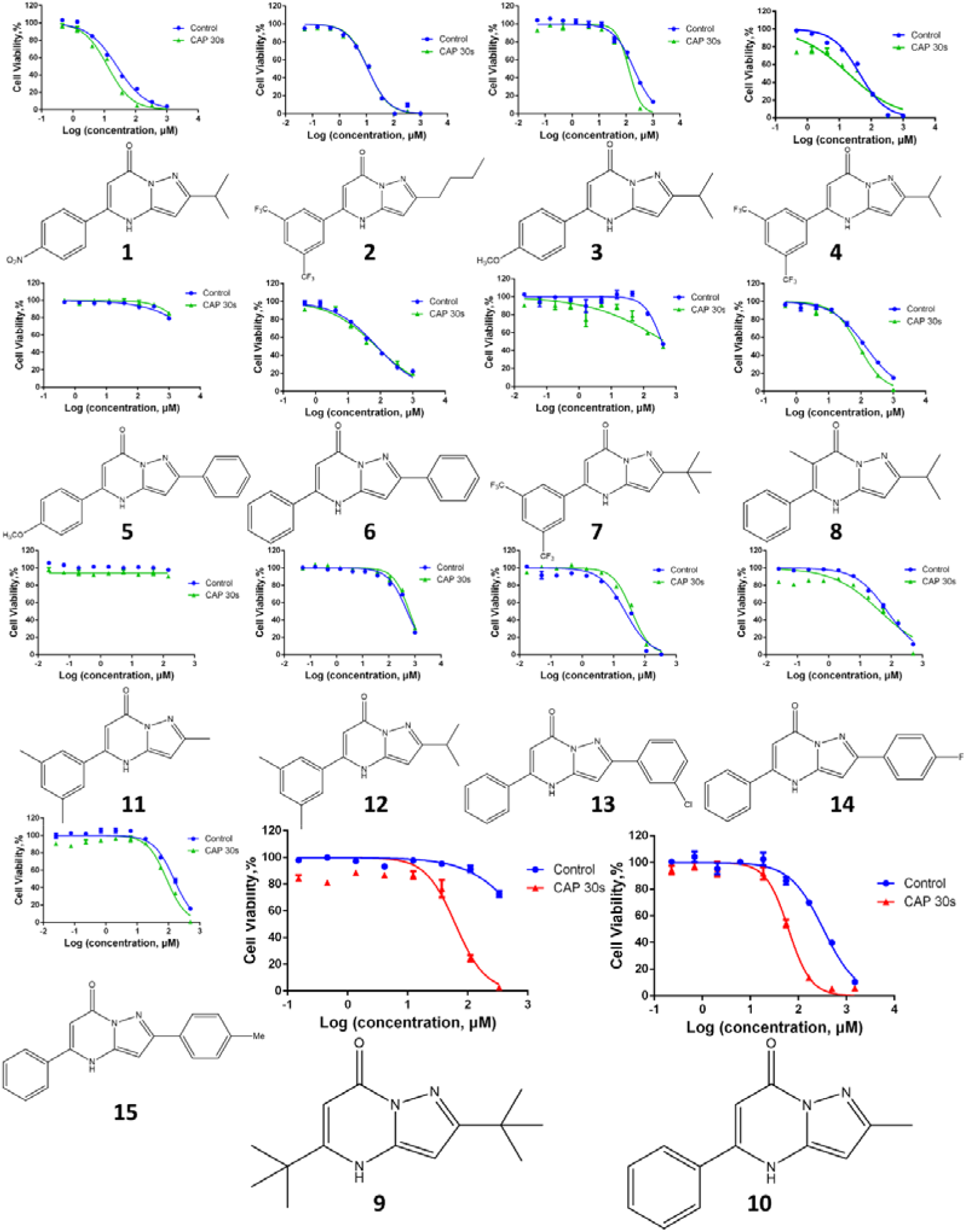
Dose response curves of Pyrazolopyrimidinones. The stock solution of pyrazolopyrimidinones **1-15** were prepared in 100% DMSO, which was then serially diluted in culture medium without pyruvate to different concentrations as indicated. U-251 MG cells were treated with **1-15** in combination with 30 s of CAP treatment as indicated in Method section. Cell viability assays were then carried out following incubation with 1-15 culture medium solution for 48 h.

## Methods

### Chemistry General Information

All reagents for synthesis were bought commercially and used without further purification. The 3-amino-5-isopropylpyrazole was purchased from Fluorochem and used as received. Reactions were monitored with thin layer chromatography (TLC) on Merck Silica Gel F_254_ plates. NMR spectra were recorded using Bruker Ascend 500 spectrometer at 293K. All chemical shifts were referenced relative to the relevant deuterated solvent residual peaks or TMS. Assignments of the NMR spectra were deduced using ^1^H NMR and ^13^C NMR, along with 2D experiments (COSY, HSQC and HMBC). The following abbreviations were used to explain the observed multiplicities; s (singlet), d (doublet), t (triplet), q (quartet), hept (heptet), m (multiplet), bs (broad singlet), pt (pseudo triplet). Flash chromatography was performed with Merck Silica Gel 60. Microwave reactions were carried out using a CEM Discover Microwave Synthesizer with a vertically focused floor mounted infrared temperature sensor, external to the microwave tube. The 10 mL reaction vessels used were supplied from CEM and were made of borosilicate glass. High resolution mass spectrometry (HRMS) was performed on an Agilent-LC 1200 Series coupled to a 6210 or 6530 Agilent Time-Of-Flight (TOF) mass spectrometer equipped with an electrospray source in both positive and negative (ESI+/−) modes. Infrared spectra were obtained as KBr disks in the region 4000–400 cm^−1^ on a Perkin Elmer Spectrum 100 FT-IR spectrophotometer. Further experimental details and associated spectra can be found in the Supporting Information.

### Synthesis of pyrazolopyrimidinones

Pyrazolopyrimidinones **1-4, 8, 9, 11, 12** were prepared as described below, NMR spectra can be found in the Supporting Information. Pyrazolopyrimidinones **5-7, 10, 13, 14, 15** were prepared as per the literature procedure reported by Kelada et al. with full experimental details and NMR spectra available in the Supporting Information^15^. Full experimental details and spectra for the synthesis of the required amino pyrazoles are also found in the Supporting Information.

### Synthesis of 2-isopropyl-5-(4-nitrophenyl)pyrazolo[1,5-a]pyrimidin-7(4H)-one (1)

Ethyl 3-(4-nitrophenyl)-3-oxopropanoate (209 mg, 0.88 mmol) was heated at reflux with 3-amino-5-isopropylpyrazole (100 mg, 0.80 mmol) in acetic acid (6 mL) for 6 hours. The reaction was cooled to room temperature and the solvent removed under reduced pressure. The residue was dissolved in tetrahydrofuran (THF) (~5 mL) and petroleum ether (40-60 fraction, 5 mL) added. The resulting precipitate was filtered and washed with a minimum of 1:1 mixture of THF: petroleum ether (40-60) to give 2-isopropyl-5-(4-nitrophenyl)pyrazolo[1,5-a]pyrimidin-7(4H)-one, 83 mg (58 %). ^1^H NMR: (300 MHz, CD_3_OD) δ 8.44 (d, *J* = 8.6 Hz, 2H), 8.05 (d, *J* = 8.6 Hz, 2H), 6.19 (s, 1H), 6.16 (s, 1H), 3.12 (m, 1H), 1.37 (s, 3H), 1.35 (s, 3H); ^13^C NMR: (75 MHz, CD_3_OD) δ 164.3, 157.5, 149.5, 148.8, 142.4, 138.7, 128.3, 123.8, 95.0, 86.6, 28.4, 21.4; HRMS: calcd for C_15_ H_15_ N_4_ O_3_ m/z: [M + H]^+^, 299.1139, found: 299.1149 [Diff (ppm) = 3.46].

### Synthesis of 5-(3,5-bis(trifluoromethyl)phenyl)-2-butylpyrazolo[1,5-a]pyrimidin-7(4H)-one (2)

Ethyl 3-(3,5-bis(trifluoromethyl)phenyl)-3-oxopropanoate (150 mg, 0.45 mmol) was heated at reflux with 3-butyl-1H-pyrazol-5-amine (63 mg, 0.45 mmol) in acetic acid (2 mL) for 6 hours. The reaction was cooled to room temperature and the solvent removed under reduced pressure. The residue was dissolved in THF (~5 mL) and petroleum ether (40-60 fraction, 5 mL) added. The resulting precipitate was filtered and washed with a minimum of 1:1 mixture of THF: petroleum ether (40-60) to give 5-(3,5-bis(trifluoromethyl)phenyl)-2-butylpyrazolo[1,5-a]pyrimidin-7(4H)-one, 43 mg (23 %). ^1^H NMR (300 MHz, CD_3_OD) δ 8.44 (s, 2H), 8.24 (s, 1H), 8.32 (s, 1H), 6.22 (s, 1H), 6.18 (s, 1H), 2.78 (t, *J* = 7.6 Hz, 2H), 1.74 (m, *J* = 7.6 Hz, 2H), 1.44 (m, *J* = 7.6 Hz, 2H), 0.98 (t, *J* = 7.6 Hz, 2H); ^13^C NMR (75 MHz, CD_3_OD) δ 158.7, 157.4, 147.9, 142.4, 135.3, 132.3 (q, *J*_CF_ = 34.1 Hz), 127.8 (m), 124.3 (m), 123.1 (q, *J*_CF_ = 272 Hz), 95.1, 88.6, 31.2, 28.0, 22.0, 12.8.

### Synthesis of 2-isopropyl-5-(4-methoxyphenyl)pyrazolo[1,5-a]pyrimidin-7(4H)-one (3)

A microwave tube was charged with ethyl 4-methoxybenzoyl acetate (0.17 mL, 0.9 mmol), 3-amino-5-isopropylpyrazole (110 mg, 0.9 mmol), and methanol (2 mL) and subjected to microwave irradiation (100 W, 120 °C) for 1 hour. At this point, three drops of acetic acid were added, and the reaction mixture subjected to microwave irradiation (100 W, 120 °C) for a further 2 hours. The resultant precipitate was removed by filtration and washed with a minimum amount of cold methanol, giving 42 mg of 2-isopropyl-5-(4-methoxyphenyl)pyrazolo[1,5-a]pyrimidin-7(4H)-one. The filtrate was evaporated to dryness and the resulting residue recrystallised from methanol to give a further 85 mg of 2-isopropyl-5-(4-methoxyphenyl)pyrazolo[1,5-a]pyrimidin-7(4H)-one. The combined yield was 127 mg (50 %). ^1^H NMR: (300 MHz, d_6_-DMSO) δ 12.21 (s, 1H, NH), 7.79 (d, *J* = 8.9 Hz, 2H), 7.13 (d, *J* = 8.9 Hz, 2H), 6.03 (s, 1H), 5.95 (s, 1H), 3.85 (s, 3H), 3.01 (m, 1H), 1.28 (s, 3H), 1.26 (s, 3H); ^13^C NMR: (75 MHz, d_6_-DMSO) δ 161.8, 161.4, 156.3, 142.1, 128.7, 114.4, 92.5, 86.4, 55.5, 27.9, 22.3; HRMS: calcd for C_16_H_18_N_3_O m/z: [M + H]^+^, 284.1394, found: 284.1388 [Diff (ppm) = 2.05].

### Synthesis of 5-(3,5-bis(trifluoromethyl)phenyl)-2-isopropylpyrazolo[1,5-a]pyrimidin-7(4H)-one (4)

A microwave tube was charged with 3-isopropyl-1H-pyrazol-5-amine (113 mg, 0.9 mmol), ethyl 3-(3,5-bis(trifluoromethyl)phenyl)-3-oxopropanoate (295 mg, 0.9 mmol), acetic acid (28.6 μL, 0.5 mmol), MeOH (1 mL), and subjected to MW irradiation (100 W, 150 °C) for 2 h. The resulting mixture was concentrated in vacuo and purified by trituration with cold ethyl acetate to give 5-(3,5-bis(trifluoromethyl)phenyl)-2-isopropylpyrazolo[1,5-a]pyrimidin-7(4H)-one, 140 mg (40 %). ^1^H NMR (500 MHz, d_6_-DMSO) δ 12.65 (bs, 1H), 8.51 (s, 2H), 8.32 (s, 1H), 6.30 (s, 1H), 6.10 (s, 1H), 3.05 – 2.95 (m, *J* = 13.5, 6.7 Hz, 1H), 1.25 (d, *J* = 6.8 Hz, 6H); ^13^C NMR (126 MHz, d_6_-DMSO) δ 162.4, 156.1, 146.3, 142.1, 135.0, 131.1 (q, *J*_CF_ = 33.3 Hz), 128.5, 124.4, 123.2 (q, *J*_CF_ = 273.4 Hz), 95.5, 86.9, 28.0, 22.3; Rf 0.5 (1:9 v/v MeOH:EtOAc); HRMS: calcd for C_17_H_14_F_6_N_3_O m/z, [M + H]+, 390.1036, found: 390.1051, [Diff(ppm) = 2.79]; IR (KBr): 3423 (N-H), 2972 (C-H), 1670 (C=O), 1617 (C=C), 1577 (N-H), 1481 (C=C), 1279 (C-F), 1139 (C-N) cm^-1^.

### Synthesis of 2-isopropyl-6-methyl-5-phenylpyrazolo[1,5-a]pyrimidin-7(4H)-one (8)

Ethyl 2-methyl-3-oxo-3-phenylpropanoate (125 mg, 0.61 mmol) was heated at reflux with 3-amino-5-isosubstitutedpyrazole (69 mg, 0.55 mmol) in acetic acid (2 mL) for 15 hours. The reaction was allowed to cool to room temperature and the solvent removed under reduced pressure. The residue was triturated with ethyl acetate to yield a solid. The solid was isolated by filtration, washed with ethyl acetate, and dried in a drying pistol to give 2-isopropyl-6-methyl-5-phenylpyrazolo[1,5-a]pyrimidin-7(4H)-one, 54 mg (37 %). ^1^H NMR (500 MHz, MeOD) δ 7.70 – 7.45 (m, 5H), 5.98 (s, 1H), 3.08 (hept, *J* = 7.0 Hz, 1H), 2.00 (s, 3H), 1.32 (d, *J* = 7.0 Hz, 6H). ^13^C NMR (126 MHz, MeOD) δ 164.0, 163.9, 158.7, 148.0, 141.6, 141.5, 133.8, 129.8, 128.5, 128.4, 101.7, 101.7, 84.7, 28.4, 28.4, 21.6, 10.7; HRMS: calcd for C_16_H_18_N_3_O m/z, [M + H]+, 268.1444, found: 268.1452, [Diff(ppm) = 2.95].

### Synthesis of 2,5-di-tert-butylpyrazolo[1,5-a]pyrimidin-7(4H)-one (9)

The amino pyrazole 5-amino-3-(tert-butyl)-1H-pyrazole (100 mg, 0.71 mmol) was dissolved in acetic acid (10 mL), and ethyl 4,4-dimethyl-3-oxopentanoate (116 μL, 0.65 mmol) added. The reaction mixture was heated at reflux overnight. After cooling to rt, the acetic acid was removed under reduced pressure and trituration with ethyl acetate (5 mL) gave 2,5-di-tert-butylpyrazolo[1,5-a]pyrimidin-7(4H)-one, 42 mg (23%). ^1^H NMR: (300 MHz, CD_3_OD) δ 6.08 (s, 1H, H5), 5.74 (s, 1H, H2), 1.38 (s, 18H, (CH_3_)_3_ × 2); ^13^C NMR: (125 MHz, CD_3_OD) δ 166.7, 161.6, 158.5, 142.2, 91.3, 85.4, 34.9, 32.4, 29.2, 27.6; R_f_: 0.6 (1:9 v/v, MeOH:DCM); HRMS: calcd for C_14_H_22_N_3_O m/z, [M + H]^+^, 248.1757, found: 248.1768, [Diff(ppm) = 4.45]; IR (KBr): 3090 (N-H), 2965 (C-H), 1670 (C=O), 1614 (C=C), 1570 (N-H), 1481 (C=C), 1240 (C-N) cm^-1^.

### Synthesis of 5-(3,5-dimethylphenyl)-2-methylpyrazolo[1,5-a]pyrimidin-7(4H)-one (11)

Ethyl 3-(3,5-dimethylphenyl)-3-oxopropanoate (0.56 mL, 3.0 mmol) was added to a solution of 3-amino-5-methylpyrazole (132 mg, 1.0 mmol) dissolved in acetic acid (5 mL) and heated at reflux for 21 hours. Volatiles were removed under reduced pressure. The product was purified by column chromatography (100 % DCM) to give 5-(3,5-dimethylphenyl)-2-methylpyrazolo[1,5-a]pyrimidin-7(4H)-one, 204 mg, (59%). ^1^H NMR: (500 MHz, CD_3_OD) δ 7.42 (s, 2H), 7.26 (s, 1H), 6.13 (s, 1H), 6.05 (s, 1H), 2.53 – 2.34 (m, 9H); ^13^C NMR: (126 MHz, CD_3_OD) δ 138.9, 132.3, 124.5, 19.9; HRMS: calcd for C_13_H_16_N_3_O m/z, [M + H]+, 254.1293’ found: 253.1207, [Diff (ppm) = 3.3]; Rf: 0.11 (100% DCM); IR (KBr): 3251 (N-H), 2917 (C-H), 1665 (C=O), 1618 (N-H), 1597 (Ar C-C) cm^-1^.

### Synthesis of 5-(3,5-dimethylphenyl)-2-isopropylpyrazolo[1,5-a]pyrimidin-7(4H)-one (12)

Ethyl 3-(3,5-dimethylphenyl)-3-oxopropanoate (28 mL, 1.4 mmol) was added to a solution of 3-isopropyl-1*H*-pyrazol-5-amine (176 mg, 1.4 mmol) dissolved in acetic acid (5 mL) and refluxed for 5.5 hrs. Volatiles were removed under reduced pressure. The product was purified by trituration using ethyl acetate (10 mL) and give 5-(3,5-dimethylphenyl)-2-isopropylpyrazolo[1,5-a]pyrimidin-7(4H)-one, 165.4 mg (42%). ^1^H NMR: (500 MHz, CDCl_3_) δ 9.51 (bs, 1H), 7.25 (s, 2H), 7.13 (s, 1H), 6.00 (m, 2H), 3.10 (septet, 1H), 2.34 (s, 6H), 1.28 (d, *J* = 7.0 Hz, 6H); ^13^C NMR: (126 MHz, CDCl_3_) δ 157.8, 141.6, 139.1, 132.8, 132.7, 124.5, 94.6, 86.7, 28.6, 22.5, 21.3; HRMS: calcd for C_17_H_20_N_3_O m/z: [M + H]^+^, 281.1528, found: 281.1528 [Diff (ppm) = 0.06]; IR (KBr): 3199 (N-H), 3081 (C-H), 1651 (C=O), 1595 (N-H), 1291 (Ar C-N) cm^-1^.

### U-251 MG Cell Culture

U-251 MG (formerly known as U373MG) (ECACC 09063001), human brain glioblastoma cancer cells (Obtained from Dr Michael Carty, Trinity College Dublin) were cultured in DMEM-high glucose medium supplemented with 10% FBS and maintained in a 37 ℃ incubator within a humidified 5% (v/v) CO_2_ atmosphere. For prodrug treatment, DMEM-high glucose medium without pyruvate was used to make up culture medium to avoid the antioxidant effects of pyruvate ^21^.

### H_2_DCFDA Assay

H_2_DCFDA was used to detect ROS induced by CAP treatment. U-251 MG cells were seeded into 35 × 10 mm tissue culture dishes (Sarstedt) at a density of 2×10^5^ cells/ml, or 35 mm glass-bottom dishes (Greiner Bio-One) at a density of 1×10^5^ cells/ml and incubated overnight as above to allow adherence. Prior to treatment with CAP at 75 kV for 30 s, cells were first incubated with 25 μM H_2_DCFDA (Thermo Fisher Scientific) in serum-free medium for 30 min at 37 ℃, then washed with PBS twice, culture medium once. Afterwards, cells were observed under a Zeiss LSM 510 confocal laser scanning microscope (excitation 488 nm, emission 505-530 nm), or were collected for measurement with flow cytometry.

### CAP Configuration and Prodrug Treatment

An experimental atmospheric dielectric barrier discharge (DBD) plasma reactor, DIT-120, was used which has been described and characterised in detail elsewhere ^22,23^. Unless otherwise stated, all U-251 MG cells were treated within 96-well plates. During CAP treatment, containers were placed in between two electrodes, at a voltage level of 75 kV for 10-40 s. The culture medium was removed before CAP treatment then replaced with fresh culture medium immediately afterwards.

An optimised prodrug treatment protocol was developed and applied, as described below. U-251 MG cells were plated into 96-well plates (Sarstedt) at a density of 1×10^4^ cells/well (100 μl standard culture medium per well) and were incubated overnight for proper adherence. Unless otherwise stated, the plating map was 6×10 wells, for ten different concentrations, negative and positive control groups (5 replicates for each group). All prodrugs were dissolved and prepared in 100% DMSO for the stock solution, which was then serially diluted in culture medium to different concentrations as indicated. 10 μl of prodrug non-pyruvate culture medium solution was then added to each well after removing the previous medium, and the remainder was added into empty wells in the plate. The plate then was treated with CAP at 75 kV for 10-40 s as indicated. 90 μl of CAP-treated prodrug culture medium solution was then added to corresponding wells with cells to final volume 100 μl immediately after CAP treatment. Cell viability assays were then carried out following incubation for the indicated time. Temozolomide (TMZ) (Merck) was dissolved in DMSO as stock (50 mM) and then subsequently diluted in culture medium to concentrations as indicated. U-251 MG cells were plated into 96-well plates at a density of 2×10^3^ cells/well and were incubated overnight to ensure proper adherence. The combination treatment of TMZ and 30 s CAP was performed with the same method as above. Meanwhile, the TMZ stock was first treated with CAP for 15 min then diluted in culture medium to indicated concentrations and incubated with untreated cells, for six days, as indicated.

Indirect CAP treatment was used to investigate the effects of CAP on pro-drugs alone to determine the mechanism of synergistic cytotoxicity to U-251 MG cells. Pro-drugs were treated with CAP at 75 kV for 10 s to 10 min in DMSO stock solution or culture medium solution as indicated. The CAP-treated prodrug solution was then added to U-251 MG cells in 96-well plates to incubate with U-251 MG cells. For CAP-activated medium, the culture medium was treated with CAP at 75 kV for 30 sec in 96-well plate and incubated overnight to remove short-lived reactive species. The prodrug stock solution was then serially diluted in the overnight storage CAP-activated medium and incubated with U-251 MG cells. For CAP-activated cells, the U-251 MG cells overlaid with 10 μl culture medium were treated with CAP for 30 sec in 96-well plates, while remainder of culture medium was added into empty wells in the plate. Then the CAP-treated culture medium was added to each well to 100 μl. Those CAP-activated cells were then incubated in the CAP-activated medium for 0-5 hours as indicated. Following incubation, the medium was replaced with fresh medium containing corresponding concentrations of prodrugs.

### Cytotoxicity and Inhibitor Assays

Cell viability was analysed using the Alamar blue assay (Thermo Fisher Scientific). 48 hours after CAP treatment, the cells were rinsed once with PBS, incubated for 3 h at 37 ℃ with a 10% Alamar blue/90% culture medium solution. The fluorescence was then measured (excitation, 530 nm; emission, 595 nm) by a Victor 3V 1420 microplate reader (Perkin Elmer). 4 M N-Acetyl Cysteine (NAC) stock solution was prepared in water. 4 M NAC solution was then diluted in culture medium to a final concentration of 4 mM before the addition of prodrugs and CAP treatment. The CAP treated culture medium containing prodrugs and 4 mM NAC was then added into 96-well plates to incubate with U-251 MG cells.

### Statistical Analysis

Triplicate independent tests were carried out for each data point unless indicated otherwise. Error bars of all figures are presented using the standard error of the mean (S.E.M). Prism 7 (GraphPad Software, San Diego, CA, USA) was used to carry out curve fitting and statistical analysis. The isobologram and synergistic analysis were carried out using CompuSyn software (ComboSyn, Inc., Paramus, NJ, USA, www.combosyn.com)^24^. Two-tailed P values were used where alpha = 0.05. The significance between data points was verified using one-way ANOVA and two-way ANOVA with Tukey’s multiple comparison post-test, as indicated in figures (*P<0.05, **P<0.01, ***P<0.001, ****p<0.0001).

## Results

### ROS generation by low dose CAP treatment

CAP has been well known for inducing the generation of ROS ^23,25,26^. In addition to flow cytometry, we have used a confocal microscope and ROS indicator H_2_DCFDA to demonstrate the ROS generated by 30 s of CAP treatment (Figure 1a, b). Meanwhile, 30 s of CAP treatment presented relatively low cytotoxicity to U-251 MG cell lines, with only decrease around 20% in cell viability as previously described^23,25^. As seen in Figure 1c, 10-40 s of CAP treatment decreased around 10-20% of cell viability when U-251 cells were treated and incubated in culture medium without pyruvate. Temozolomide (TMZ), the standard chemotherapy for glioblastoma, has been tested in combination with 30 s and 15 min of CAP treatment on cells or TMZ stock solution separately. As seen in Figure 1d, there is a marginal decrease of IC_50_ values of TMZ when U-251 MG cells were treated with both TMZ and 30s of CAP treatment (direct treatment), which is in agreement with previously published studies demonstrating that CAP possesses an ability to restore the sensitivity of U-251 MG cells to TMZ^9,23^. We also found that 15 min of CAP treatment on TMZ stock solution did not induce a significant degrade TMZ cytotoxicity, rather, we observed a similar marginal decrease in IC_50_ values (Figure 1d and Support Information Figure S35). In the following section, we carried out the cytotoxicity study for a series of synthesised prodrug candidates combined with CAP treatment and have identified two leading candidates that possess remarkable synergistic anti-cancer effects compared to TMZ.

### Pyrimidone Bicycle Family Compounds

A series of pyrazolopyrimidinones^15^ were tested using the U-251 MG cell line. As seen in Figure 2, most pyrazolopyrimidinones had no synergistic cytotoxicity in combination with CAP treatment, the dose-response curves of CAP-treated groups presented high comparability and similar trends with untreated groups. Two compounds from the pyrazolopyrimidinone family, **9** and **10**, presented significant synergistic cytotoxicity with 30 s of CAP treatment at 75 kV (Figure 2). The CAP treatment induced a pronounced descent of dose-response curves with the increasing of concentrations of prodrugs. As seen in Table 1, CAP treatment induced significant decreases of IC_50_ values of **9** (from 940.6 μM to 62.41 μM) and **10** (from 292.8 μM to 56.79 μM) while presenting low cytotoxicity by themselves. Therefore, among all 15 candidates, **9** and **10** were determined as the leading candidates of CAP-activated pro-drugs.

**Table 1.**
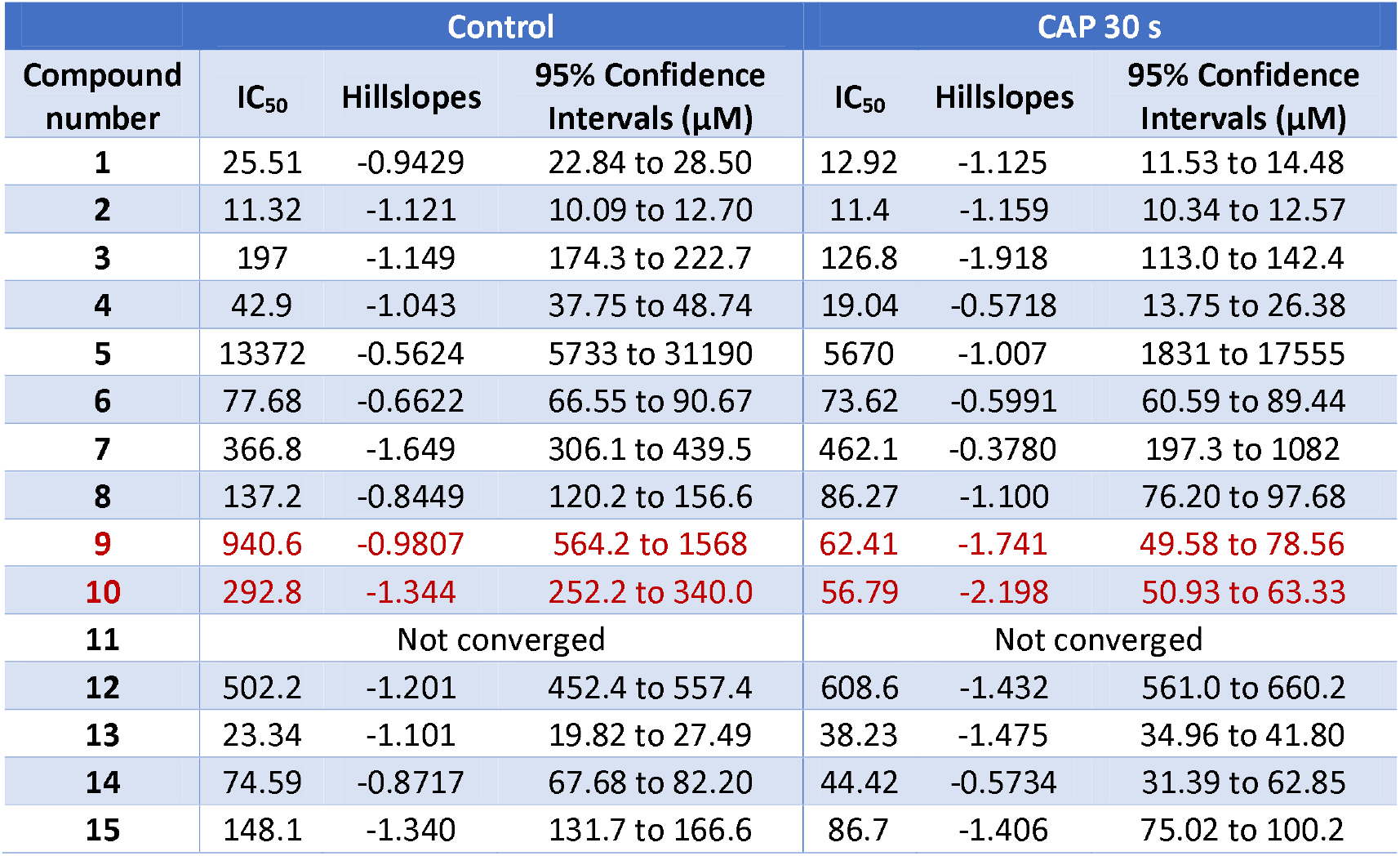
IC_50_ values and 95% confidence intervals of pyrazolopyrimidinones alone or in combination with 30 s of CAP treatment against U-251 MG cells.

| Compound number | Control |  |  | CAP 30 s |  |  |
| --- | --- | --- | --- | --- | --- | --- |
|  | IC<sub>50</sub> | Hillslopes | 95% Confidence Intervals (μM) | IC<sub>50</sub> | Hillslopes | 95% Confidence Intervals (μM) |
| 1 | 25.51 | -0.9429 | 22.84 to 28.50 | 12.92 | -1.125 | 11.53 to 14.48 |
| 2 | 11.32 | -1.121 | 10.09 to 12.70 | 11.4 | -1.159 | 10.34 to 12.57 |
| 3 | 197 | -1.149 | 174.3 to 222.7 | 126.8 | -1.918 | 113.0 to 142.4 |
| 4 | 42.9 | -1.043 | 37.75 to 48.74 | 19.04 | -0.5718 | 13.75 to 26.38 |
| 5 | 13372 | -0.5624 | 5733 to 31190 | 5670 | -1.007 | 1831 to 17555 |
| 6 | 77.68 | -0.6622 | 66.55 to 90.67 | 73.62 | -0.5991 | 60.59 to 89.44 |
| 7 | 366.8 | -1.649 | 306.1 to 439.5 | 462.1 | -0.3780 | 197.3 to 1082 |
| 8 | 137.2 | -0.8449 | 120.2 to 156.6 | 86.27 | -1.100 | 76.20 to 97.68 |
| 9 | 940.6 | -0.9807 | 564.2 to 1568 | 62.41 | -1.741 | 49.58 to 78.56 |
| 10 | 292.8 | -1.344 | 252.2 to 340.0 | 56.79 | -2.198 | 50.93 to 63.33 |
| 11 | Not converged |  |  | Not converged |  |  |
| 12 | 502.2 | -1.201 | 452.4 to 557.4 | 608.6 | -1.432 | 561.0 to 660.2 |
| 13 | 23.34 | -1.101 | 19.82 to 27.49 | 38.23 | -1.475 | 34.96 to 41.80 |
| 14 | 74.59 | -0.8717 | 67.68 to 82.20 | 44.42 | -0.5734 | 31.39 to 62.85 |
| 15 | 148.1 | -1.340 | 131.7 to 166.6 | 86.7 | -1.406 | 75.02 to 100.2 |

### Investigate the Mechanism behind the Synergistic Cytotoxicity Between Leading Pro-Drug Candidate **10** and CAP Treatment

We selected **10** to further investigate the synergistic anti-cancer effects combined with CAP treatment. To determine the mechanism of CAP activation of **10**, it was first dissolved in 100% DMSO to 200 mM and then treated with 30 s and 10 min with CAP. Afterwards, CAP-treatment **10** DMSO solution was diluted in culture medium to 2000 μM (DMSO final concentration 1%) before carrying out a serial dilution on U-251 MG cells. Cytotoxicity was measured using Alamar blue, 48 hours post incubation with **10**. As seen from Figure 3a and Table 2, no significant increase in cytotoxicity was observed whether the **10** DMSO solution was treated with CAP or not. ROS generation is absent in 100% DMSO solvent, whereas the generation of reactive species relies on the presence of reactants in treated liquid^27^ and DMSO may function as a scavenger of OH radicals^28^. Therefore, we hypothesised that ROS generation might be essential for synergistic activation of prodrugs.

**Table 2.** IC_50_ values of 10 treated by CAP in DMSO solution.

| | $IC_{50}$ | 95% Confidence Intervals | Hill Slopes |
| --- | --- | --- | --- |
| **10** Control<br>(2<sup>nd</sup> repeat) | 240.5 μM | 170.8 μM to 338.8 μM | -1.408 |
| CAP 30 s treated **10** | 215.3 μM | 169.4 μM to 273.5 μM | -1.139 |
| CAP 10 min treated **10** | 246.0 μM | 235.4 μM to 257.1 μM | -3.385 |

**Figure 3.**
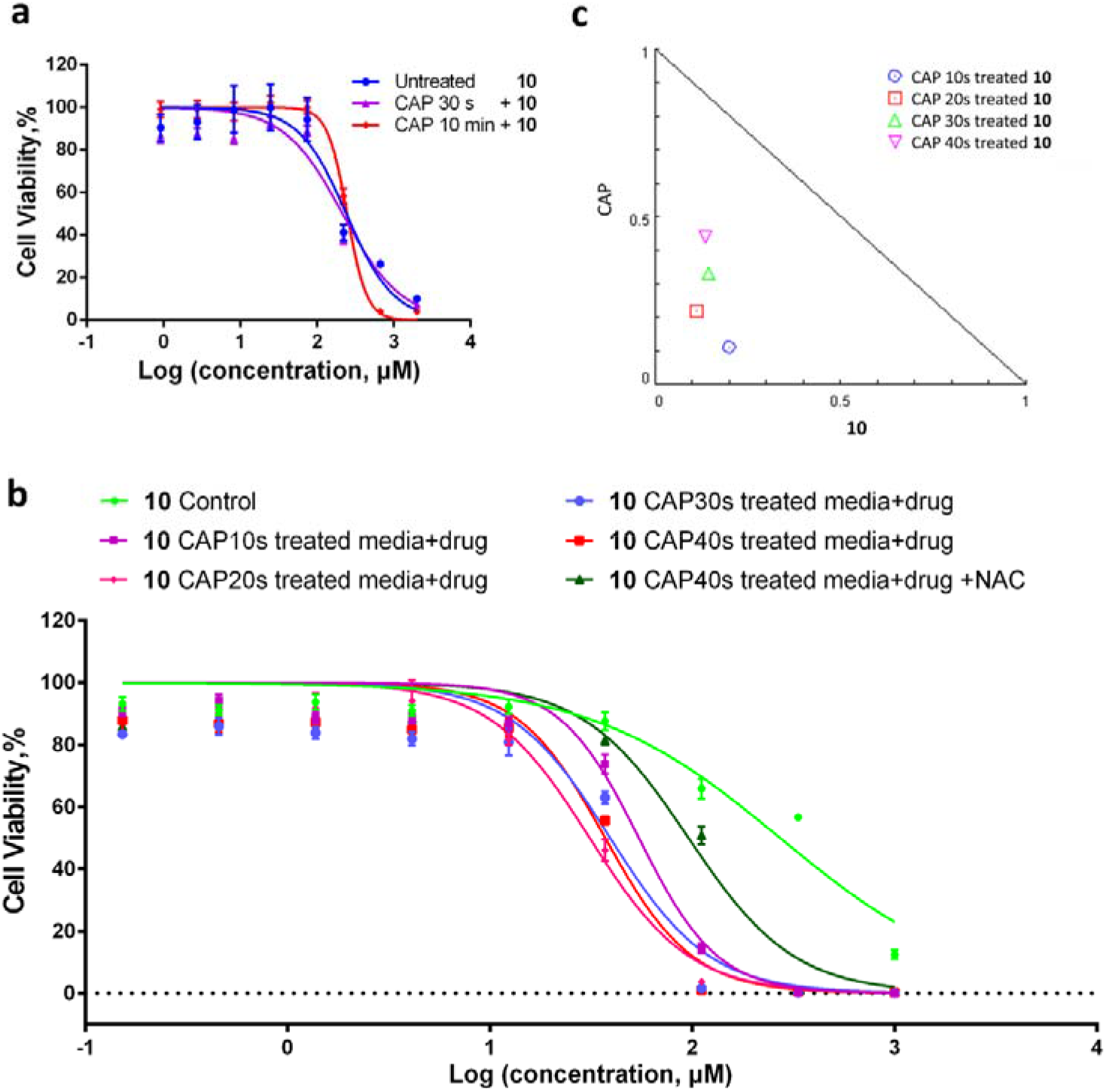
Dose responses of U-251 MG cells to 10 treated by CAP in culture medium solution or DMSO stock solution. (a) **10** was prepared in DMSO stock solution and treated with CAP for 30 s and 10 min, serially diluted into fresh culture medium and incubated with U-251 MG cells for 48 hours and compared with control groups. (b) **10** was prepared in the culture medium, treated with CAP for 0-40 s with or without NAC and then incubated with U-251 MG cells in 96-well plates for 48 hours before cell viability was assessed. (c) Isobologram analysis of the combinational effect of **10** and CAP. The single doses CAP on the y-axis and **10** on the x-axis were used to draw the additivity line. Four combination points were indicated in the isobologram (CAP 10-40s and corresponding IC_50_ values of CAP-treated **10**). The localisation of combined 10 and CAP at different time exposures can be translated to synergism CI<1, additivity CI=1 or antagonism CI>1.

We next diluted **10** in culture medium and treated with CAP in 10 s increments from 0 s to 40 s. The CAP-treated **10** culture medium solution was subsequently incubated with U-251 MG cells for 48 h before the Alamar Blue assay. As seen in Figure 1c, 3b and Table 3, CAP 10-40 s treated **10** culture medium solution induced a pronounced decrease of cell viability with concentrations higher than 10 μM compared to cells treated with media not exposed to CAP. CAP treatment of media alone had low cytotoxicity (~20% decrease in cell viability). Meanwhile, NAC was applied as an antioxidant to investigate the mechanism further. The 4 M NAC solution was diluted in the culture medium until final concentration 4 mM before adding **10** and 40 s CAP treatment. As seen in Figure 3b and Table 3, the NAC treated group presented much higher cell viability and a higher IC_50_ value compared with the CAP treated groups, which demonstrated that ROS generated by CAP treatment played an important role in the activation of anti-cancer effects of **10**. The normalised isobologram was also analysed and presented in Figure 3c, which further confirms the synergistic effects between CAP treatment and **10** treated in the culture medium. Using CompuSyn software, a synergistic analysis was carried out, and the combination index (CI) values have been calculated. The CI values of **10** with 10 s CAP were 0.31099, 0.33520 with 20 s, 0.47561 with 30 s, and 0.57796 with 40 s, which were all less than 1.00 and confirmed the significant synergistic cytotoxicity between CAP treatment and **10** (Figure 3c). **11**, which has a highly similar structure, only two more methyl groups on the benzene ring, compared to **10**, has also been further tested for comparison. Interestingly, as seen in Support Information Figure S36a, **11** showed no synergistic cytotoxicity when treated in culture medium with CAP for 10-40 s, which presented the unique prodrug property of **10**.

**Table 2.**
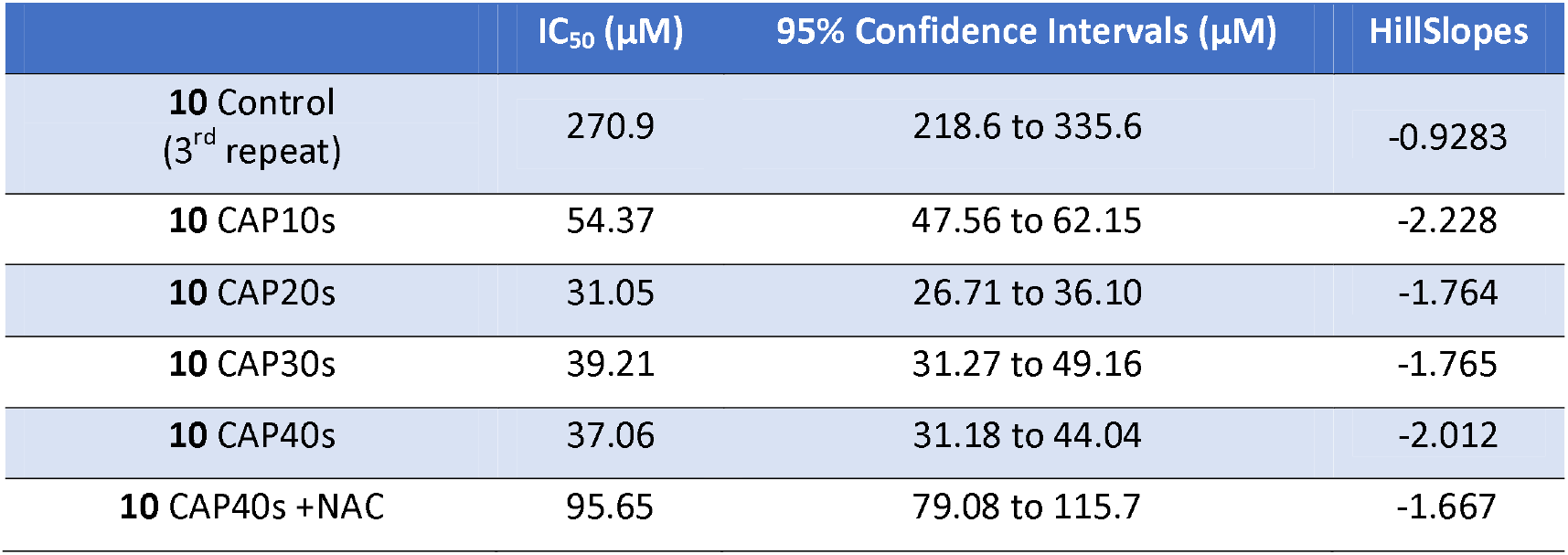
IC_50_ values of 10 treated by CAP 0-40s in culture medium solution.

Our results demonstrate that **10** was activated by reactive species generated during CAP treatment, therefore presenting significant synergistic cytotoxicity to U-251 MG cells combined with CAP treatment. Furthermore, overnight storage CAP-activated medium and CAP-activated cells have been used to study the dose-response of U-251 MG cells to **10**. When cells were treated with overnight storage CAP-activated medium and prodrug, no synergistic cytotoxicity was observed (Supporting Information Figure S36). Meanwhile, the U-251 MG cells have been activated and incubated for 0 h to 5 h after CAP treatment, then incubated with fresh medium containing prodrug. There was no significant synergistic cytotoxicity observed between CAP-activated cells and prodrug compared to the direct combination of CAP treatment and prodrug. However, the CAP-activated cells presented relatively lower cell viability at low prodrug concentration than the direct combination of CAP treatment and prodrug (Supporting Information Figure S36).

## Discussion

Despite improvements in precision surgery, targeted and focused chemotherapy and radiotherapy and quick and accurate diagnosis of tumours, cancer mortality remains a significant global health burden, accounting for almost 10 million deaths in 2018, or 1 in 6 of all mortalities^29^. Current treatments still present several side effects, including poor patient experience, secondary malignancy and a variety of long-term sequelae^2,30–32^. In this case, combination therapy has been considered as a promising way to solve problems in a short time while generating less side effects.

As the altered metabolism and rapid proliferation, the high ROS level in cancer cells is considered a promising target for developing specific therapies^33^. The chemotherapy agents related to ROS can be divided into ROS-trigger drugs and ROS-activated drugs. ROS-trigger drugs can further increase the ROS level in cancer cells beyond their tolerable allowance to trigger cell death, such as fenretinide^34^, nitric oxide-donating aspirin^35^, imexone^36^ and motexafin gadolinium^37^. On the other hand, ROS-activated drugs, also termed as prodrugs, can be activated and become cytotoxic by the relative high ROS level in cancer cells, leading to specific cancer cell killing effects. Meanwhile, CAP, a novel technology allows a localised generation of ROS at a tumour site, which has been demonstrated to have significant therapeutic potential in cancer treatment^1,38–40^. In this case, CAP provides the ideal mechanism to augment or activate ROS-sensitive pro-drugs in targeted areas.

There are various isomeric forms of pyrazolopyrimidines, such as pyrazolo[1,5-a]pyrimidines, pyrazolo[5,1-b]pyrimidines, pyrazolo[3,4-d]pyrimidines and pyrazolo[4,3-d]pyrimidines, which all have found use in drug development and presented potential in antiviral, antimicrobial, anticoccidials, antitumour and anti-inflammatory activities^41^. Pyrazolopyrimidine derivatives have been found to inhibit many protein kinases, such as Src kinase^42^, cyclin-dependent kinases^43,44^, Chk1 Checkpoint kinase^45^, B-Raf protein kinase^46^ and Aurora-A kinase^47^, which are considered significant targets for anti-cancer drug development, as the alterations in many kinase activities are involved in cancerous mutations^48^. For example, Kamal A. et al. (2013) reported a run of aminobenzothiazole linked pyrazolo[1,5-a]pyrimidine conjugates and demonstrated their anti-cancer ability against 5 human cancer cell lines including A549 (lung), ACHN (renal), DU-145 (prostate), Hela (cervical) and MCF-7 (breast)^49^.

Some of them were found to be involved in cellular ROS generation. Tamta et al. (2006) found that a pyrazolopyrimidine derivative (4-amino-6-hydroxypyrazolo-3,4-d-pyrimidine) inhibits xanthine oxidase and decrease the enzymatic formation of uric acid and ROS. Gaonkar S. et al. (2020) reported a novel pyrazolo[3,4-d]pyrimidine derivative (6-Benzyl-3-methyl-1-[4-(4-trifluoromethoxyphenyl)phenyl]-1H-pyrazolo[3,4-d] pyrimidin-4-amine) that was able to generate ROS in NCI-H460 (non-small-cell lung cancer) cells, leading to a loss in mitochondrial membrane potential and apoptosis of the cancer cells^50^. Therefore, the development of new pyrazolopyrimidines as novel efficient anti-cancer drugs is important, especially drugs that may possess synergistic anti-cancer effects with other innovations/treatments.

To the best of our knowledge, we have demonstrated, for the first time, novel pyrazolopyrimidine derivatives, specifically pyrazolopyrimidinones, presented synergistic anti-cancer effects in combination with ROS generated by CAP and may encompass pharmacological potential as anti-cancer prodrugs. We have tested 15 pyrazolopyrimidinones in combination with low dose CAP treatment using culture medium without pyruvate. The pyrazolopyrimidinones were synthesised using a two-step process, Figure 4. This involved generating an isolatable amino pyrazole intermediate (step one) and its subsequent reaction with the corresponding β-ketoester to give the desired pyrazolopyrimidinone (step two)15. A family of 15 pyrazolopyrimidinones was generated, which incorporated structural variation at R1 (R1 = phenyl, substituted aryl, or alkyl groups) and R^2^ (R^2^ = phenyl, substituted aryl, or alkyl groups). The pyrazolopyrimidinones were generated in 16-67% yields and underwent structural characterised using NMR spectroscopy, IR spectroscopy, and mass spectrometry (see Supporting Information).

**Figure 4.**
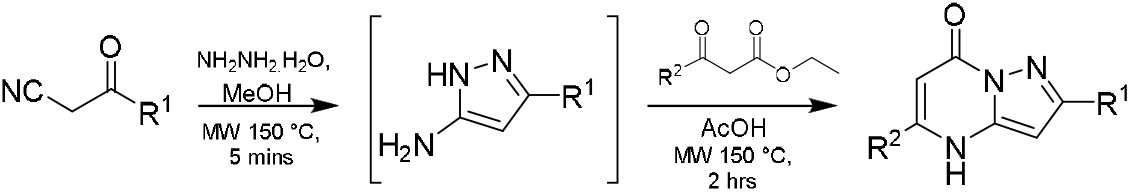
Two-step synthesis of pyrazolopyrimidinones.

We have identified two leading candidates with synergistic cytotoxicity combined with CAP treatment, which are **10** and **9**. Furthermore, as seen in Figure 3, we have investigated the synergistic effects between **10** and CAP treatment. To distinguish the effects induced by CAP to cells and prodrugs, we have investigated the cytotoxicity of CAP-treated pro-drugs without direct and immediate effects from CAP treatment to cells. To remove the direct and immediate effects from CAP to U-251 MG cells, we first treated **10** in DMSO solution. There is no significant synergistic cytotoxicity observed when the CAP treatment was performed to prodrug DMSO solution (Figure 3a). 100% DMSO contains no H_2_O in the solution which impacts generation of ROS^51^. Moreover, DMSO has been demonstrated to be a scavenger of OH radicals^28^, together, this explains the absence of synergistic cytotoxicity of **10** in a DMSO solution without substrate. Therefore, **10** was first diluted in water based DMEM culture medium to corresponding concentrations, then was exposed to CAP treatment before being incubated with cells. Significant synergistic effects were observed in the combination of CAP treatment and **10** treated in culture medium solution (Figure 3b, c and Table 3). NAC has been commonly used as ROS scavenger^52^, and we have found that the cytotoxicity of CAP-treated **10** in culture medium was significantly decreased in culture medium containing NAC (Figure 3b). Our results demonstrate that without direct and immediate effects from CAP to U-251 MG cells, **10** still has significant synergistic cytotoxicity in combination with CAP treatment providing the drug is activated directly by CAP in the culture medium.

It has been demonstrated that short-lived reactive species are removed by overnight storage and most of the long-lived reactive species, such as hydrogen peroxides in the CAP-activated medium are remained^53^. We can determine that the oxidised components in CAP-activated medium do not affect the synergistic cytotoxicity between CAP-treatment and prodrug 10 (Supporting Information Figure S36b). On the other hand, CAP has been found to activate various cellular responses, such as lipid peroxidation^54^, accelerated cellular uptake^25,55^, oxidation of protein/DNA/RNA^38^, alteration of intercellular ROS signalling pathways^56^, and other oxidation stress response^57^. We incubated CAP-activated U-251 MG cells with 10, therefore isolated the compound from ROS, but observed no significant synergistic cytotoxicity (Supporting Information Figure S36c), which demonstrated that the synergistic cytotoxicity is mainly from CAP-activated prodrug rather than CAP-activated cells.

We discovered and demonstrated new compounds that can synergistically kill cancer cells combined with low dose CAP treatment, which has not been previously reported. Our results indicate that these compounds can be directly activated by CAP in tumour cells, or alternatively immediately prior to adding to tumour cells. The detailed activation mechanisms of compounds **10** and **9** will be further investigated in future studies. This research has made a novel contribution to the development of CAP/ROS-trigger prodrug research and will inspire the development of more efficient prodrugs that can be combined with targeted and precise CAP treatment for cancer therapy.

## Acknowledgements

This work is supported by Irish Research Council Government of Ireland Postdoctoral Fellowship Award GOIPD/2020/788 (Z.H., J.C.) and IRCSET EMBARK grant (G.E.C., J.C.); Science Foundation Ireland Grant Numbers 14/IA/2626 (P.C., J.C.) and 17/CDA/4653 (B.T., P.C., J.C.); and by the TU Dublin Fiosraigh Scholarship Programme (S.B., J.M., J.C.); Science Foundation Ireland infrastructure grants 16/RI/3399 and 12/RI/2346/SOF; Maynooth University John Hume Scholarship (M.K.); Government of Ireland Postgraduate Scholarship from the Irish Research Council (R. D.); Maynooth University Teaching Studentship (C.C.).

## Conflict of Interest

The authors declare no conflict of interest.

## Notes

### Competing Interest Statement

The authors have declared no competing interest.

## Bibliography

1. Metelmann, H. R. et al. Head and neck cancer treatment and physical plasma. Clin. Plasma Med. 3, 17–23 (2015).

2. von Woedtke, T., Reuter, S., Masur, K. & Weltmann, K. D. Plasmas for medicine. Phys. Rep. 530, 291–320 (2013).

3. Peng, X. & Gandhi, V. ROS-activated anticancer prodrugs: A new strategy for tumor-specific damage. Ther. Deliv. 3, 823–833 (2012).

4. Welz, C. et al. Cold atmospheric plasma: A promising complementary therapy for squamous head and neck cancer. PLoS One 10, 1–15 (2015).

5. Görmen, M. et al. Synthesis, cytotoxicity, and COMPARE analysis of ferrocene and [3]ferrocenophane tetrasubstituted olefin derivatives against human cancer cells. ChemMedChem 5, 2039–2050 (2010).

6. Utsumi, F. et al. Effect of indirect nonequilibrium atmospheric pressure plasma on anti-proliferative activity against chronic chemo-resistant ovarian cancer cells in vitro and in vivo. PLoS One (2013) doi:10.1371/journal.pone.0081576.

7. Park, S. et al. Cold atmospheric plasma restores paclitaxel sensitivity to paclitaxel-resistant breast cancer cells by reversing expression of resistance-related genes. Cancers (Basel). 11, (2019).

8. Lee, S. et al. Cold atmospheric plasma restores tamoxifen sensitivity in resistant MCF-7 breast cancer cell. Free Radic. Biol. Med. (2017) doi:10.1016/j.freeradbiomed.2017.06.017.

9. Köritzer, J. et al. Restoration of Sensitivity in Chemo - Resistant Glioma Cells by Cold Atmospheric Plasma. PLoS One 8, e64498 (2013).

10. Lin, H. et al. Pyrazolopyrimidine Derivatives as PI3 Kinase Inhibitors. WO Pat WO2013028263 (2013).

11. Gu, B., Block, T. & Cuconati, A. Small Molecule Inhibitors Against West Nile Virus Replication. WO Pat WO2007005541 (2007).

12. Cherukupalli, S. et al. An insight on synthetic and medicinal aspects of pyrazolo[1,5-a]pyrimidine scaffold. European Journal of Medicinal Chemistry vol. 126 298–352 (2017).

13. Griffith, D. A. et al. Discovery and evaluation of pyrazolo[1,5-a]pyrimidines as neuropeptide Y1 receptor antagonists. in Bioorganic and Medicinal Chemistry Letters vol. 21 2641–2645 (2011).

14. McCoull, W. et al. Identification of pyrazolo-pyrimidinones as GHS-R1a antagonists and inverse agonists for the treatment of obesity. Medchemcomm 4, 456–462 (2013).

15. Kelada, M., Walsh, J. M. D., Devine, R. W., McArdle, P. & Stephens, J. C. Synthesis of pyrazolopyrimidinones using a “one-pot” approach under microwave irradiation. Beilstein J. Org. Chem. 14, 122–1228 (2018).

16. Binch, H., Fanning, L., MMD Numa - US Patent 8, 314,239 & 2012, undefined. Modulators of cystic fibrosis transmembrane conductance regulator. Google Patents www.genet.sickkids.on.ca/cftr/ (2012).

17. Venkatesan, A. M. et al. Novel imidazolopyrimidines as dual PI3-Kinase/mTOR inhibitors. Bioorganic Med. Chem. Lett. 20, 653–656 (2010).

18. Folkes, A. J. et al. The identification of 2-(1H-indazol-4-yl)-6-(4-methanesulfonyl-piperazin-1-ylmethyl)-4-morpholin-4-yl-thieno[3,2-d]pyrimidine (GDC-0941) as a potent, selective, orally bioavailable inhibitor of class I PI3 kinase for the treatment of cancer. J. Med. Chem. 51, 5522–5532 (2008).

19. Peat, A. J. et al. Novel pyrazolopyrimidine derivatives as GSK-3 inhibitors. Bioorganic Med. Chem. Lett. 14, 2121–2125 (2004).

20. Martina Ferrari, S. et al. Pyrazolopyrimidine Derivatives as Antineoplastic Agents: with a Special Focus on Thyroid Cancer. Mini-Reviews Med. Chem. 16, 86–93 (2015).

21. O’donnell-Tormey, J., Nathan, C. F., Lanks, K., Deboer, C. J. & De La Harpe, J. Secretion of pyruvate: An antioxidant defense of mammalian cells. J. Exp. Med. 165, 500–514 (1987).

22. Moiseev, T. et al. Post-discharge gas composition of a large-gap DBD in humid air by UV-Vis absorption spectroscopy. Plasma Sources Sci. Technol. 23, (2014).

23. Conway, G. E. et al. Non-thermal atmospheric plasma induces ROS-independent cell death in U373MG glioma cells and augments the cytotoxicity of temozolomide. Br. J. Cancer 114, 435–443 (2016).

24. Chou, T. C. & Martin, N. CompuSyn for drug combinations. A Comput. Softw. Quant. Synerg. Antagon. Determ. IC50, ED50 LD50 Values.[PC Softw. user’s Guid. Paramus, NJ) (2005).

25. He, Z. et al. Cold Atmospheric Plasma Induces ATP-Dependent Endocytosis of Nanoparticles and Synergistic U373MG Cancer Cell Death. Sci. Rep. 8, 1–11 (2018).

26. Babington, P. et al. Use of cold atmospheric plasma in the treatment of cancer. Biointerphases vol. 10 029403 (2015).

27. Tanaka, H. et al. Plasma-activated medium selectively kills glioblastoma brain tumor cells by down-regulating a survival signaling molecule, AKT kinase. Plasma Med. 1, 265–277 (2011).

28. Eberhardt, M. K. & Colina, R. The reaction of OH radicals with dimethyl sulfoxide. A comparative study of fenton’s reagent and the radiolysis of aqueous dimethyl sulfoxide solutions. J. Org. Chem. 53, 1071–1074 (1988).

29. Oppermann, S. et al. High-content screening identifies kinase inhibitors that overcome venetoclax resistance in activated CLL cells. Blood 128, 934–947 (2016).

30. Murray, L. J. & Robinson, M. H. Radiotherapy: Technical aspects. Med. (United Kingdom) 44, 10–14 (2016).

31. Tachibana, K., Feril, L. B. & Ikeda-Dantsuji, Y. Sonodynamic therapy. Ultrasonics 48, 253–259 (2008).

32. Calin, M. A. & Parasca, S. V. Photodynamic therapy in oncology. J. Optoelectron. Adv. Mater. 8, 1173–1179 (2006).

33. Gupta, S. C. et al. Upsides and downsides of reactive oxygen species for Cancer: The roles of reactive oxygen species in tumorigenesis, prevention, and therapy. Antioxidants Redox Signal. 16, 1295–1322 (2012).

34. Sun, S. Y. et al. Mediation of N-(4-hydoxyphenyl)retinamide-induced apoptosis in human cancer cells by different mechanisms. Cancer Res. 59, 2493–2498 (1999).

35. Gao, J., Liu, X. & Rigas, B. Nitric oxide-donating aspirin induces apoptosis in human colon cancer cells through induction of oxidative stress. Proc. Natl. Acad. Sci. U. S. A. 102, 17207–17212 (2005).

36. Hersh, E. M. et al. Antiproliferative and antitumor activity of the 2-cyanoaziridine compound imexon on tumor cell lines and fresh tumor cells in vitro. J. Natl. Cancer Inst. 84, 1238–1244 (1992).

37. Evens, A. M. et al. Motexafin gadolinium generates reactive oxygen species and induces apoptosis in sensitive and highly resistant multiple myeloma cells. Blood 105, 1265–1273 (2005).

38. Suschek, C. V. & Opländer, C. The application of cold atmospheric plasma in medicine: The potential role of nitric oxide in plasma-induced effects. Clin. Plasma Med. 4, 1–8 (2016).

39. Xiang, L., Xu, X., Zhang, S., Cai, D. & Dai, X. Cold atmospheric plasma conveys selectivity on triple negative breast cancer cells both in vitro and in vivo. Free Radic. Biol. Med. 124, 205–213 (2018).

40. Yan, D. et al. Principles of using Cold Atmospheric Plasma Stimulated Media for Cancer Treatment. Sci. Rep. 5, 18339 (2015).

41. Chauhan, M. & Kumar, R. Medicinal attributes of pyrazolo[3,4-d]pyrimidines: A review. Bioorganic Med. Chem. 21, 5657–5668 (2013).

42. Scheppke, L. et al. Retinal vascular permeability suppression by topical application of a novel VEGFR2/Src kinase inhibitor in mice and rabbits. J. Clin. Invest. 118, 2337–2346 (2008).

43. Phillipson, L. J. et al. Discovery and SAR of novel pyrazolo[1,5-a]pyrimidines as inhibitors of CDK9. Bioorganic Med. Chem. 23, 6280–6296 (2015).

44. Heathcote, D. A. et al. A novel pyrazolo[1,5-a]pyrimidine is a potent inhibitor of cyclin-dependent protein kinases 1, 2, and 9, which demonstrates antitumor effects in human tumor xenografts following oral administration. J. Med. Chem. 53, 8508–8522 (2010).

45. Dwyer, M., Paruch, K., Labroli, M., … C. A.-B. & medicinal & 2011, undefined. pyrimidine-based CHK1 inhibitors: A template-based approach—Part 1. Elsevier.

46. Berger, D. M. et al. Non-hinge-binding pyrazolo[1,5-a]pyrimidines as potent B-Raf kinase inhibitors. Bioorganic Med. Chem. Lett. 19, 6519–6523 (2009).

47. Shaaban, M. R., Saleh, T. S., Mayhoub, A. S. & Farag, A. M. Single step synthesis of new fused pyrimidine derivatives and their evaluation as potent Aurora-A kinase inhibitors. Eur. J. Med. Chem. 46, 3690–3695 (2011).

48. Ismail, N. S. M., Ali, G. M. E., Ibrahim, D. A. & Elmetwali, A. M. Medicinal attributes of pyrazolo[1,5-a]pyrimidine based scaffold derivatives targeting kinases as anticancer agents. Futur. J. Pharm. Sci. 2, 60–70 (2016).

49. Kamal, A. et al. Synthesis of pyrazolo[1,5-a]pyrimidine linked aminobenzothiazole conjugates as potential anticancer agents. Bioorganic Med. Chem. Lett. 23, 3208–3215 (2013).

50. Gaonkar, S. et al. Novel pyrazolo[3,4-d]pyrimidine derivatives inhibit human cancer cell proliferation and induce apoptosis by ROS generation. Arch. Pharm. (Weinheim). 353, 1–13 (2020).

51. Lu, P., Boehm, D., Bourke, P. & Cullen, P. J. Achieving reactive species specificity within plasma-activated water through selective generation using air spark and glow discharges. Plasma Process. Polym. 14, 1–9 (2017).

52. Downs, I., Liu, J., Aw, T. Y., Adegboyega, P. A. & Ajuebor, M. N. The ROS scavenger, NAC, regulates hepatic Vα14iNKT cells signaling during Fas mAB-dependent fulminant liver failure. PLoS One 7, (2012).

53. Yan, D. et al. Stabilizing the cold plasma-stimulated medium by regulating medium’s composition. Sci. Rep. 6, (2016).

54. He, Z. et al. Cold Atmospheric Plasma Stimulates Clathrin-Dependent Endocytosis to Repair Oxidised Membrane and Enhance Uptake of Nanomaterial in Glioblastoma Multiforme Cells. Sci. Rep. 10, (2020).

55. Cheng, X. et al. Synergistic effect of gold nanoparticles and cold plasma on glioblastoma cancer therapy. J. Phys. D. Appl. Phys. 47, 335402 (2014).

56. Stoffels, E., Sakiyama, Y. & Graves, D. B. Cold atmospheric plasma: Charged species and their interactions with cells and tissues. IEEE Trans. Plasma Sci. 36, 1441–1457 (2008).

57. Keidar, M., Yan, D. & Sherman, J. H. Cold Plasma Cancer Therapy. Morgan & Claypool Publishers (2019). doi:10.1088/2053-2571/aafb9c.

