## Supplementary Data for "Synergistic Cytotoxicity Between Cold Atmospheric Plasma and Pyrazolopyrimidinones Against Glioblastoma Cells"

^1^BioPlasma Research Group, School of Food Science and Environmental Health, Technological University Dublin, Dublin, Ireland; ^2^Nanolab, FOCAS Research Institute, Technological University Dublin, Dublin, Ireland; ^3^Environmental, Sustainability and Health Research Institute, Technological University Dublin, Dublin, Ireland; ^4^Charles Institute of Dermatology, School of Medicine, University College Dublin, Dublin, Ireland; ^5^Department of Chemistry, Maynooth University, Maynooth, Co. Kildare, Ireland; ^6^In-Vitro Toxicology Group, Institute of Life Science, Swansea University Medical School, Swansea University, Singleton Park, Swansea, Wales, United Kingdom. ^7^Department of Food Biosciences, Teagasc Food Research Centre, Ashtown, Dublin, Ireland; ^8^School of Chemical and Biomolecular Engineering, University of Sydney, Australia; ^9^The Kathleen Lonsdale Institute of Human Health Research, Maynooth University, Maynooth, Co. Kildare, Ireland.

**Corresponding author***

**Table of contents**

1. **Chemistry experimental**	S3
   1.1 **General information**	S3
   1.2 **General procedure for the microwave synthesis of pyrazoles**	S3
       1.2.1 3-Methyl-1H-pyrazol-5-amine	S4
       1.2.2 3-Butyl-1H-pyrazol-5-amine	S4
       1.2.3 3-(*t*-Butyl)-1H-pyrazol-5-amine	S4
   1.3 **General procedure for the one pot synthesis of pyrazolopyrimidones**	S5
       1.3.1 2,5-Diphenylpyrazolo[1,5-a]pyrimidin-7(4H)-one	S5
       1.3.2 5-(4-Methoxyphenyl)-2-phenylpyrazolo[1,5-a]pyrimidin-7(4H)-one	S5
       1.3.3 2-(3-Chlorophenyl)-5-phenylpyrazolo[1,5-a]pyrimidin-7(4H)-one	S6
       1.3.4 2-(4-Fluorophenyl)-5-phenylpyrazolo[1,5-a]pyrimidin-7(4H)-one	S6
       1.3.5 5-Phenyl-2-(p-tolyl)pyrazolo[1,5-a]pyrimidin-7(4H)-one	S6
       1.3.6 2-Methyl-5-phenylpyrazolo[1,5-a]pyrimidin-7(4H)-one	S7
   1.4 **NMR spectra**	S8
   1.5 **References**	S42
2. **Biology experimental**	S43

**1. Chemistry Experimental**

**1.1 General information**

All reagents for synthesis were bought commercially and used without further purification. The 3-amino-5-isosubstitutedpyrazole was purchased from Fluorochem and used as received. Reactions were monitored with thin layer chromatography (TLC) on Merck Silica Gel F_254_ plates. NMR spectra were recorded using Bruker Ascend 500 spectrometer at 293K. All chemical shifts were referenced relative to the relevant deuterated solvent residual peaks or TMS. Assignments of the NMR spectra were deduced using ^1^H NMR and ^13^C NMR, along with 2D experiments (COSY, HSQC and HMBC). The following abbreviations were used to explain the observed multiplicities; s (singlet), d (doublet), t (triplet), q (quartet), m (multiplet), bs (broad singlet), pt (pseudo triplet). Flash chromatography was performed with Merck Silica Gel 60. Microwave reactions were carried out using a CEM Discover Microwave Synthesizer with a vertically focused floor mounted infrared temperature sensor, external to the microwave tube. The 10 mL reaction vessels used were supplied from CEM and were made of borosilicate glass. High resolution mass spectrometry (HRMS) was performed on an Agilent-LC 1200 Series coupled to a 6210 or 6530 Agilent Time-Of-Flight (TOF) mass spectrometer equipped with an electrospray source in both positive and negative (ESI+/−) modes. Infrared spectra were obtained as KBr disks in the region 4000–400 cm^−1^ on a Perkin Elmer Spectrum 100 FT-IR spectrophotometer.

**1.2 General procedure for the microwave synthesis of pyrazoles^1^**

A microwave tube was charged with ketonitrile (2.0 mmol), methanol (1 mL), and hydrazine monohydrate (2.6 mmol) and subjected to microwave irradiation (100 W, 150 °C) for 5 minutes.^1^ Volatiles were subsequently removed under reduced pressure. The residue was purified by either trituration with cold methanol or cyclohexane, or by using column chromatography to give the final product.

**1.2.1 3-Methyl-1H-pyrazol-5-amine**

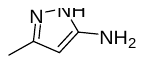

3-Methyl-1H-pyrazol-5-amine was prepared as per general procedure and purified by column chromatography methanol in DCM (0-10%). Light brown oil; Yield (0.097 g, 50%); R_f_ 0.25 (9:1v/v, DCM:MeOH); ^1^H NMR (500 MHz, CDCl_3_) δ 6.14 (bs, 2H), 5.36 (s, 1H), 2.13 (s, 3H); ^13^C NMR (126 MHz, CDCl_3_) δ 153.9, 141.8, 92.0, 11.4; HRMS calcd for C_4_H_8_N_3_ [M + H]^+^: 98.0713, found 98.0717. Matches literature data.^1,3^

**1.2.2 3-Butyl-1H-pyrazol-5-amine**

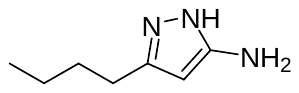

3-Butyl-1H-pyrazol-5-amine was prepared as per general procedure and purified by column chromatography eluting with Methanol in DCM 10%; Yellow oil; Yield (0.054 g, 19%); R_f_ 0.40 (9:1 v/v, DCM:MeOH; ^1^H NMR (500 MHz, CDCl_3_) δ 5.44 (s, 1H), 2.71 – 2.34 (m, 2H), 1.58 (dt, *J* = 15.3, 7.5 Hz, 2H), 1.36 (dq, *J* = 14.7, 7.4 Hz, 2H), 0.92 (t, *J* = 7.4 Hz, 3H); ^13^C NMR (126 MHz, CDCl_3_) ^13^C NMR (126 MHz, CDCl_3_) δ 154.6, 146.1, 91.6, 31.1, 25.9, 22.3, 13.8; C_7_H_14_N_3_ (M + H)+ calc. 140.1182; found 140.1184. Matches literature data.^2^

**1.2.3 3-(*t*-Butyl)-1H-pyrazol-5-amine**

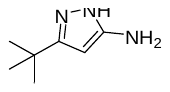

3-(*t*-Butyl)-1H-pyrazol-5-amine was prepared as per general procedure and purified by column chromatography (3:2 EtOAc:Petroleum Ether). Red solid; Yield (0.2155 g, 77%); R_f_ 0.12 (3:2 v/v, EtOAc:Petroleum Ether); ^1^H NMR (500 MHz, CDCl_3_) δ 5.42 (s, 1H), 1.26 (s, 9H,); ^13^C NMR (126 MHz, CDCl_3_) δ 155.2, 154.1, 89.3, 31.0, 30.0; HRMS calcd for C_7_H_14_N_3_ [M + H]^+^: 140.1182, found 140.1185. Matches literature data. ^1,3^

**1.3 General procedure for the one pot synthesis of pyrazolopyrimidones**

Pyrazolopyrimidinones 5, 6. 10, 13, 14, 15 were prepared as per the following literature procedure.^1^ A microwave tube was charged with ketonitrile (0.9 mmol), methanol (1 mL), and hydrazine monohydrate (1.2 mmol) and subjected to microwave irradiation (100 W, 150 °C) for 5 minutes. Subsequently, to this solution was added ketoester (0.9 mmol) and acetic acid (0.5 mmol), and the mixture was subjected to microwave irradiation (100 W, 150 °C) for 2 hours. Volatiles were removed under reduced pressure. The mixture was purified by either trituration with cold methanol or ethyl acetate, or by using column chromatography.

**1.3.1 2,5-Diphenylpyrazolo[1,5-a]pyrimidin-7(4H)-one (6)**

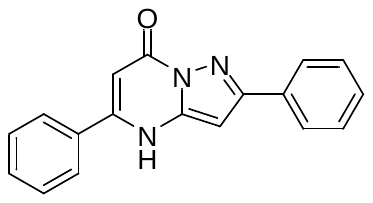

2,5-Diphenylpyrazolo[1,5-a]pyrimidin-7(4H)-one was prepared as per general and purified by trituration with cold MeOH. White solid; Yield (0.135g, 52%); ^1^H NMR (300 MHz, DMSO) δ 12.61 (bs, 1H), 8.01 (m, 2H), 7.87 (m, 2H), 7.60 (m, 3H), 7.46 (m, 3H), 6.67 (s, 1H), 6.10 (s, 1H); ^13^C NMR (75 MHz, DMSO) δ 156.2, 153.3, 149.8, 143.2, 132.4, 132.3, 131.1, 129.0, 128.9, 128.7, 127.2, 126.2, 94.0, 86.6; HRMS calcd for C_18_H_14_N_3_O_2_ [M + H]^+^: 288.1131, found 288.1133. IR (KBr) 3033, 1669, 1611, 1448, 768, 692, 548 cm^-1^. Matches literature data.^1,4^

**1.3.2 5-(4-Methoxyphenyl)-2-phenylpyrazolo[1,5-a]pyrimidin-7(4H)-one (5)**

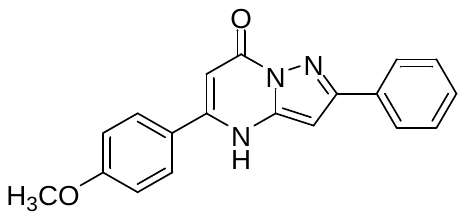

5-(4-Methoxyphenyl)-2-phenylpyrazolo[1,5-a]pyrimidin-7(4H)-one was prepared as per general procedure and purified by column chromatography (1: 1 v/v, EtOAc:Petroluem Ether). Yield (0.0467 g, 16 %); Rf: 0.59 (1:1 v/v, EtOAc:Petroluem Ether); ^1^H NMR (300 MHz, DMSO) δ 8.00 (m, 2H), 7.85 (d, *J* = 8.8 Hz, 2H), 7.47 (m, 3H), 7.14 (d, *J* = 8.8 Hz, 2H), 6.63 (s, 1H), 6.05 (s, 1H), 3.86 (s, 3H); ^13^C NMR (75 MHz, DMSO) δ 161.5, 156.3, 153.0, 149.6, 132.5, 130.7, 128.9, 128.8, 128.7, 126.1, 124.4, 114.4, 92.8, 86.5, 55.5; HRMS calcd for C_19_H_16_N_3_O_2_ [M + H]^+^: 318.1237, found 318.1233; IR (KBr) 3062, 1665, 1607, 1248, 770, 543 cm^-1^. Matches literature data.^1^

**1.3.3. 2-(3-Chlorophenyl)-5-phenylpyrazolo[1,5-a]pyrimidin-7(4H)-one (13)**

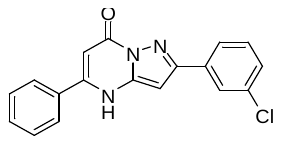

2-(3-Chlorophenyl)-5-phenylpyrazolo[1,5-a]pyrimidin-7(4H)-one was prepared as per general procedure and purified by trituration with cold MeOH. Yield (0.1313 g, 45 %); ^1^H NMR (500 MHz, DMSO) δ 12.69 (bs, 1H), 8.07 (s, 1H), 8.00 (d, *J* = 7.5 Hz, 1H), 7.91 – 7.84 (m, 2H), 7.67 – 7.56 (m, 3H), 7.56 – 7.46 (m, 2H), 6.76 (s, 1H), 6.12 (s, 1H). ^13^C NMR (126 MHz, DMSO) δ 158.9, 156.8, 152.3, 150.7, 135.0, 134.1, 131.6, 131.2, 129.6, 129.2, 127.8, 126.2, 125.3, 94.5, 87.7; HRMS calcd for C_18_H_13_ClN_3_O [M + H]^+^: 322.0742, found 322.0744; IR (KBr) 3062, 1662 (C=O), 1607, 1321, 772, 541 cm^-1^. Matches literature data.^1^

**1.3.4 2-(4-Fluorophenyl)-5-phenylpyrazolo[1,5-a]pyrimidin-7(4H)-one (14)**

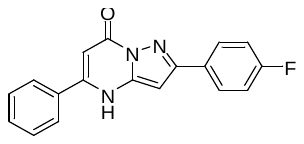

2-(4-Fluorophenyl)-5-phenylpyrazolo[1,5-a]pyrimidin-7(4H)-one was prepared as per general procedure and purified by trituration with cold MeOH. Yield (0.1837 g, 67 %); ^1^H NMR (500 MHz, DMSO) δ 12.61 (bs, 1H), 8.12 – 8.02 (m, 2H), 7.91 – 7.84 (m, 2H), 7.66 – 7.56 (m, 3H), 7.33 (m, 2H), 6.67 (s, 1H), 6.10 (s, 1H). ^13^C NMR (126 MHz, DMSO) δ 163.1 (d, *J*_CF_ = 246.3 Hz), 156.7, 152.9, 150.3, 143.7, 132.8, 131.6, 129.6, 129.4 (d, *J*_CF_ = 3.0 Hz), 128.8 (d, *J*_CF_ = 8.4 Hz), 127.8, 116.2 (d, *J*_CF_ = 21.5 Hz), 94.6, 87.1; HRMS calcd for C_18_H_13_FN_3_O [M + H]^+^: 306. 1037, found 306.1043; IR (KBr) 3060, 1667, 1609, 1528, 767, 694 cm^-1^. Matches literature data.^1,5^

**1.3.5 5-Phenyl-2-(*p*-tolyl)pyrazolo[1,5-a]pyrimidin-7(4H)-one (15)**

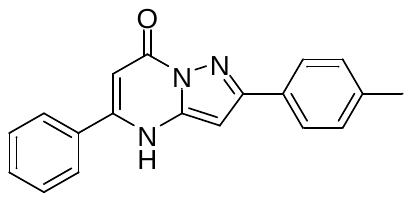

5-Phenyl-2-(*p*-tolyl)pyrazolo[1,5-a]pyrimidin-7(4H)-one was prepared as per general procedure and purified by trituration with cold MeOH. Yield (0.1223 g, 45 %); ^1^H NMR (500 MHz, DMSO) δ 12.57 (bs, 1H), 7.96 – 7.82 (m, 4H), 7.65 – 7.56 (m, 3H), 7.30 (d, *J* = 7.8 Hz, 2H), 6.62 (s, 1H), 6.09 (s, 1H), 2.37 (s, 3H). ^13^C NMR (126 MHz, DMSO) δ 156.7, 153.8, 150.2, 143.6, 138.9, 132.8, 131.5, 130.1, 129.8, 129.5, 127.7, 126.6, 94.5, 86.9, 21.4; HRMS calcd for C_19_H_16_N_3_O [M + H]^+^: 302.1288, found 302.1291. IR (KBr) 3019, 1667, 1609, 1448, 767, 689 cm^-1^. Matches literature data.^1^

**1.3.6 2-Methyl-5-phenylpyrazolo[1,5-a]pyrimidin-7(4H)-one (10)**

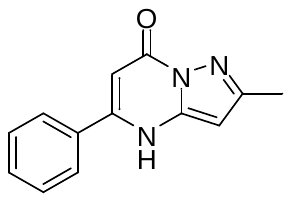

2-Methyl-5-phenylpyrazolo[1,5-a]pyrimidin-7(4H)-one was prepared as per general procedure and purified by trituration with EtOAc. Off-white solid; Yield (0.066 g, 33%); ^1^H NMR (500 MHz, DMSO) δ 12.34 (bs, 1H), 7.86 – 7.81 (m, 2H), 7.62 – 7.55 (m, 3H), 6.05 (s, 1H), 6.00 (s, 1H), 2.32 (s, 3H); ^13^C NMR (126 MHz, DMSO) δ 156.6, 152.6, 149.7, 142.9, 132.9, 131.5, 129.5, 127.6, 94.1, 89.7, 14.6; HRMS calcd for C_13_H_12_N_3_O [M + H]^+^: 226.0975, found 226.0979; IR (KBr) 3441, 3088, 1674, 771 cm^-1^. Matches literature data. ^1,6^

**1.4 NMR spectra**

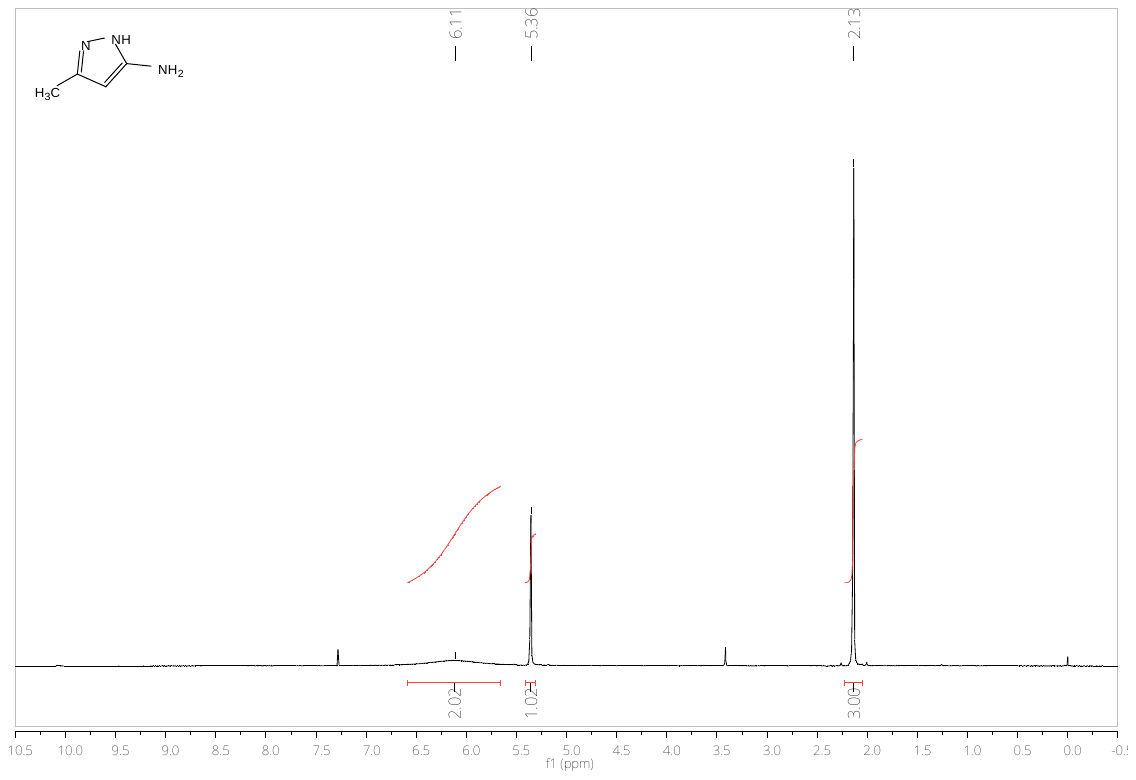

**Figure S1.** ^1^H NMR spectrum of 3-methyl-1H-pyrazol-5-amine

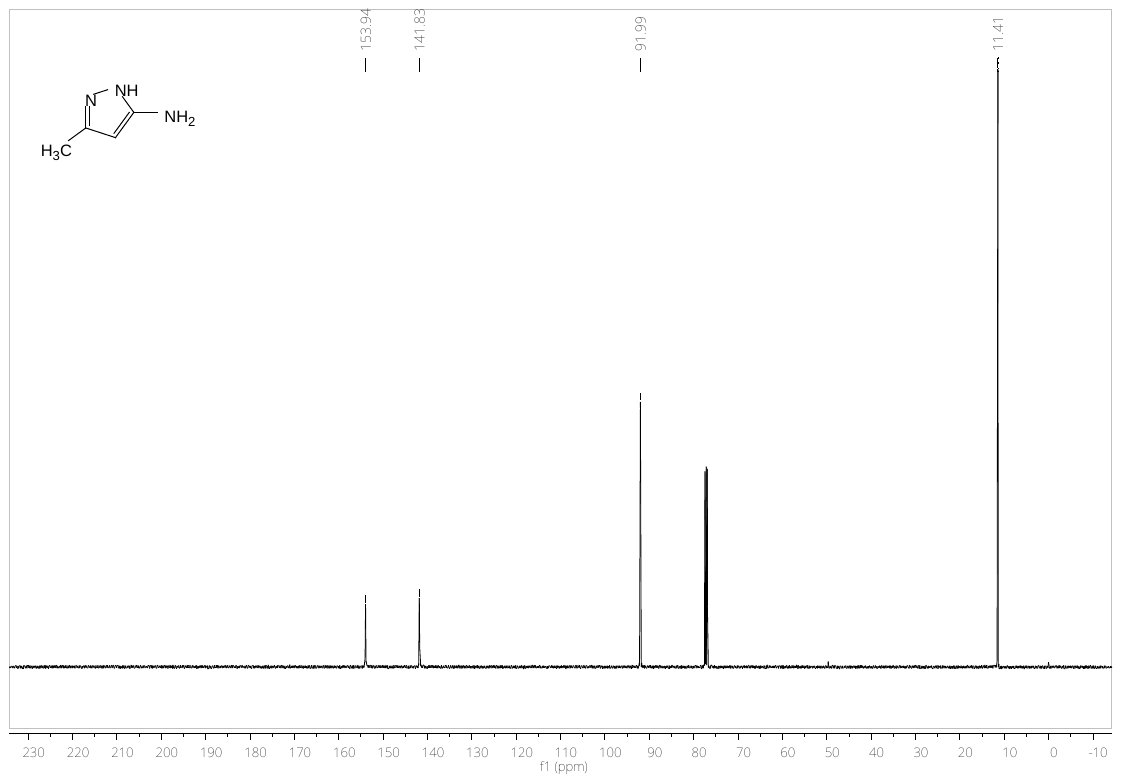

**Figure S2.** ^13^C NMR spectrum of 3-methyl-1H-pyrazol-5-amine

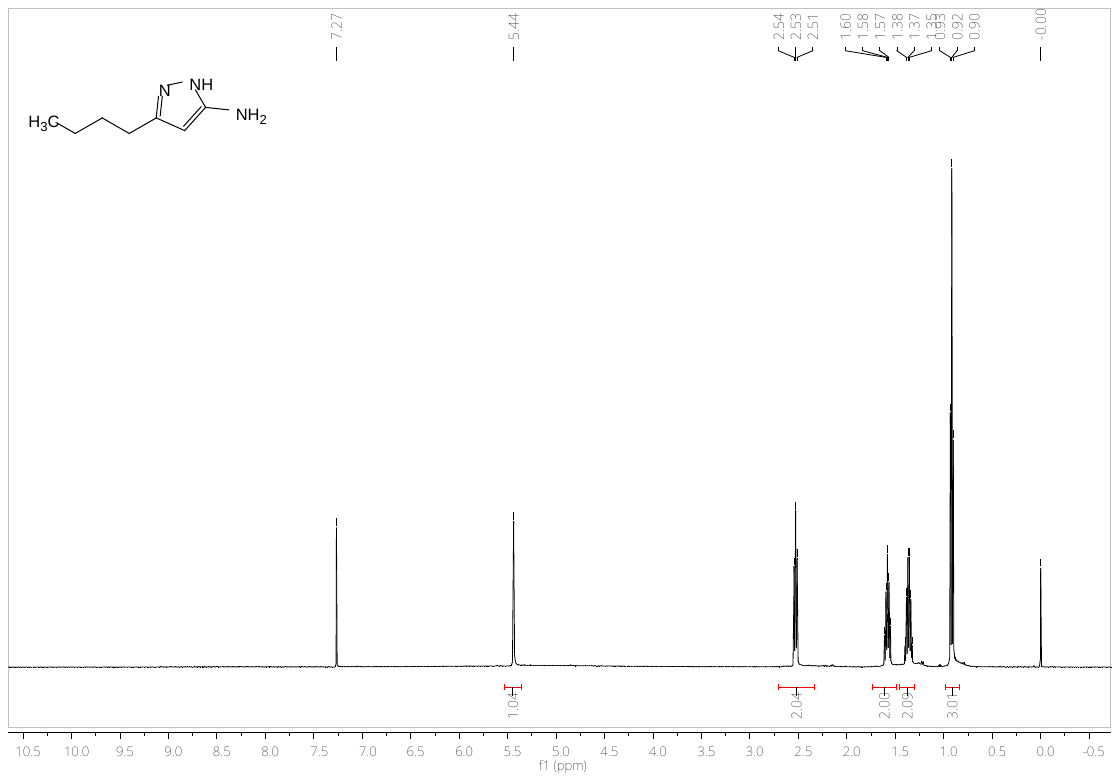

**Figure S3.** ^1^H NMR spectrum of 3-butyl-1H-pyrazol-5-amine

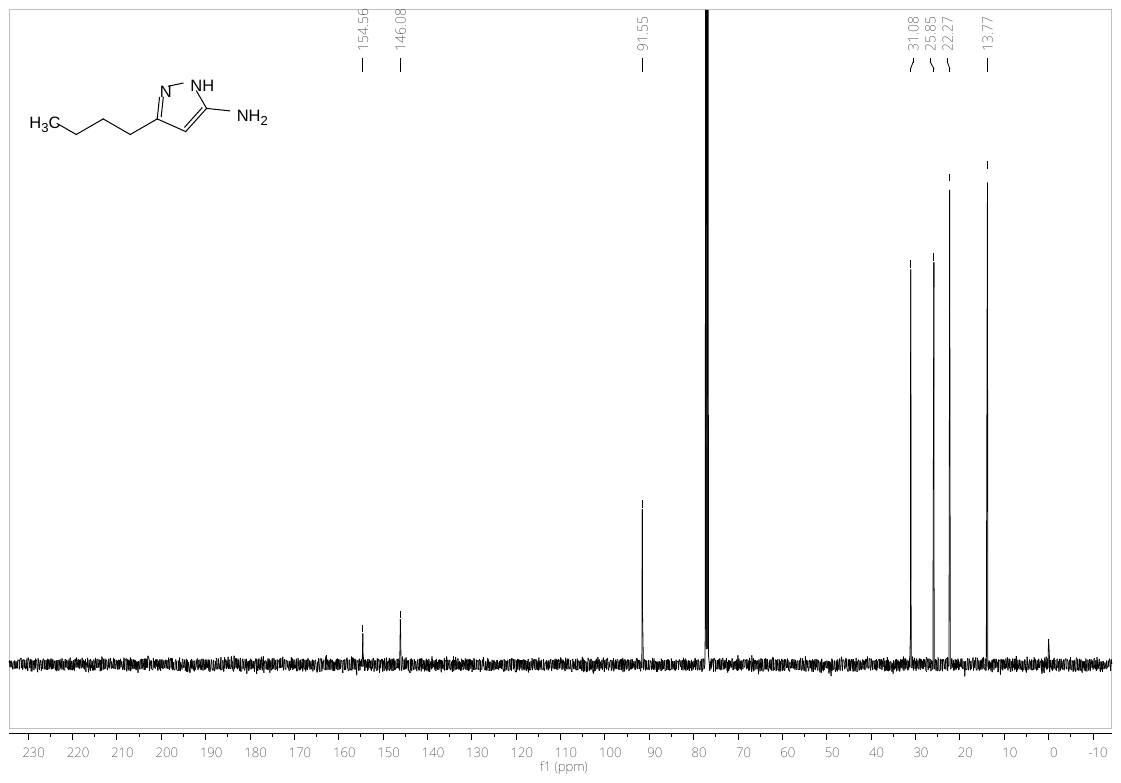

**Figure S4.** ^13^C NMR spectrum of 3-butyl-1H-pyrazol-5-amine

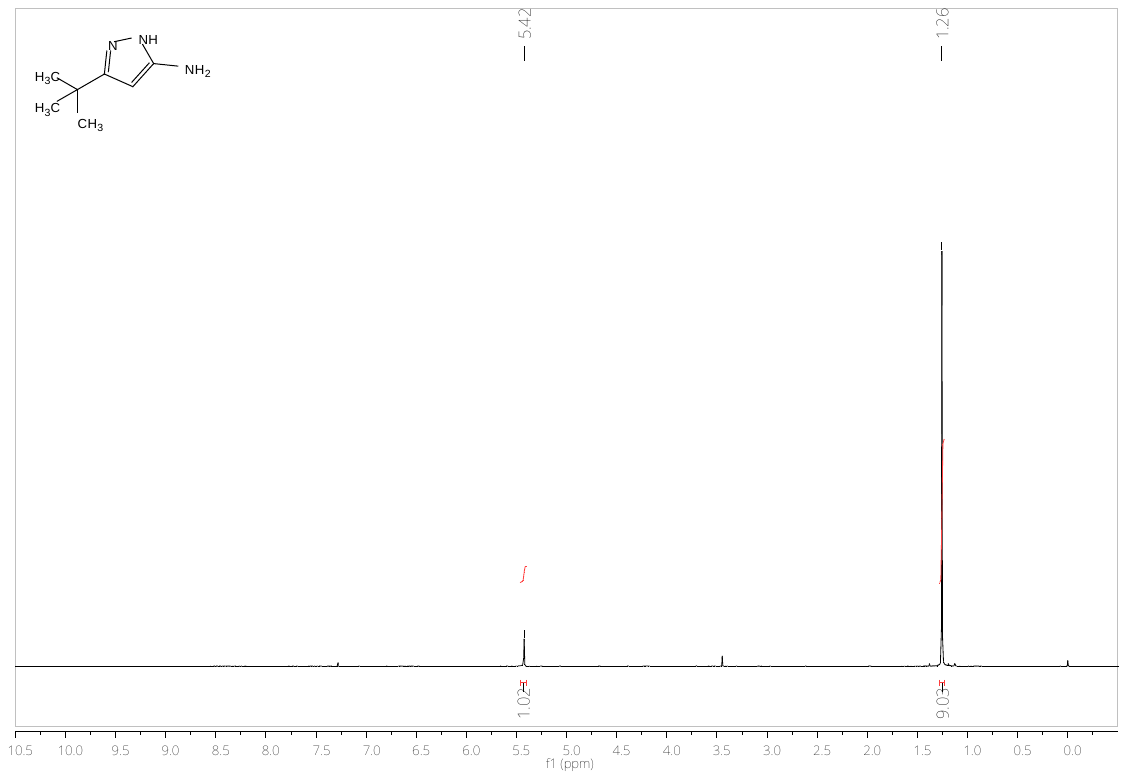

**Figure S5.** ^1^H NMR spectrum of 3-(*t*-butyl)-1H-pyrazol-5-amine

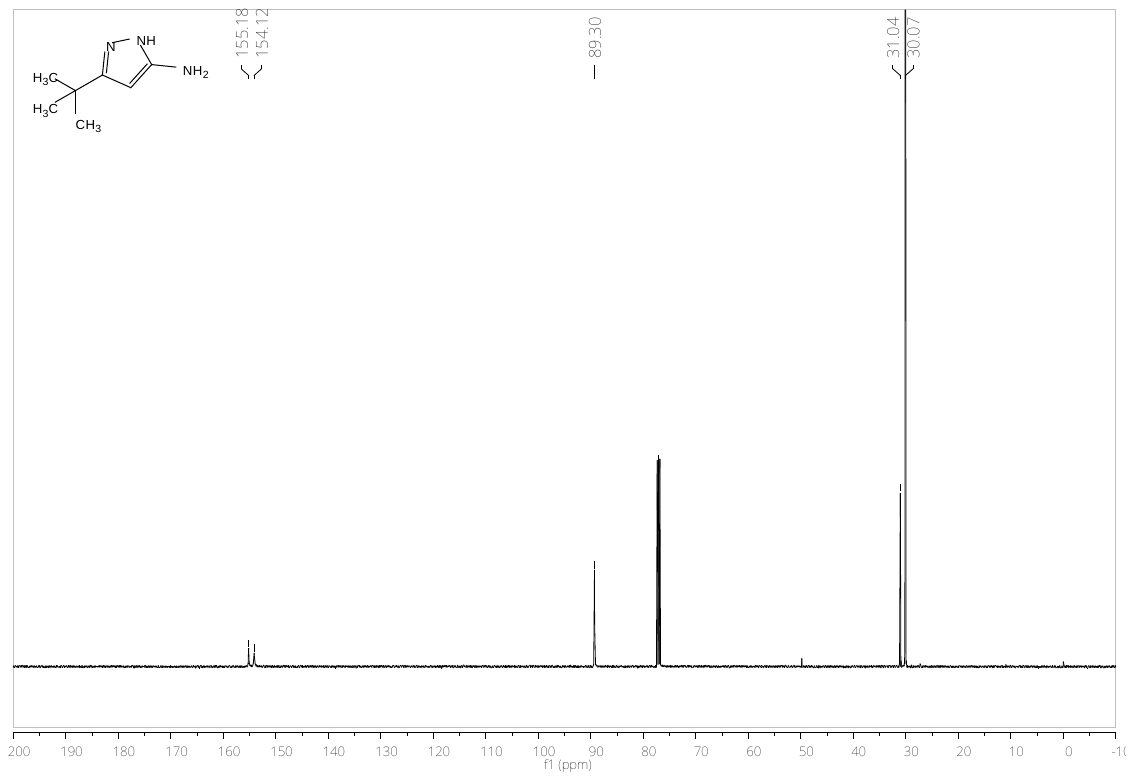

**Figure S6.** ^13^C NMR spectrum of 3-(*t*-butyl)-1H-pyrazol-5-amine

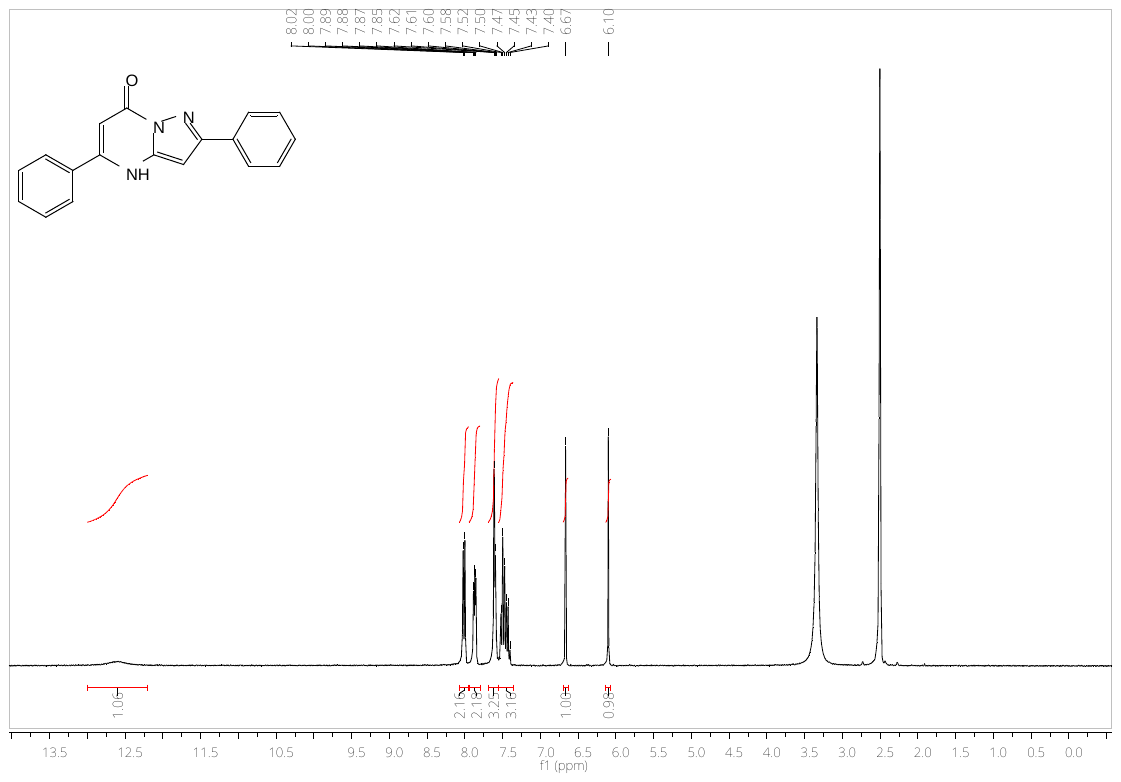

**Figure S7.** ^1^H NMR spectrum of 2,5-diphenylpyrazolo[1,5-a]pyrimidin-7(4H)-one (**6**)

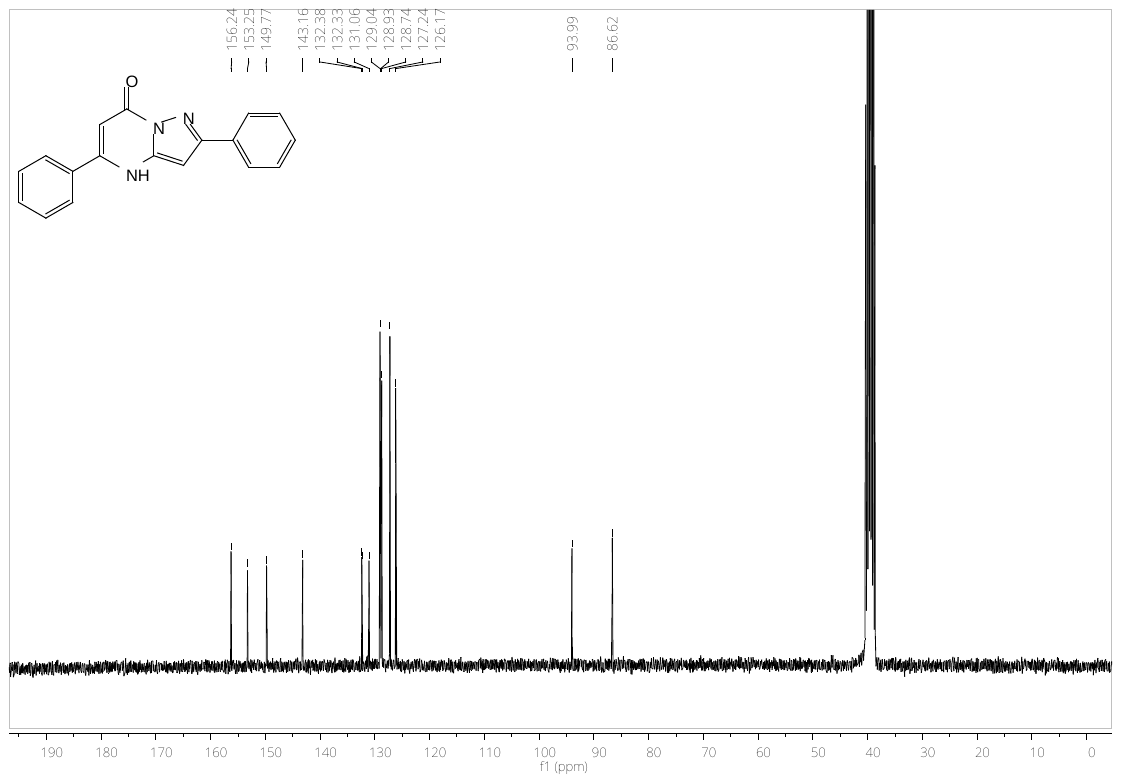

**Figure S8.** ^13^C NMR spectrum of 2,5-diphenylpyrazolo[1,5-a]pyrimidin-7(4H)-one (**6**)

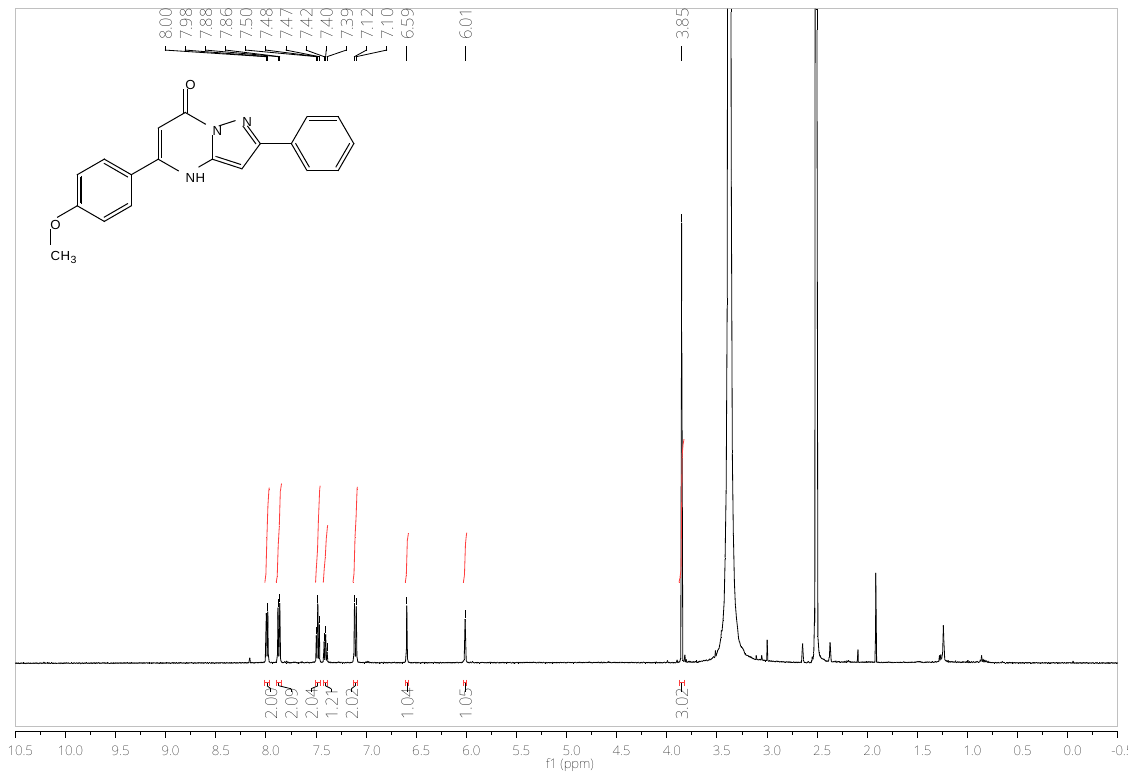

**Figure S9.** ^1^H NMR spectrum of 5-(4-methoxyphenyl)-2-phenylpyrazolo[1,5-a]pyrimidin-7(4H)-one (**5**)

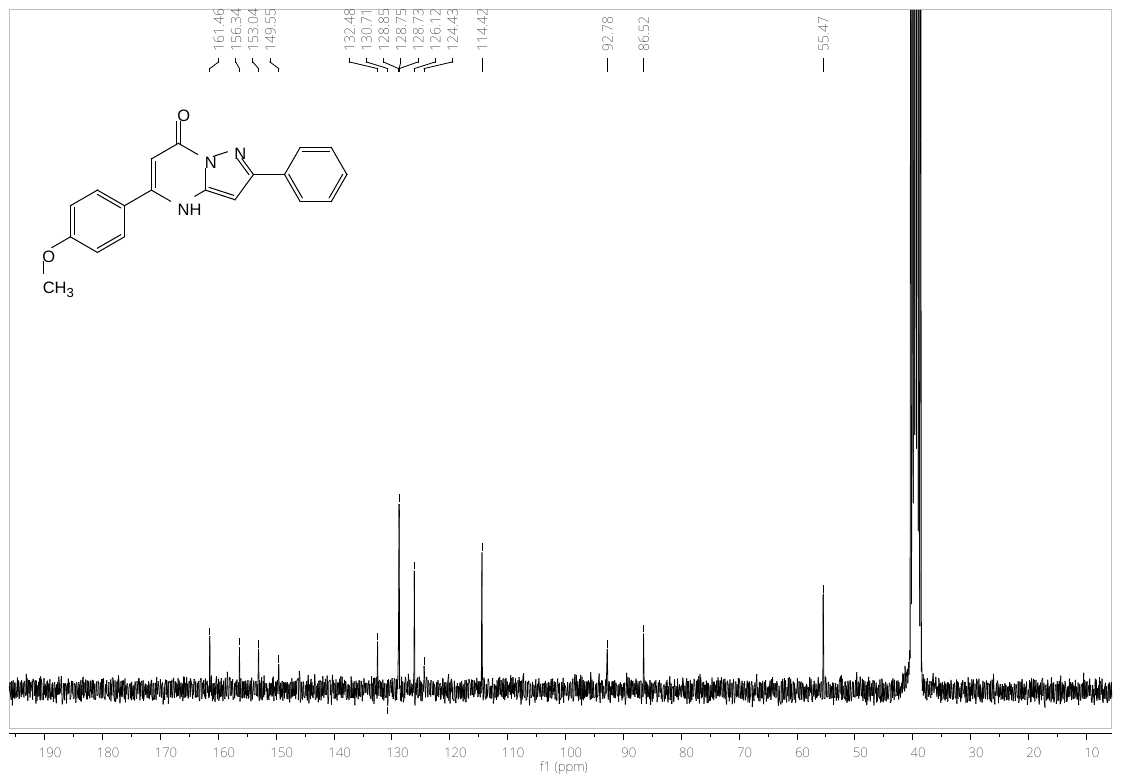

**Figure S10.** ^13^C NMR spectrum of 5-(4-methoxyphenyl)-2-phenylpyrazolo[1,5-a]pyrimidin-7(4H)-one (**5**)

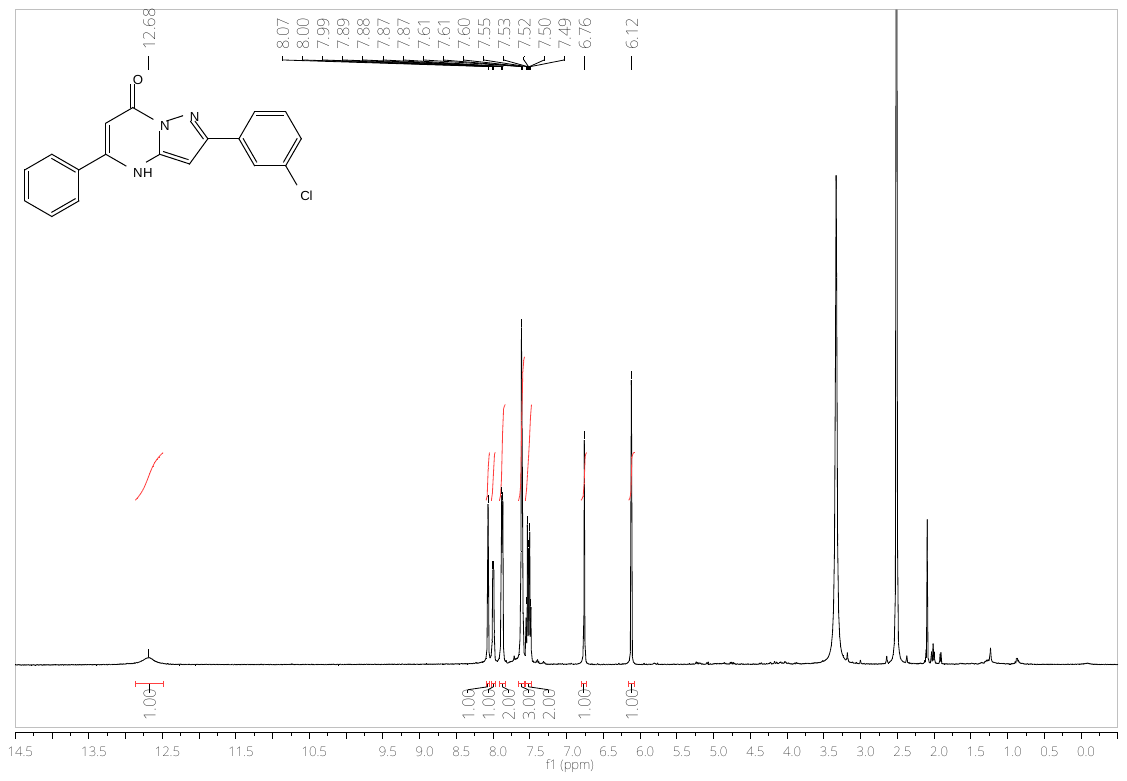

**Figure S11.** ^1^H NMR spectrum of 2-(3-chlorophenyl)-5-phenylpyrazolo[1,5-a]pyrimidin-7(4H)-one (**13**)

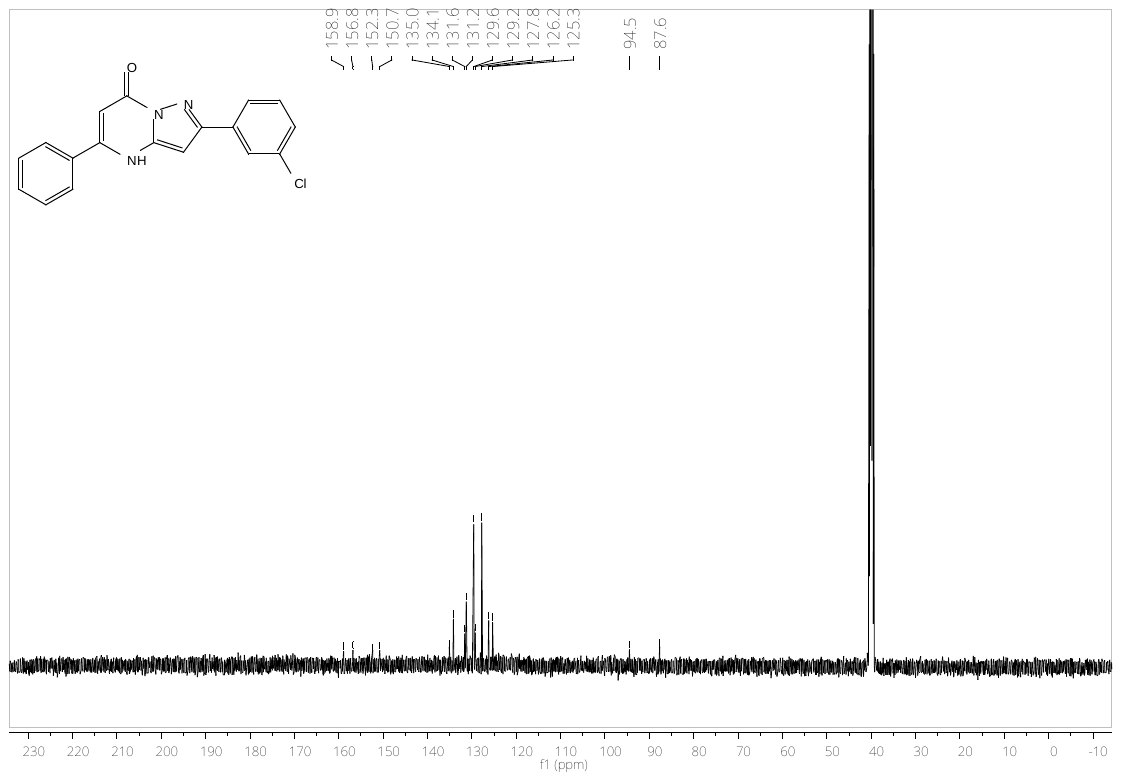

**Figure S12.** ^13^C NMR spectrum of 2-(3-chlorophenyl)-5-phenylpyrazolo[1,5-a]pyrimidin-7(4H)-one (**13**)

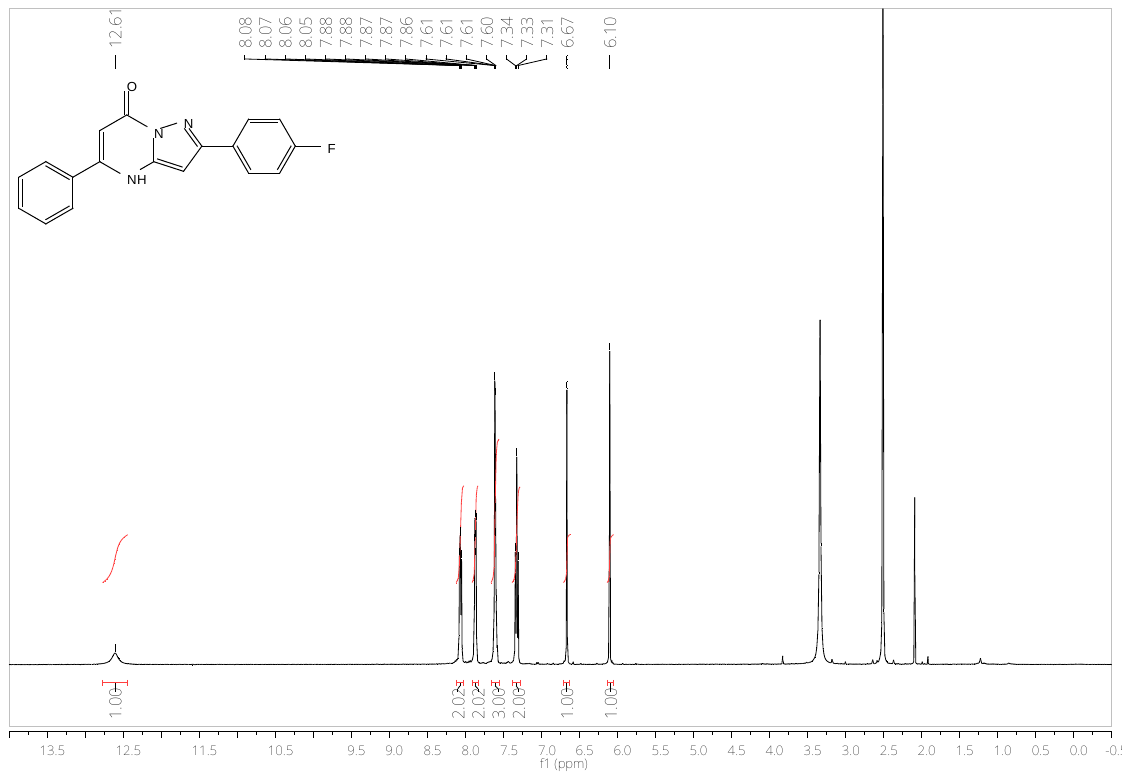

**Figure S13.** ^1^H NMR spectrum of 2-(4-fluorophenyl)-5-phenylpyrazolo[1,5-a]pyrimidin-7(4H)-one (**14**)

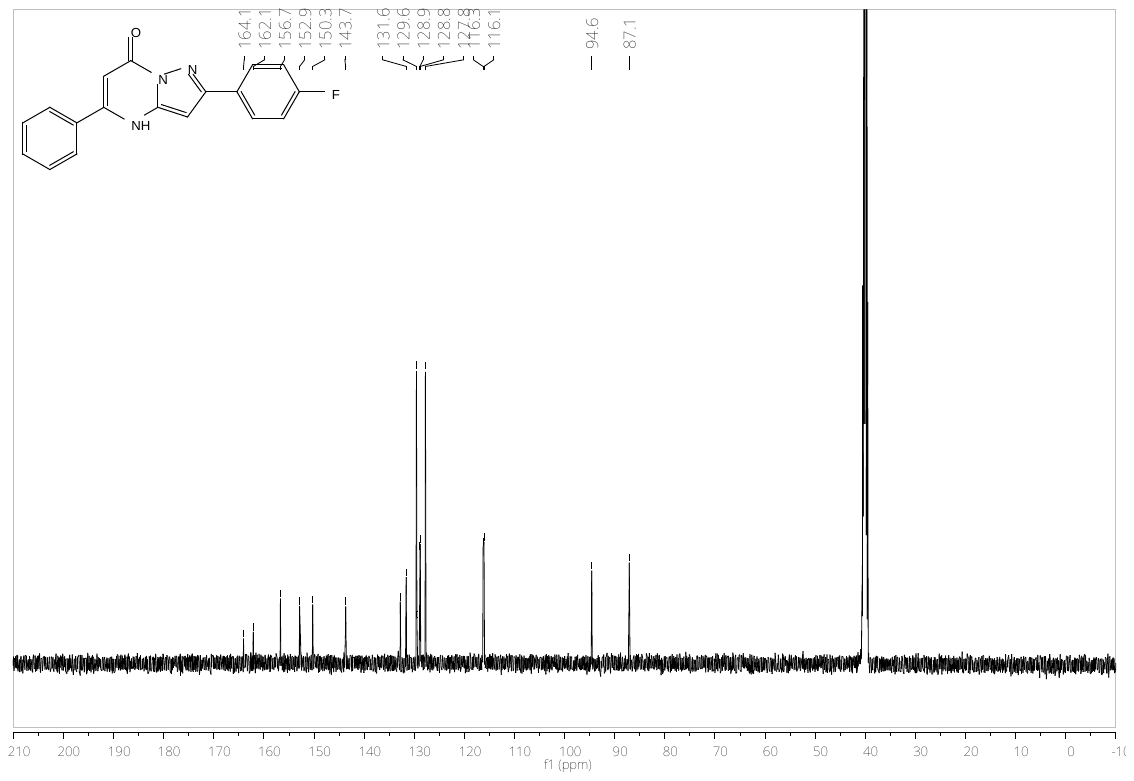

**Figure S14.** ^13^C NMR spectrum of 2-(4-fluorophenyl)-5-phenylpyrazolo[1,5-a]pyrimidin-7(4H)-one (**14**)

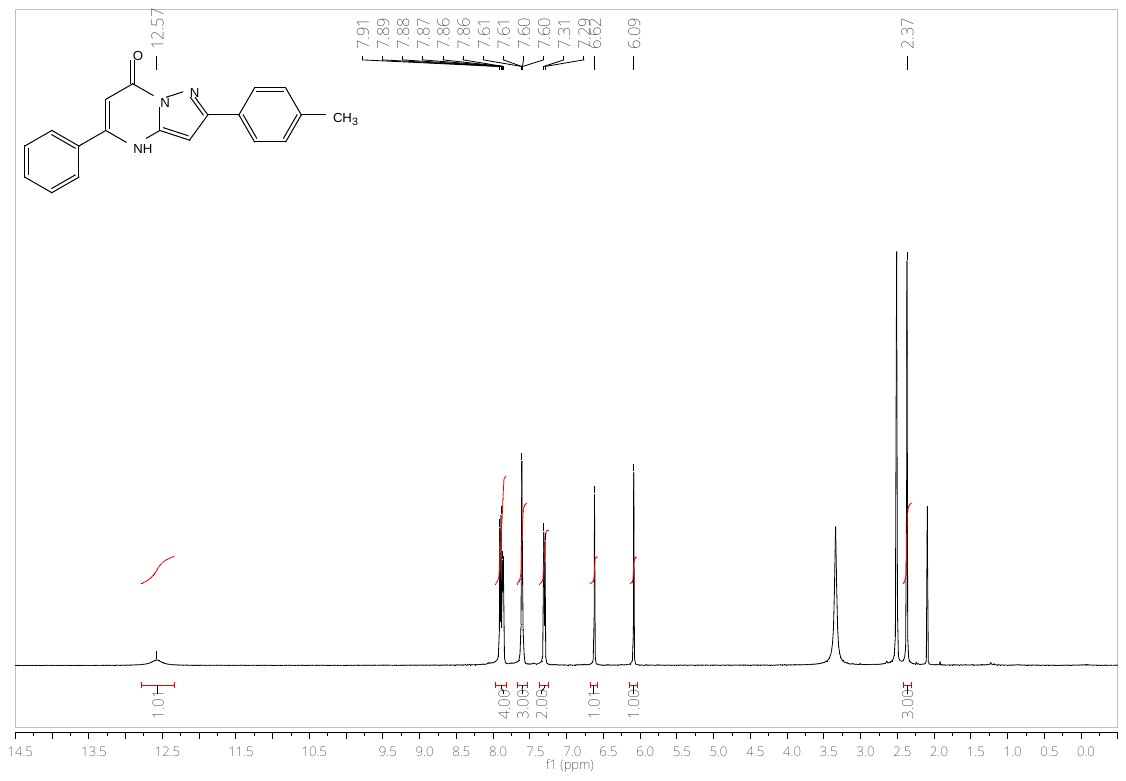

**Figure S15.** ^1^H NMR spectrum of 5-phenyl-2-(p-tolyl)pyrazolo[1,5-a]pyrimidin-7(4H)-one (**15**)

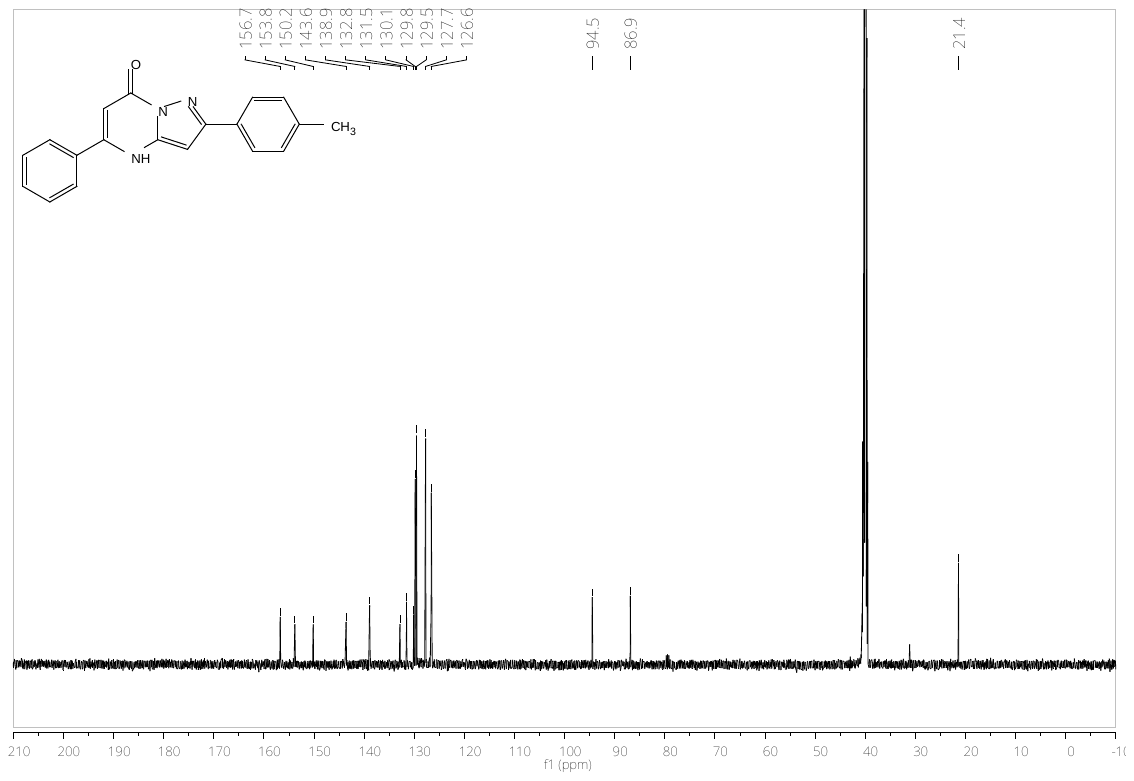

**Figure S16.** ^13^C NMR spectrum of 5-phenyl-2-(p-tolyl)pyrazolo[1,5-a]pyrimidin-7(4H)-one (**15**)

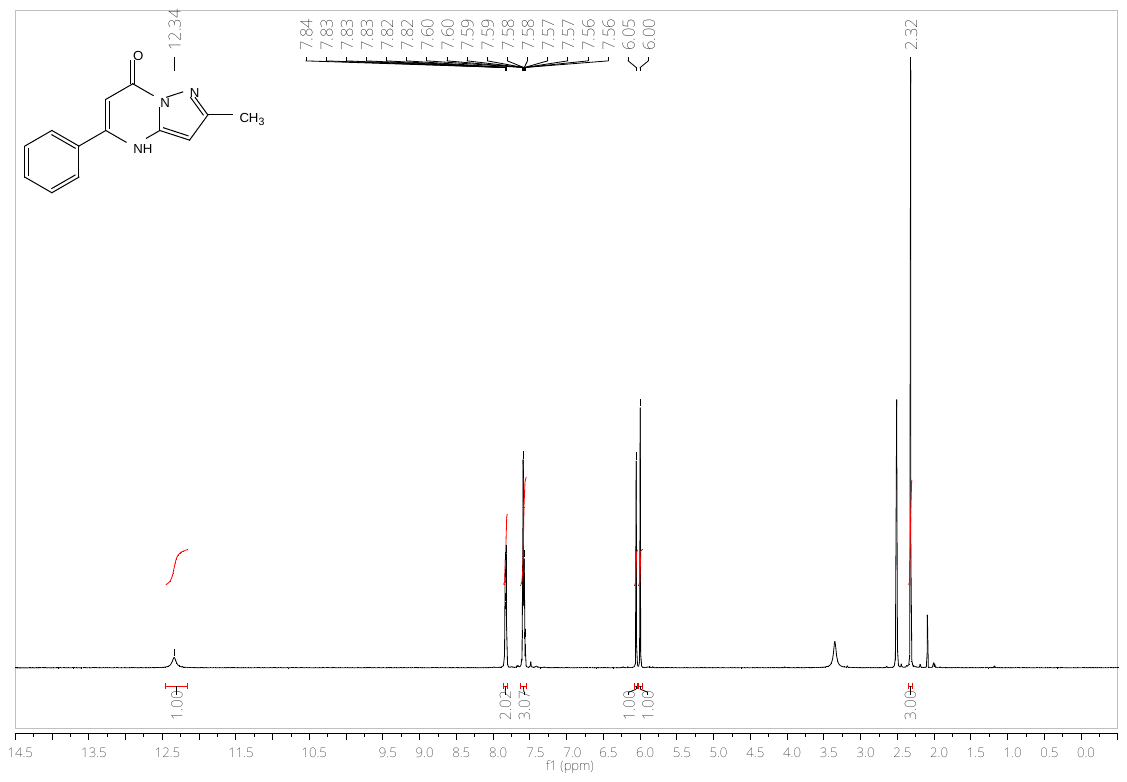

**Figure S17.** ^1^H NMR spectrum of 2-methyl-5-phenylpyrazolo[1,5-a]pyrimidin-7(4H)-one (**10**)

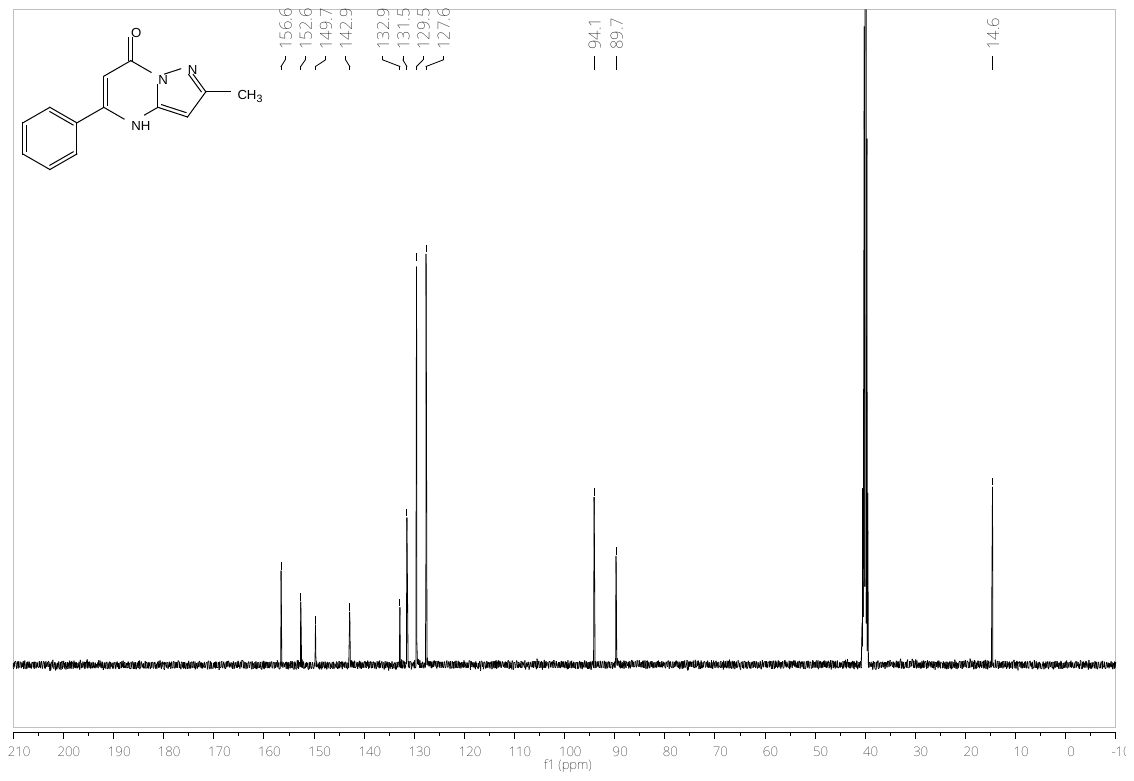

**Figure S18.** ^13^C NMR spectrum of 2-methyl-5-phenylpyrazolo[1,5-a]pyrimidin-7(4H)-one (**10**)

Synthesis of 5-(3,5-dimethylphenyl)-2-isopropylpyrazolo[1,5*-a*]pyrimidin-7(4*H*)-one (MK6) (58) mark thesis

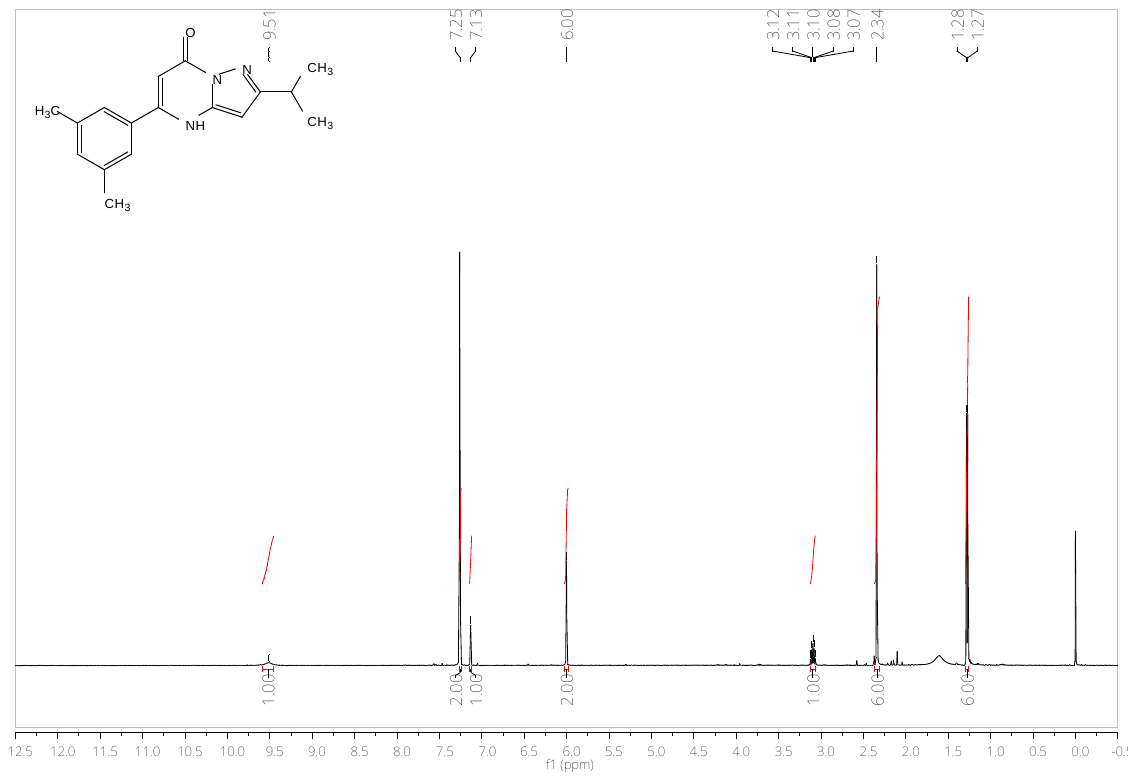

**Figure S19.** ^1^H NMR spectrum of 5-(3,5-dimethylphenyl)-2-isopropylpyrazolo[1,5-a]pyrimidin-7(4H)-one (**12**)

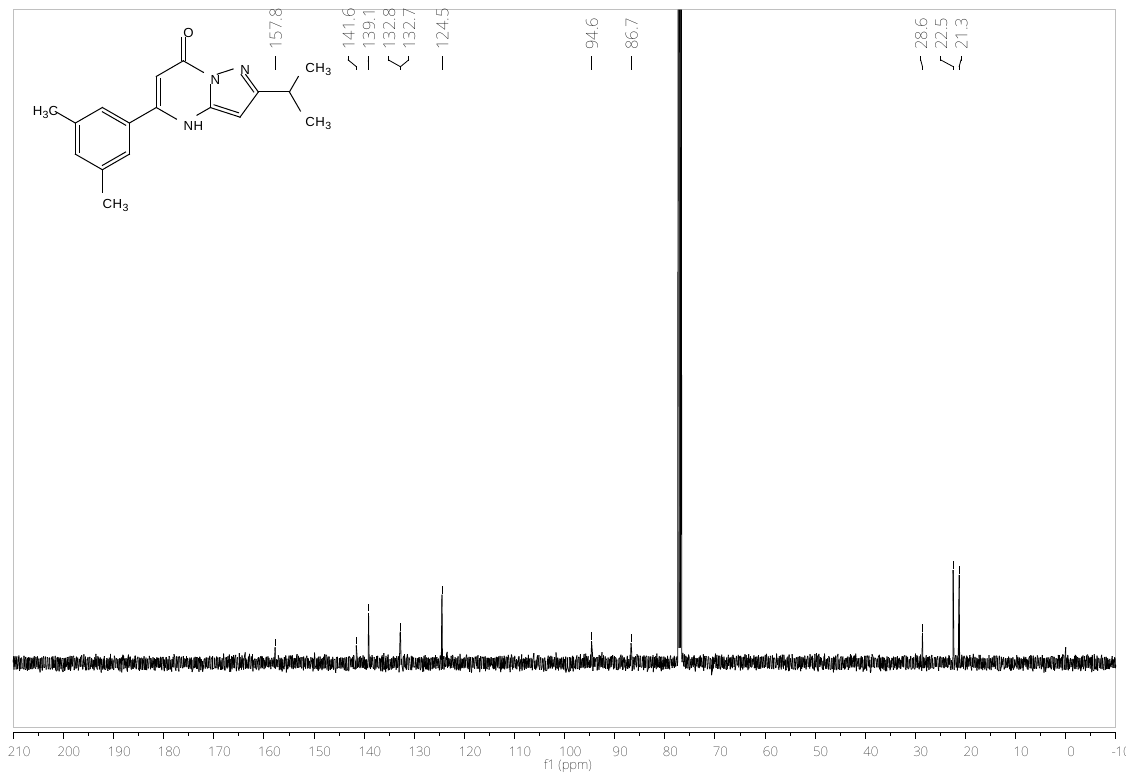

**Figure S20.** ^13^C NMR spectrum of 5-(3,5-dimethylphenyl)-2-isopropylpyrazolo[1,5-a]pyrimidin-7(4H)-one (**12**)

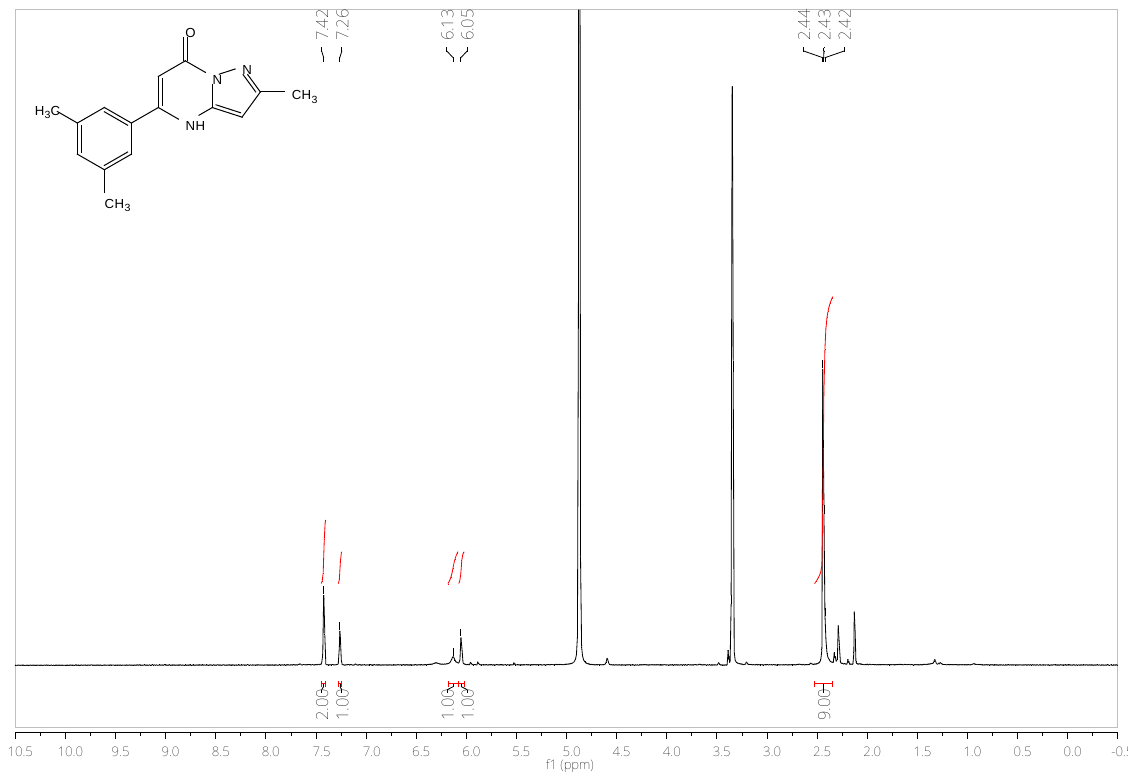

**Figure S21.** ^1^H NMR spectrum of 5-(3,5-dimethylphenyl)-2-methylpyrazolo[1,5-a]pyrimidin-7(4H)-one (**11**)

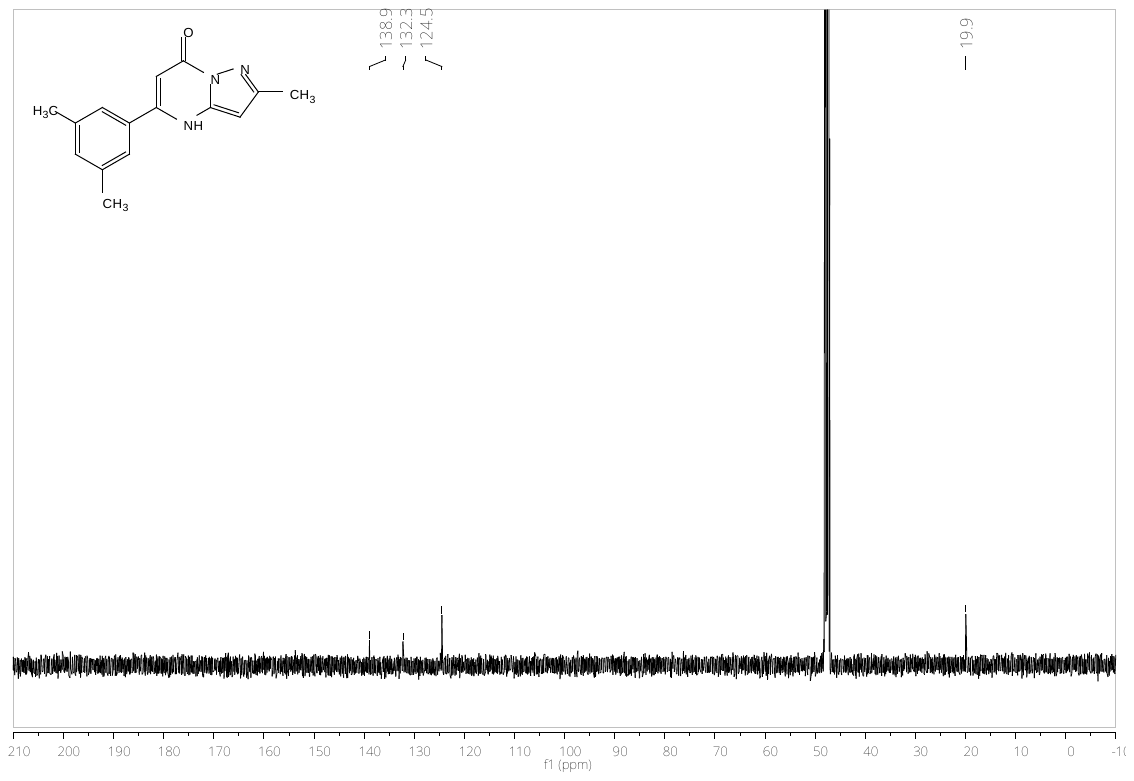

**Figure S22.** ^13^C NMR spectrum of 5-(3,5-dimethylphenyl)-2-methylpyrazolo[1,5-a]pyrimidin-7(4H)-one (**11**)

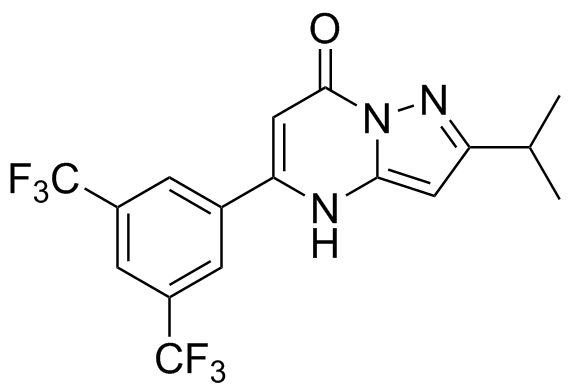

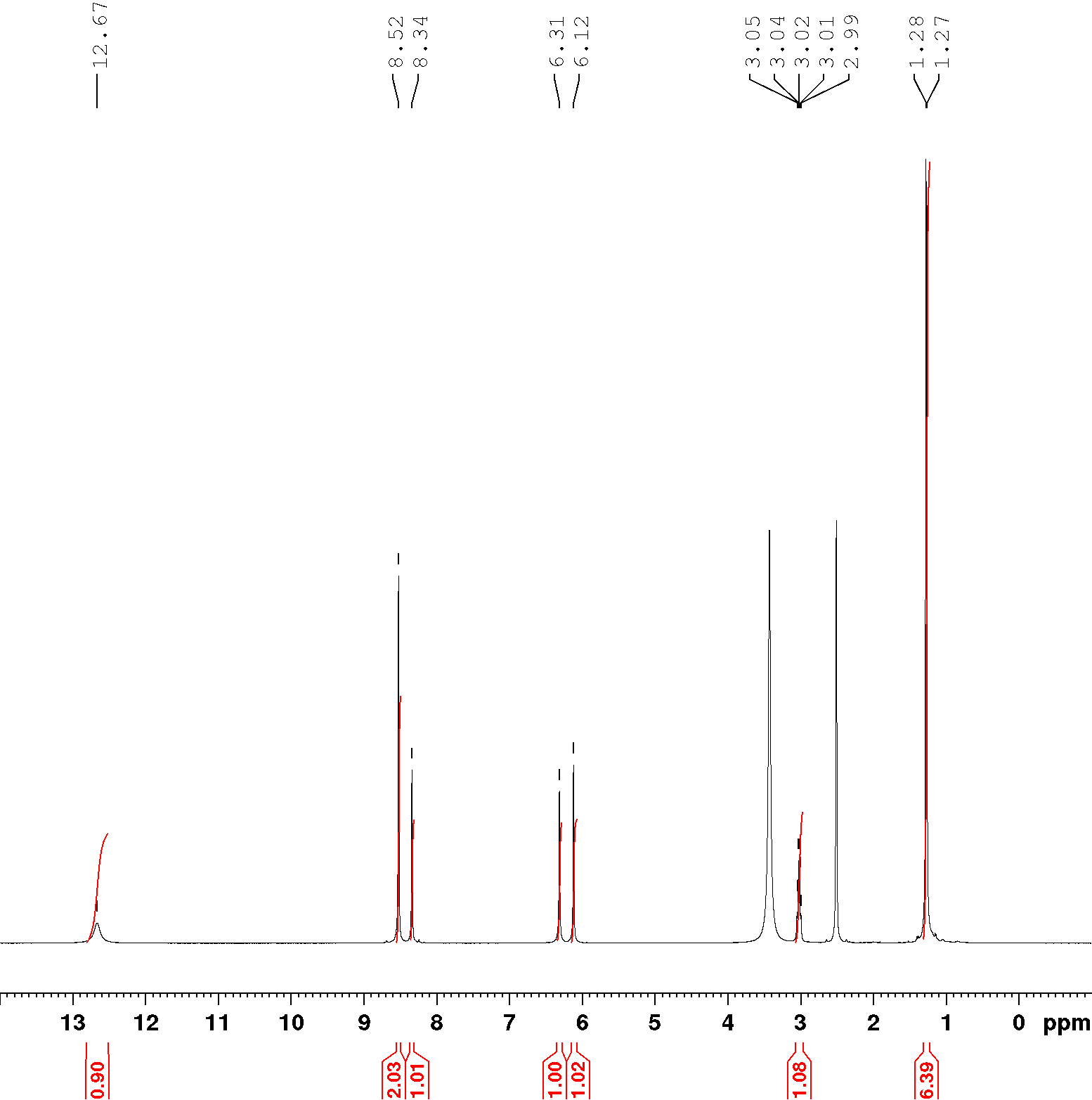

**Figure S23.** ^1^H NMR spectrum of 5-(3,5-bis(trifluoromethyl)phenyl)-2-isopropylpyrazolo[1,5-a]pyrimidin-7(4H)-one (**7**)

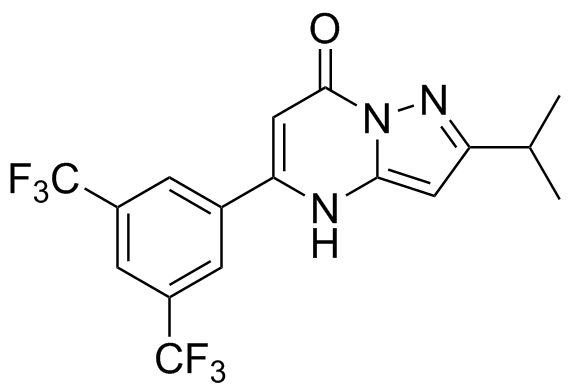

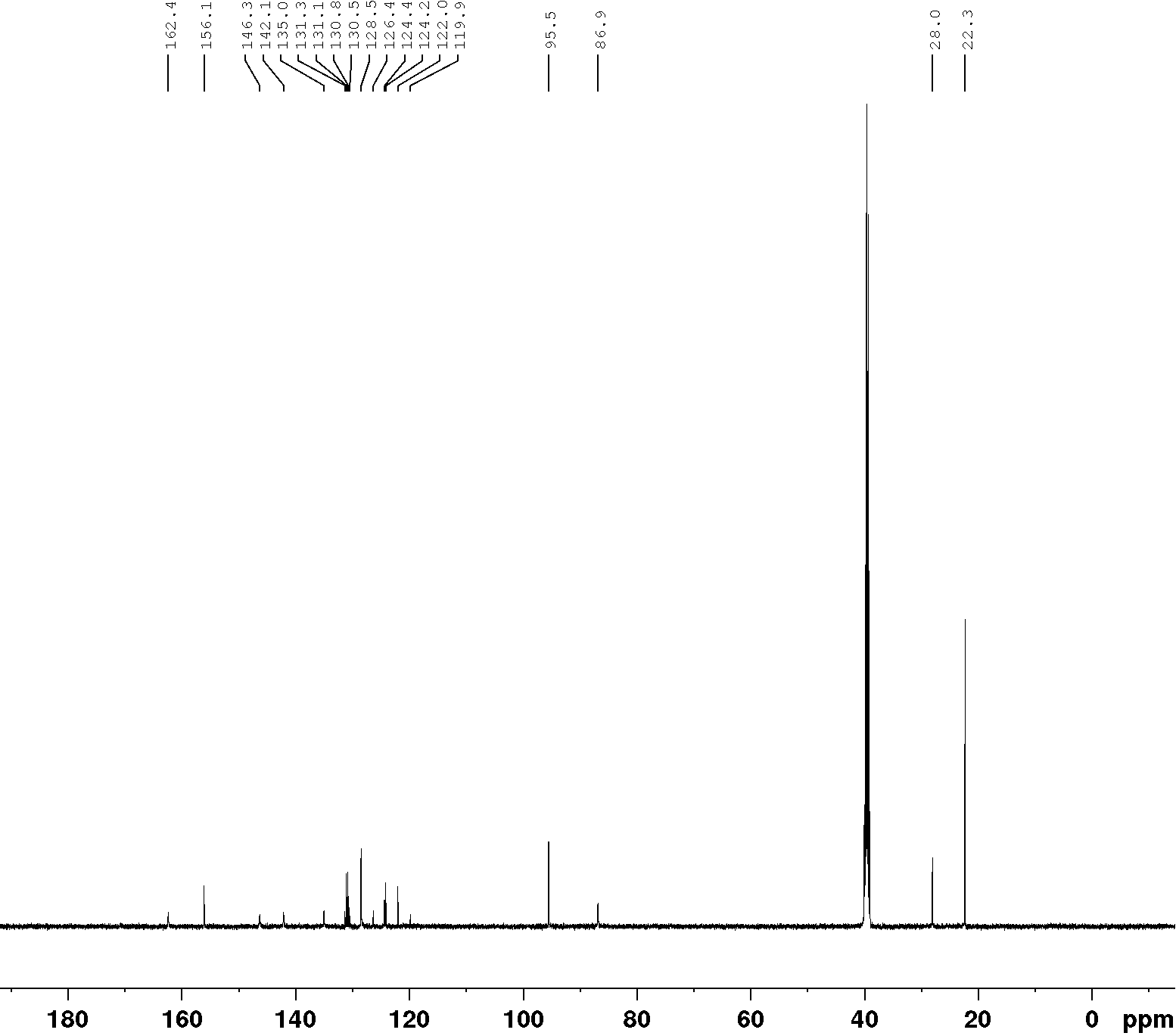

**Figure S24.** ^13^C NMR spectrum of 5-(3,5-bis(trifluoromethyl)phenyl)-2-isopropylpyrazolo[1,5-a]pyrimidin-7(4H)-one (**7**)

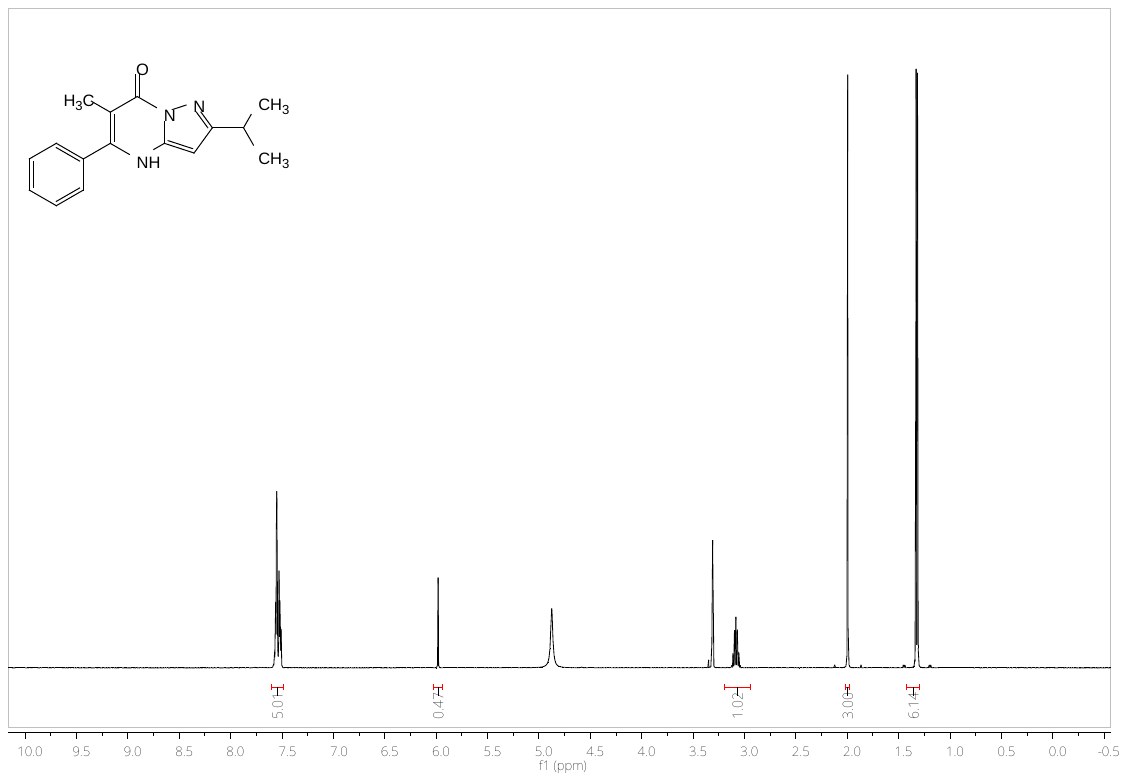

**Figure S25.** ^1^H NMR spectrum of 2-isopropyl-6-methyl-5-phenylpyrazolo[1,5-a]pyrimidin-7(4H)-one (**8**)

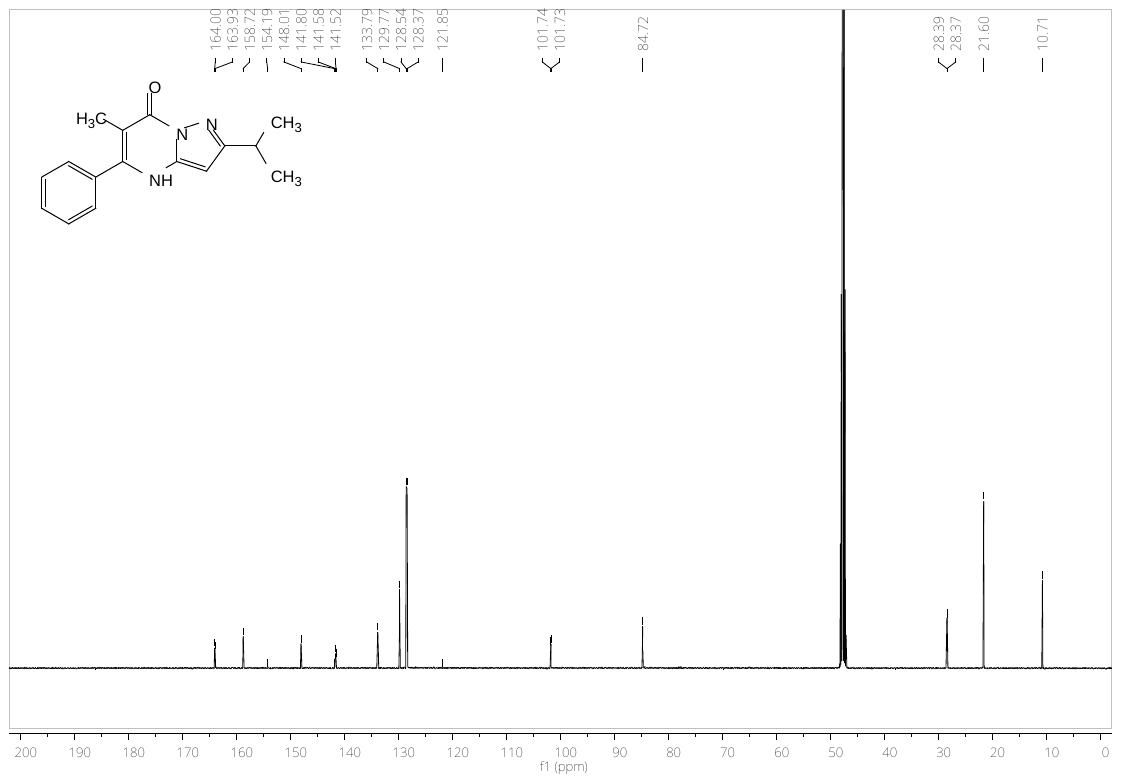

**Figure S26.** ^13^C NMR spectrum of 2-isopropyl-6-methyl-5-phenylpyrazolo[1,5-a]pyrimidin-7(4H)-one (**8**)

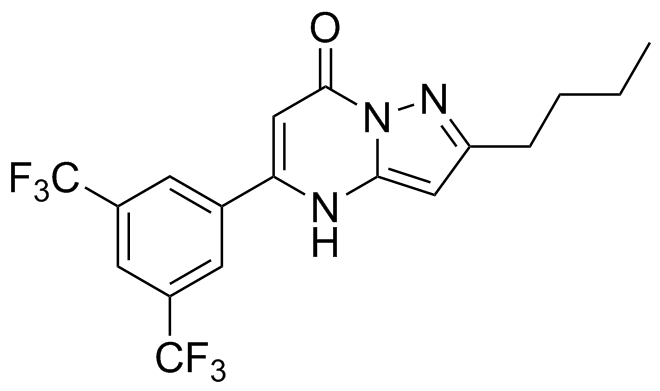

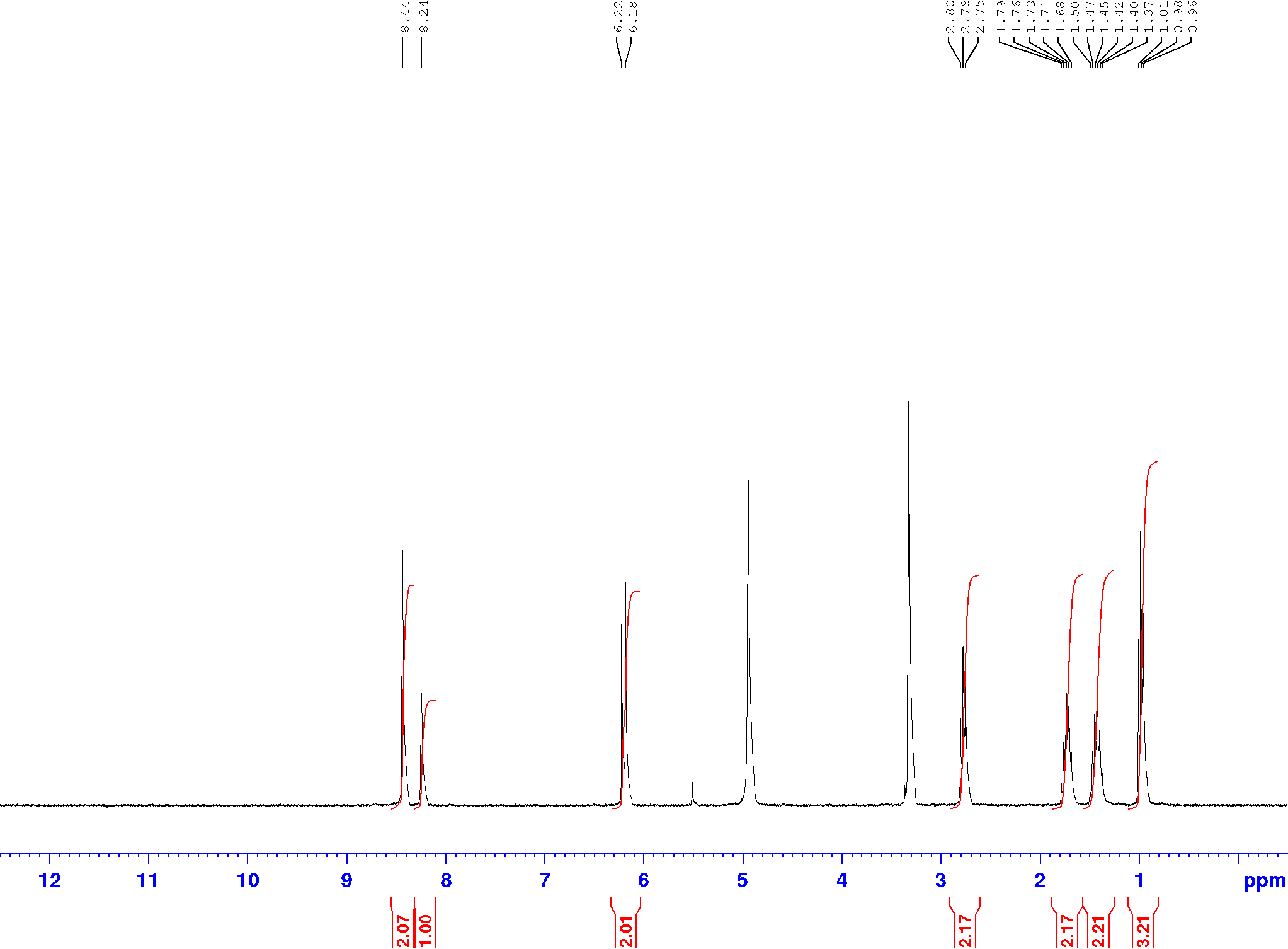

**Figure S27.** ^1^H NMR spectrum of 5-(3,5-bis(trifluoromethyl)phenyl)-2-butylpyrazolo[1,5-a]pyrimidin-7(4H)-one (**2**)

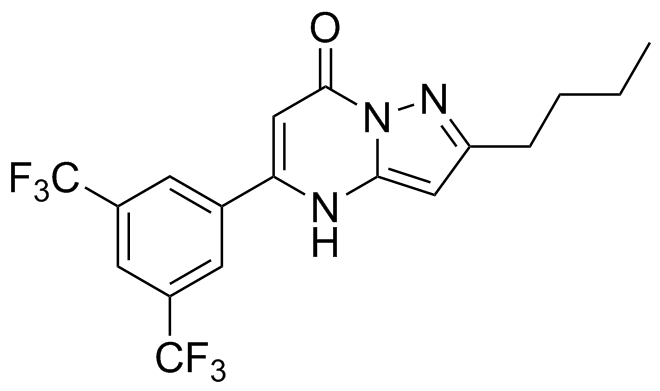

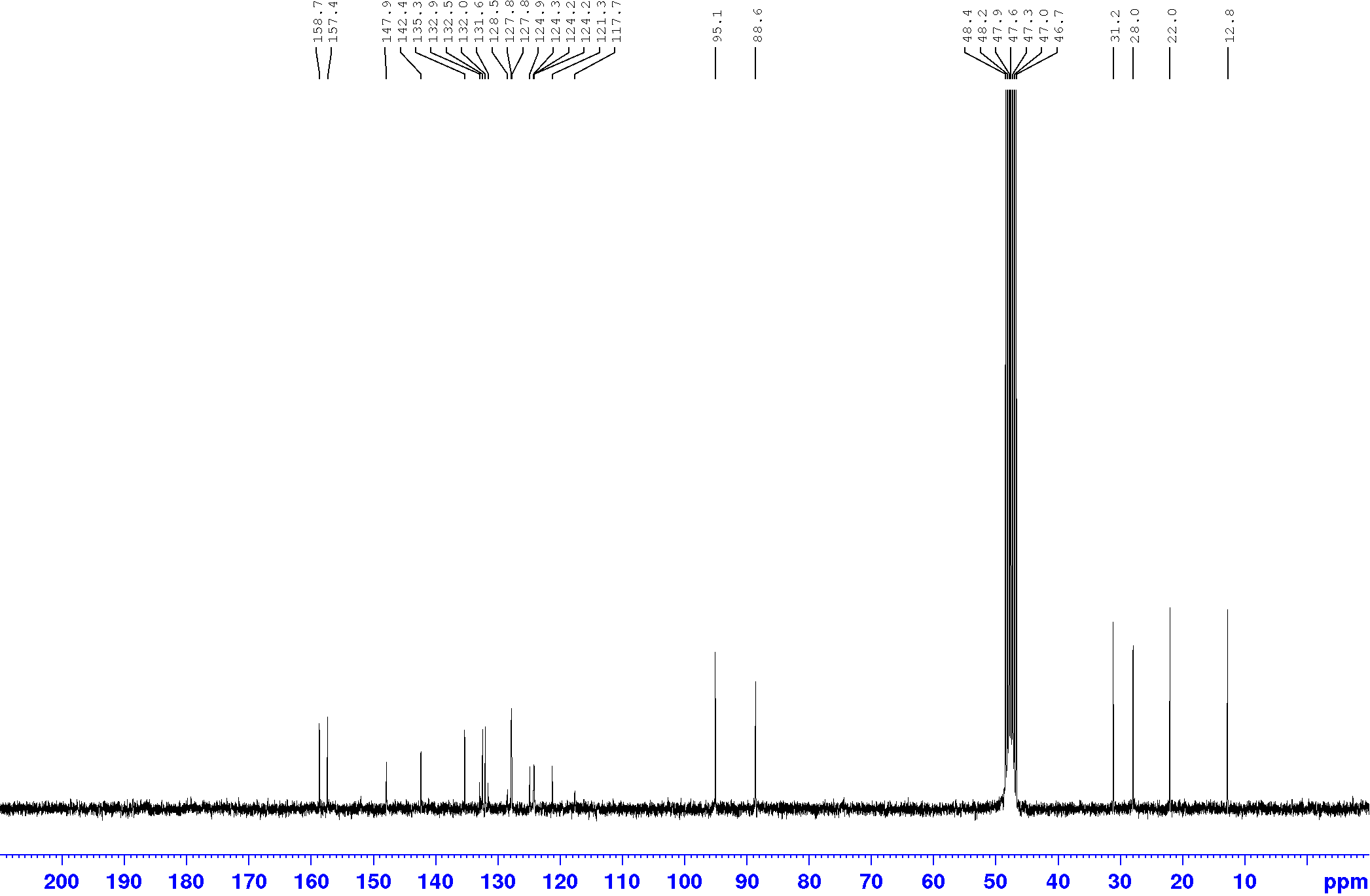

**Figure S28.** ^13^C NMR spectrum of 5-(3,5-bis(trifluoromethyl)phenyl)-2-butylpyrazolo[1,5-a]pyrimidin-7(4H)-one (**2**)

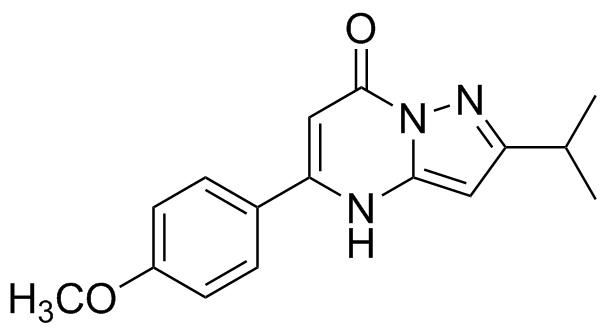

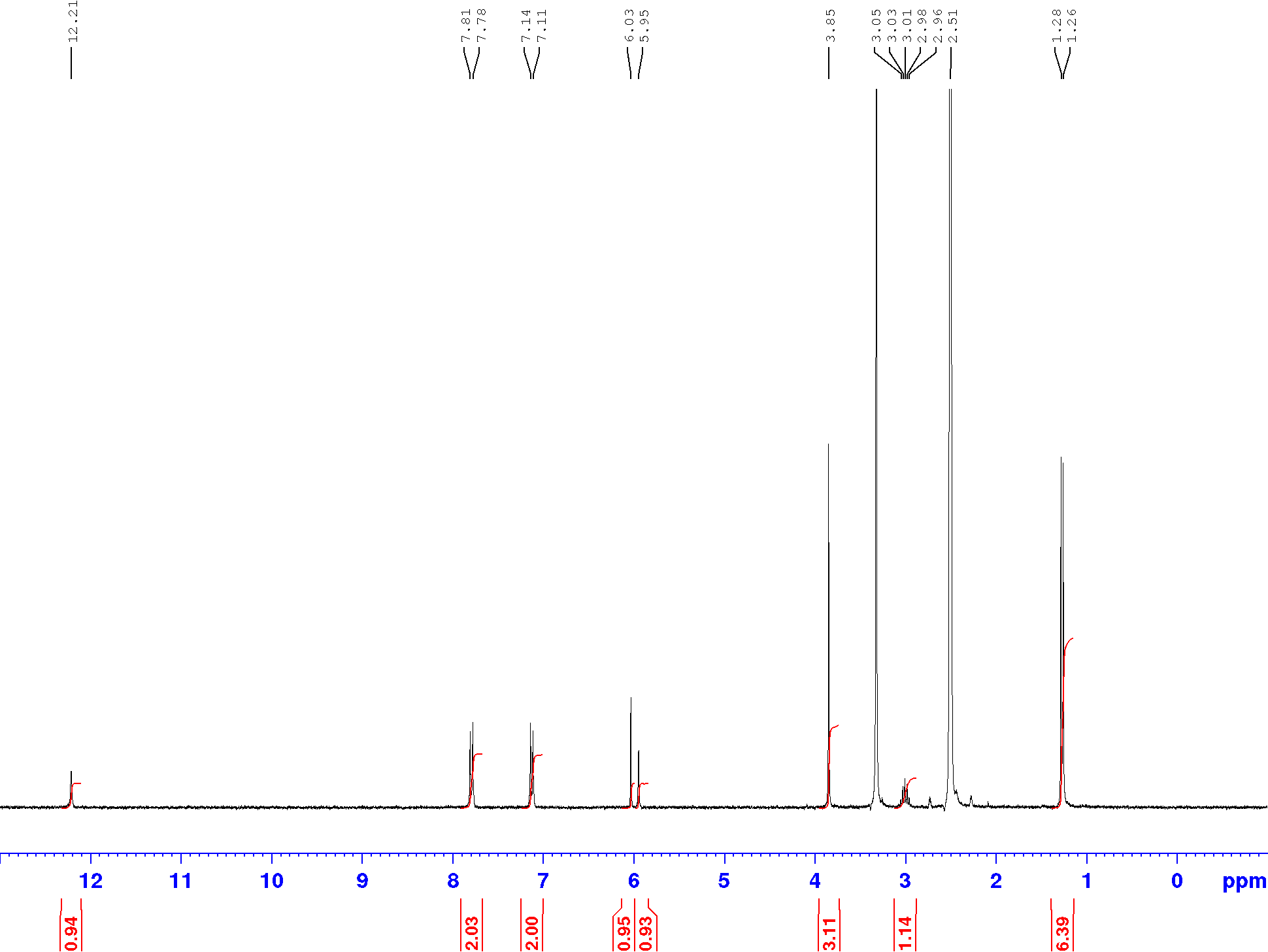

**Figure S29.** ^1^H NMR spectrum of 2-isopropyl-5-(4-methoxyphenyl)pyrazolo[1,5-a]pyrimidin-7(4H)-one (**3**)

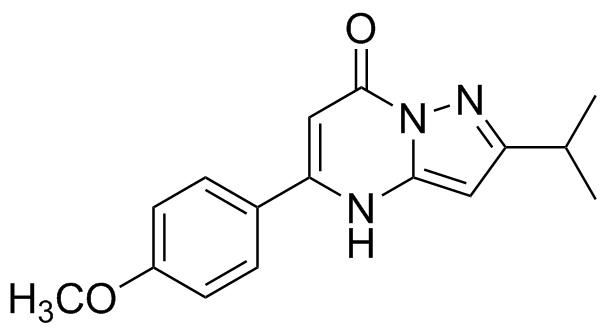

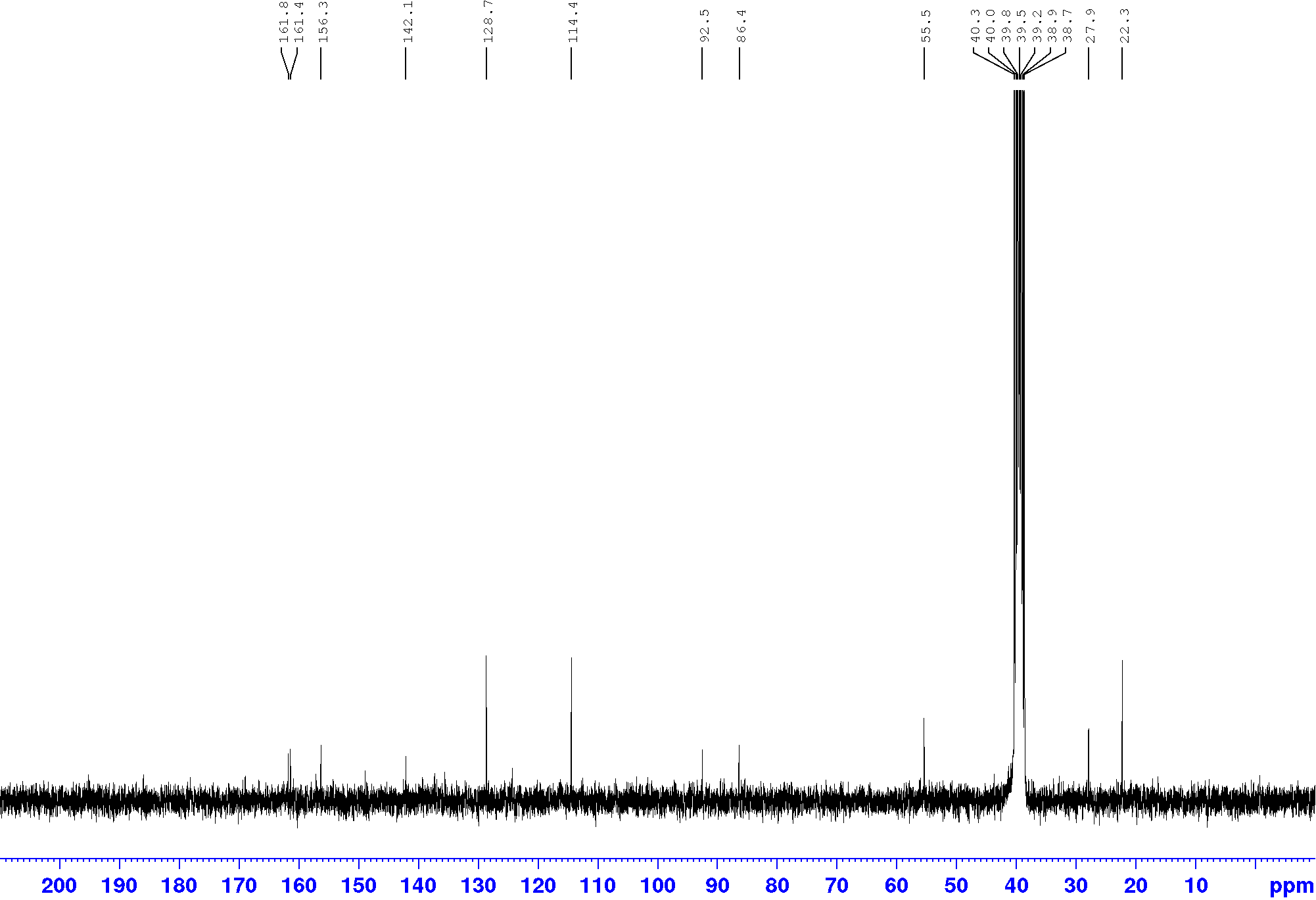

**Figure S30.** ^13^C NMR spectrum 2-isopropyl-5-(4-methoxyphenyl)pyrazolo[1,5-a]pyrimidin-7(4H)-one (**3**)

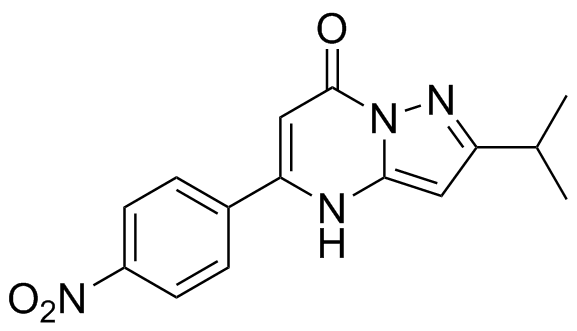

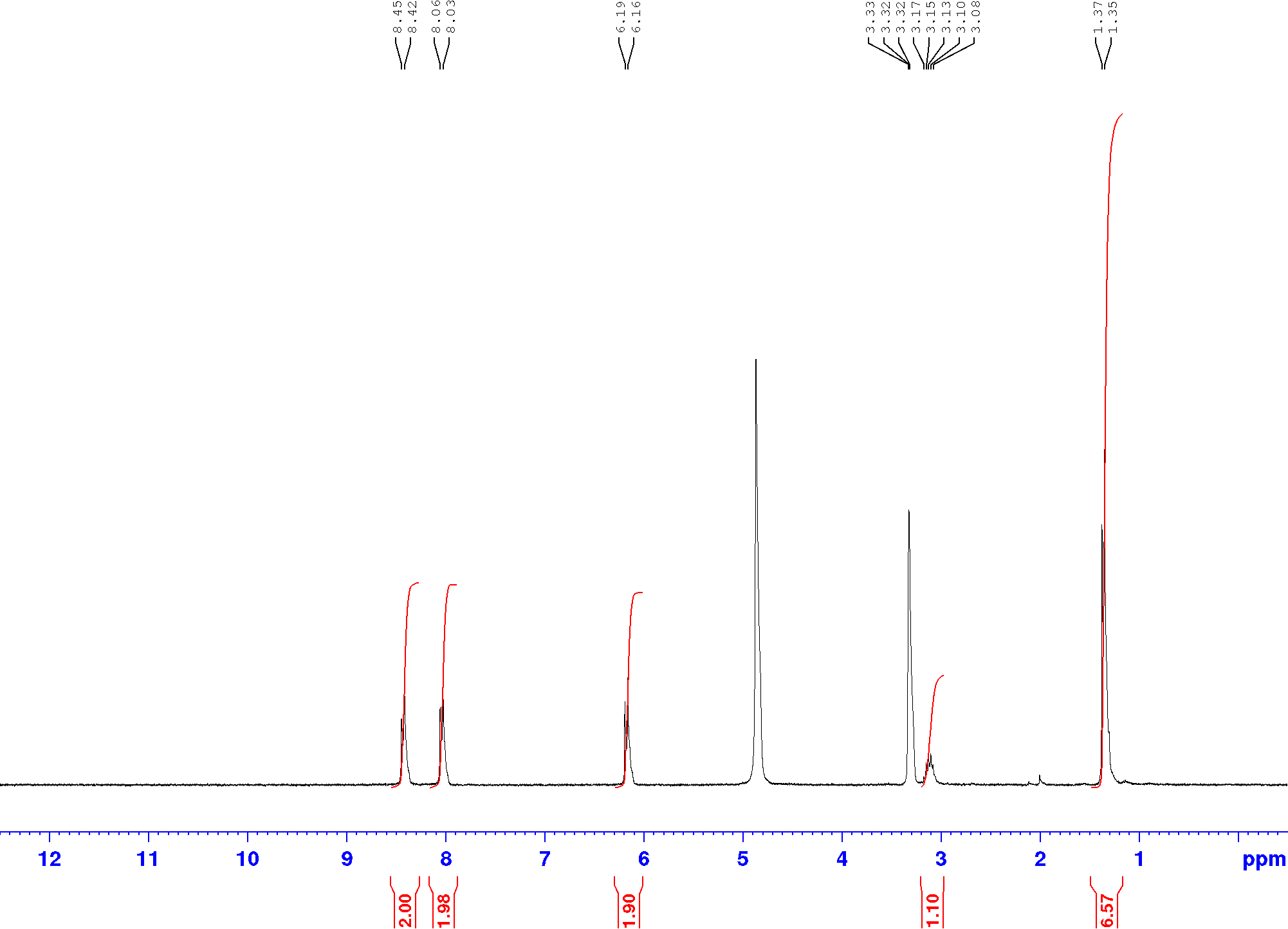

**Figure S31.** ^1^H NMR spectrum of give 2-isopropyl-5-(4-nitrophenyl)pyrazolo[1,5-a]pyrimidin-7(4H)-one (**1**)

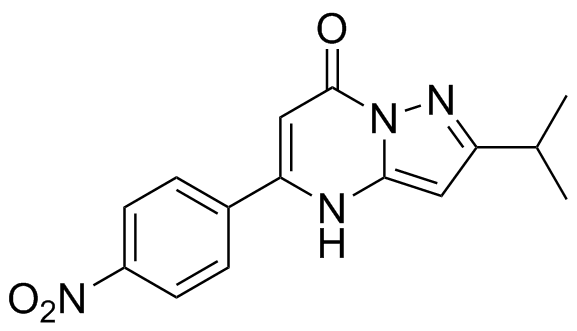

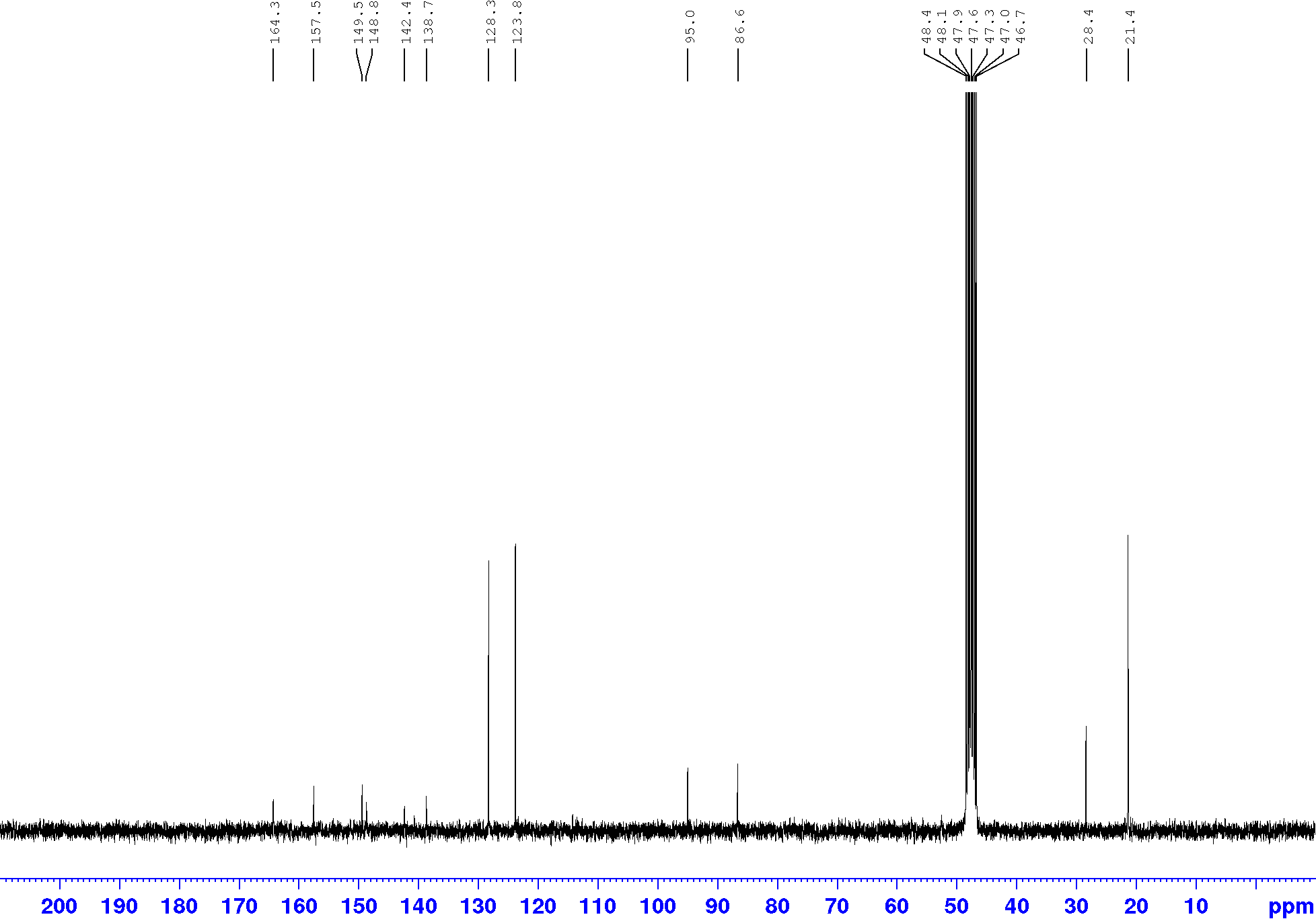

**Figure S32.** ^13^C NMR spectrum of give 2-isopropyl-5-(4-nitrophenyl)pyrazolo[1,5-a]pyrimidin-7(4H)-one (**1**)

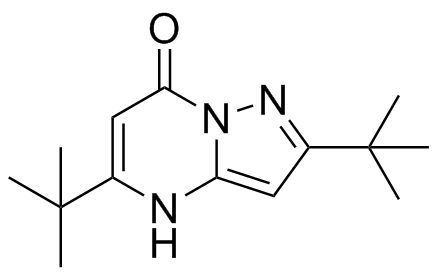

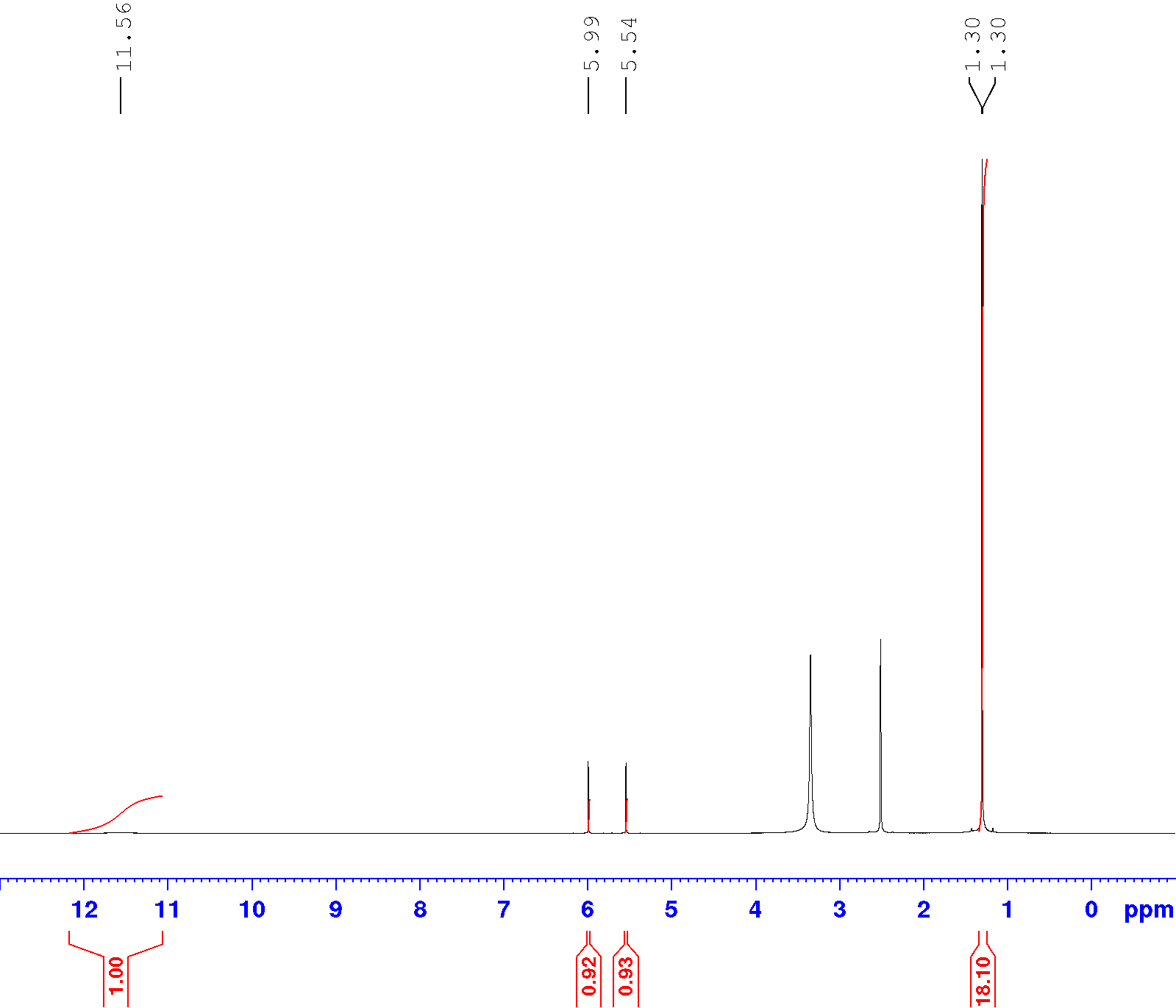

**Figure S33.** ^1^H NMR spectrum of 2,5-di-tert-butylpyrazolo[1,5-a]pyrimidin-7(4H)-one (**9**)

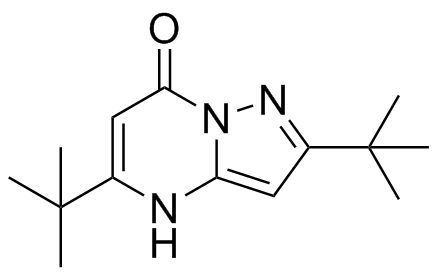

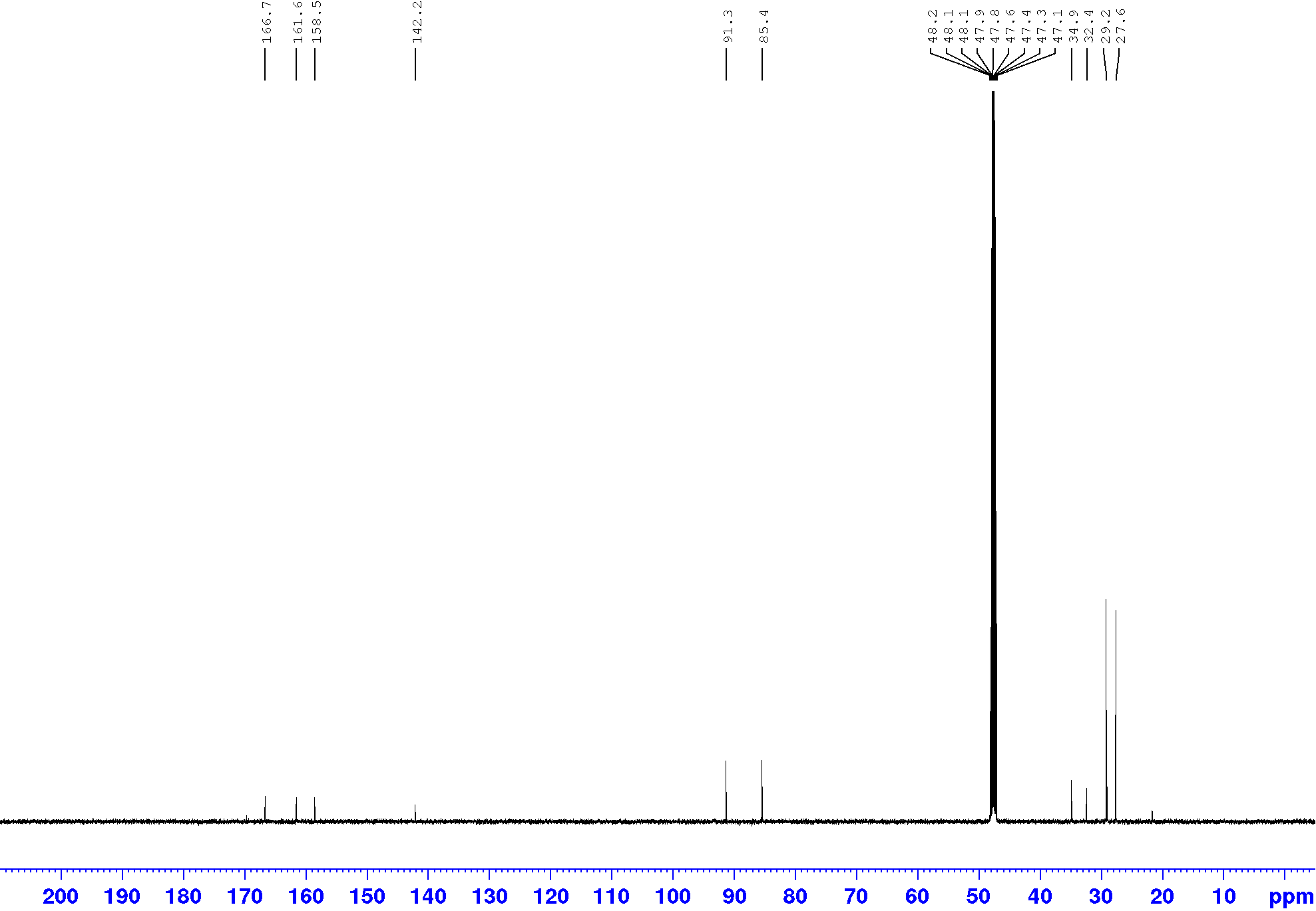

**Figure S34.** ^13^C NMR spectrum of 2,5-di-tert-butylpyrazolo[1,5-a]pyrimidin-7(4H)-one (**9**)

**1.5 References**

1. Kelada, M., Walsh, J. M. D., Devine, R. W., McArdle, P., Stephens, J. C. *Beilstein J. Org. Chem.* 2018, **14**, 1222–1228.

2. Z. T. Fomum, S. R. Landor, P. D. Landor, and G. W. P. Mpango, *J. Chem. Soc. Perkin Trans. 1* 1981, 2997–3001.

3. 7. S. Gogoi, K. Shekarrao, P. P. Kaishap, S. Gogoi and R. C. Boruah, *Tetrahedron Lett.* 2014, **55**, 5251–5255.

4. 9. K. Senga, T. Novinson, H. R. Wilson and R. K. Robins, *J. Med. Chem.* 1981, **24**, 610–613.

5. 11. S. J. Tantry, V. Shinde, G. Balakrishnan et al., *Med. Chem. Commun.* 2016, **7**, 1022–1032.

6. 10. N. L. Nam, I. I. Grandberg and V. I. Sorokin, *Chem. Heterocycl. Compd.* 2003, **39**, 1210–1212.

**2. Biology Experimental**
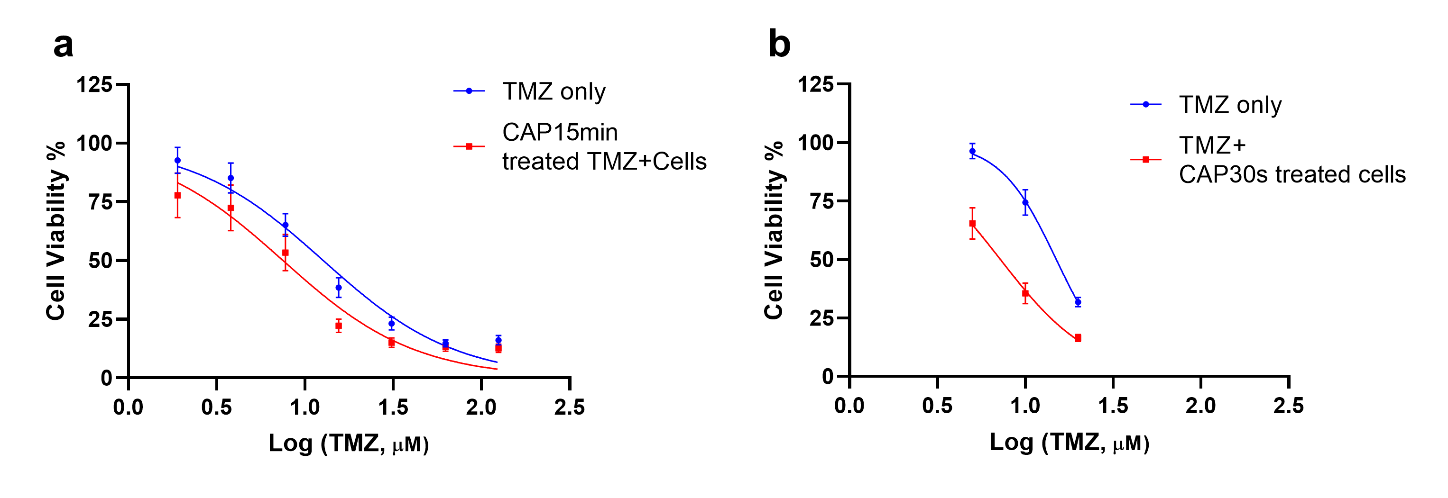

**Figure S35. Combination treatment of CAP and TMZ.** (a) Stock concentrations of TMZ (50mM in DMSO) were directly exposed CAP at 75kV for 15 minutes. Immediately after CAP treatment, working stocks of TMZ were made up in full media and added to untreated cells. As a negative control, cells were also treated with the same concentrations of TMZ that were not exposed to CAP. No deleterious effects were observed from vehicle DMSO control. (b) Cells were seeded and allowed adhere overnight, medium was then removed from each well and the cells were exposed to CAP for 30 secs. Following CAP treatment, cells were treated with a low medium and high dose of either TMZ (0, 5, 10, 20µM). Cells were analysed after 6 days by Alamar blue. No deleterious effects were observed from vehicle DMSO control.

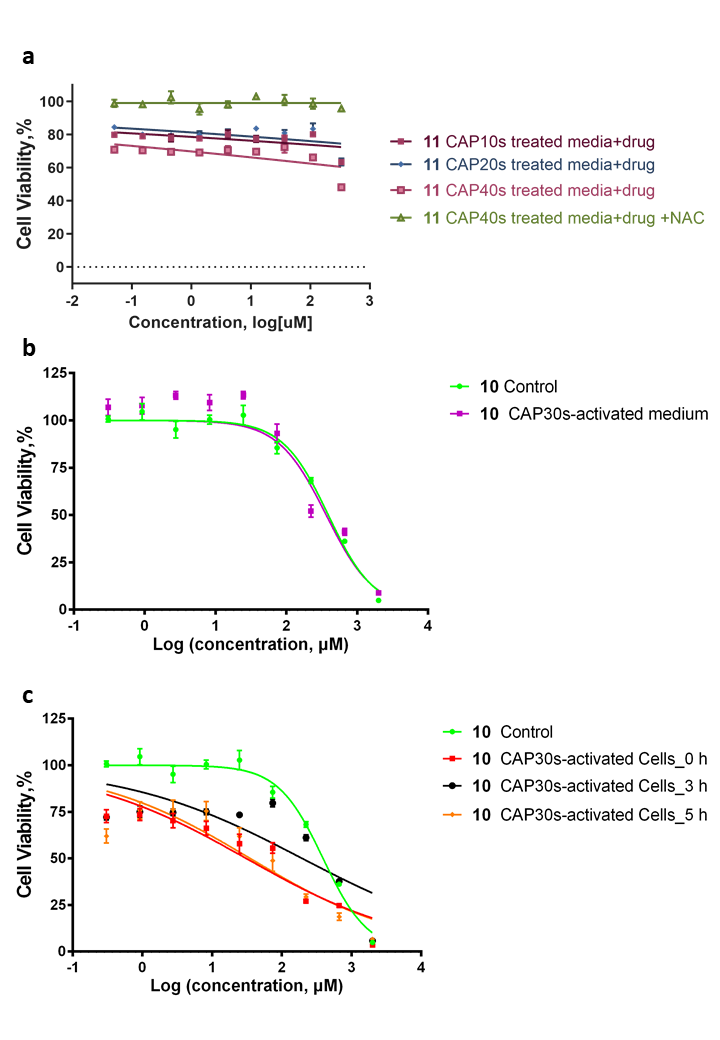
**Figure S36. Determine the synergistic cytotoxicity between CAP treatment and pro-drug 10.** (a) **11** were prepared in the culture medium, treated with CAP for 0-40 s with or without NAC and then incubated with U-251 MG cells in 96-well plates for 48 hours before cell viability was assessed. (b) The culture medium was treated with CAP for 30 s and storage overnight. The U-251 MG cells were then incubated with pro-drug **10** in overnight storage CAP-activated medium for 48 hours. (c) U-251 MG cells were treated with CAP and then incubated in the same CAP-treated culture medium for 0-5 h, and then incubated with fresh medium containing pro-drug **10** for 48 h before cell viability assay.
